# Leucine-rich repeat kinase 2 impairs the release sites of Parkinson’s disease vulnerable dopamine axons

**DOI:** 10.1101/2025.08.28.672006

**Authors:** Chuyu Chen, Qianzi He, Giulia Tombesi, Eve Napier, Matthew Jaconelli, Oscar Andrés Moreno-Ramos, Hannah Serio, Yahaira Naaldijk, Vanessa Promes, Amanda Schneeweis, Kaitlyn Quinn, Christopher Nasios, Elisa Greggio, Yevgenia Kozorovitskiy, Daniel Arango, Amir R. Khan, Dario R. Alessi, Daniel A. Dombeck, Sabine Hilfiker, Rajeshwar Awatramani, Loukia Parisiadou

## Abstract

The end-stage pathology of Parkinson’s disease (PD) involves the loss of dopamine-producing neurons in the substantia nigra pars compacta (SNc). However, synaptic deregulation of these neurons begins much earlier. Understanding the mechanisms behind synaptic deficits is crucial for early therapeutic intervention, yet these remain largely unknown. In the SNc, different dopamine neuron subtypes show varying susceptibility patterns to PD, complicating our understanding. This study uses intersectional genetic mouse models to uncover synaptic perturbations in vulnerable dopamine neurons, focusing on the LRRK2 kinase, a protein closely linked to PD. Through a combination of immunofluorescence and advanced proximity labeling methods, we found higher LRRK2 expression in the most vulnerable dopamine neuron subclusters. High-resolution imaging revealed that pathogenic LRRK2 disrupts release sites in vulnerable dopamine axons, leading to decreased in vivo spontaneous and evoked striatal dopamine release in mice with LRRK2 mutations. Proteomic and biochemical analyses indicate that mutant LRRK2 increases the phosphorylation of RAB3 proteins, reducing their interactions with RIM1and RIM2 effector proteins and impacting their synaptic functions. Overall, this research highlights the cell-autonomous dysfunctions caused by mutant LRRK2 in neurons, which are primarily affected by the disease. It also provides a framework for future therapeutic strategies for early nigrostriatal synaptic deficits in PD.

## Introduction

Parkinson’s disease (PD) is marked by the progressive degeneration of dopamine neurons in the substantia nigra pars compacta (SNc).^1,2^ These neurons are particularly vulnerable due to their unique intrinsic properties, such as large numbers of synaptic sites, massive arborization, and high bioenergetic burden.^3^ Notably, in PD, there is a selective loss of dopamine neurons within the SNc, with greater ventral than dorsal loss.^4–6^ This suggests subtype-specific dysfunction among SNc dopamine neurons. The difficulty in defining and isolating these subtypes has hindered the direct study of vulnerable neurons. To address this, single-cell profiling studies have identified a ventrally biased *Sox6+*/Aldehyde Dehydrogenase 1A1 (*Aldh1a1+*) dopamine neuron subcluster in humans and mice.^7–9^ The location of the cell bodies (ventral SNc) and axonal projections (dorsal striatum) of Aldh1a1+ cells corresponds to the patterns of neuron and projection loss observed in PD.^10–12^ Additionally, recent research identified Annexin 1 (*Anxa1*) as a subset of *Aldh1a1+* dopamine neurons occupying the ventral-most SNc. These *Anxa1+* neurons project densely to the most dorsal striatum and serve as markers of vulnerability in PD mouse models ^13–15^ and in the brains of human PD patients.^16^ This molecular classification has enabled the development of new intersectional genetic models ^12,13,17^ to study these otherwise intermingled SNc dopamine neuron populations, ultimately enhancing our understanding of PD mechanisms.

Deficits in dopamine release and synaptic dysfunction are well-documented hallmarks of prodromal and early PD. Neuropathological comparison of dopaminergic axon versus cell body loss in PD patients suggests that axons may be the initial site of dysfunction and degeneration.^18–20^ Impaired nigrostriatal dopamine transmission often precedes noticeable dopamine neuron loss—a convergent feature in human and numerous preclinical rodent models of PD (comprehensively reviewed in^21^). This suggests that synaptic dysregulation in dopamine neurons is one of the earliest manifestations in the context of diverse PD factors. Additionally, the presynaptic region is a known site of action for gene products associated with PD (e.g., auxilin, synaptojanin, α-synuclein).^22–25^ Despite these observations, our understanding of the molecular and cellular mechanisms behind early dopaminergic synaptic impairments remains limited.

Mutations in the *LRRK2* gene are among the most common genetic causes of PD, accounting for up to 30% in certain ethnic groups.^26–28^ The most prevalent pathogenic point mutation, G2019S, leads to a hyperactive LRRK2 kinase. Increased kinase activity is also observed in idiopathic PD,^29^ suggesting that aberrant LRRK2 kinase activity is a broad pathophysiological mechanism for this disorder. Currently, the clinical development of LRRK2 kinase inhibitors is underway, highlighting their potential as a viable therapeutic option for PD.^30,31^ However, despite its clinical relevance, the mechanism by which increased LRRK2 kinase leads to cellular dysfunction remains a critical knowledge gap. Several rodent models were developed to investigate the dysfunctions caused by LRRK2 mutations.^32,33^ While these models typically do not show loss of dopamine neurons^34^ in the absence of secondary hits, they do exhibit deficits in striatal dopamine transmission,^35–38^ suggesting that mutant LRRK2 affects the nigrostriatal synapse. LRRK2 protein is thought to be expressed in dopaminergic neurons in the SNc^39^, and deficits in striatal dopamine signaling are also observed in mouse models that express the pathogenic LRRK2 variant specifically within dopamine neurons.^40,41^ This indicates a cell-autonomous deregulation of dopamine release in these neurons. However, research on LRRK2 function in dopamine neurons is limited, and to date, no studies have examined the effects of LRRK2 mutations on the dopamine neuron subpopulations most vulnerable to disease.

Previous research on non-neuronal cells identified RAB proteins as robust downstream targets of LRRK2 kinase activity. The phosphorylation of RAB proteins by LRRK2 disrupts their normal function in intracellular membrane trafficking, resulting in cellular dysfunction.^42,43^ Due to various technical challenges, it remains unclear whether the aberrant phosphorylation of RAB proteins by LRRK2 occurs specifically in the most vulnerable dopamine neurons and how this relates to dopamine release deficits.

Using a combination of advanced intersectional genetic strategies, cell-type-specific proteomics methods, sophisticated biochemical approaches, super-resolution microscopy, and a platform to access in vivo dopamine release, we describe a novel pathophysiological role for LRRK2 in the synapses of the most vulnerable ventral tier SNc neurons. Our study provides high-resolution insights into the mechanisms underlying functional disruptions in dopamine signaling in a well-established preclinical LRRK2 model of early PD. Our findings enhance understanding of LRRK2 dysfunction and highlight biologically and pathologically relevant cellular changes in the neurons primarily affected in the disease.

## Results

### LRRK2 is expressed in dopamine neurons and is enriched in the vulnerable subpopulation of neurons

Single-cell profiling studies and research on both postmortem PD tissues and preclinical mouse models demonstrated that Sox6/Aldh1a1 markers molecularly define the vulnerable ventral tier dopamine neurons,^11,12^ while Calbindin1 (Calb1) marks the resilient dorsal tier neurons (Figure 1A).^1,44^ These distinct dopamine neuronal subtypes differ in their molecular profile, and they also exhibit unique projection patterns and functional responses.^10,13,17^ Therefore, studying these neurons in a subtype-specific manner can yield important insights into cell-intrinsic mechanisms of vulnerability in PD. We first addressed whether LRRK2 is expressed in dopamine neurons, as this is a prerequisite for any cell-autonomous effects. We performed immunolabeling on mouse brain sections and confirmed LRRK2 expression in SNc dopamine neurons (Figure 1B). No LRRK2 signal was observed by Western blot (WB) or immunolabeling in *Lrrk2*^KO^ tissue, confirming the specificity of the antibodies used (Supplementary Figure 1A, B).

**Figure 1:**
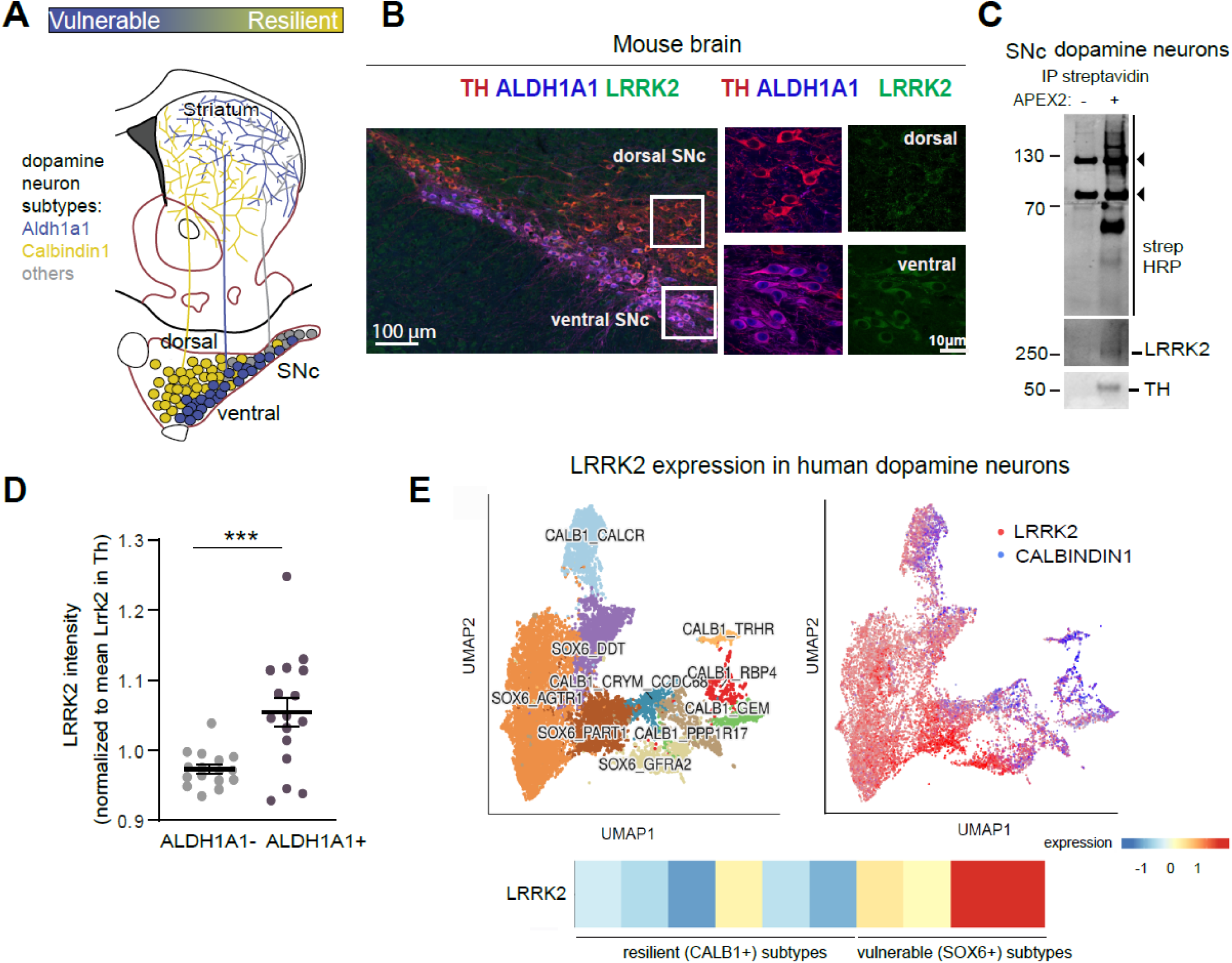
LRRK2 is expressed in SNc dopamine neurons and enriched in vulnerable subpopulations. **A.** Schematic illustrating the localization of selective dopamine neuron subtypes in the mouse SNc and their projection patterns in the striatum. Aldh1a1 defines the vulnerable dopamine neuron populations in PD that are located more ventrally. The resilient dopamine neuron subtypes are characterized by calbindin 1 (*Calb1*+) and are located more dorsally.^11–13^ **B.** *Left,* Representative image showing TH, ALDH1A1, and LRRK2 immunostaining in the SNc of wildtype mice. *Right,* High magnification images of the boxed regions in the dorsal (ALDH1A1-) and ventral (ALDH1A1+) SNc areas. **C**. WB analysis of streptavidin pulldowns from SNc dissections from DAT^cre^ mice crossed with APEX2 EGFP mice (detailed in Supplementary Figure 1) probed for streptavidin, LRRK2, and TH. Arrowheads to the right for the strong bands at ∼75 and ∼150 kDa indicate endogenous biotinylated carboxylase proteins. Each lane contains an equal amount of proteins eluted from the streptavidin beads. **D.** Quantification of LRRK2 intensity from B in Aldh1a1-(less vulnerable) and Aldh1a1+ (vulnerable) dopamine neurons. Each dot represents the average LRRK2 intensity of 60-170 cells/section. n=16 sections/3 mice. Data are represented as mean ± SEM. Asterisks show statistical significance for the unpaired t-test. \*\*\**p*< 0.001. **E.** *Left,* Clusters of dopamine neuron subtypes based on the human snRNA sequencing dataset from^49^ colored by subtype and plotted using Shinny Cell.^93^ *Right,* Feature plots from the same dataset showing limited coexpression between *LRRK2* and *CALB1+*. *Bottom,* Heatmap showing expression of *LRRK2* with apparent enrichment in vulnerable (*SOX6+*) but not resilient (*CALB1+*) subclusters.

As an orthogonal approach, we developed an independent methodology that employs proximity labeling of genetically labeled neuronal cell types in the mouse brain (Supplementary Figure 1C), focusing on APEX2. APEX2 is a soy peroxidase that rapidly biotinylates proximal proteins.^45^ We achieved specific APEX2 expression in dopamine neurons by crossing the Cre-dependent APEX2 EGFP reporter mouse line^46^ with DAT^Cre^ mice. We then prepared acute brain slices from DATCre; APEX2 EGFP and induced rapid biotinylation following optimized published protocols for the mouse brain.^46–48^ Immunostaining with EGFP and tyrosine hydroxylase (TH) confirmed APEX2 expression in dopamine neurons and their axons, as shown in sections from the SNc and striatum, respectively (Supplementary Figure 1D). Following tissue lysis, we purified biotinylated dopamine neuron proteins and subjected them to mass spectrometry analysis (MS). By referencing a publicly available APEX2 proteome database of dopamine neurons, our analysis confirmed enrichment of dopamine neuron proteins (Supplementary Figure 1E)^48^. We also conducted streptavidin pulldowns to isolate biotinylated proteins from either dopamine neurons or their axons in the striatum, followed by WB. Slices from control DAT^Cre^ and APEX2 negative mice were included to assess non-specific binding. Streptavidin–horseradish peroxidase (HRP) blotting in SNc and striatal lysates showed broad biotinylation in DAT^Cre^; APEX2 EGFP samples but not in control samples. TH immunoblotting further indicated the specificity of the labeling (Supplementary Figure 1F, G). Using this refined methodology, we independently confirmed the presence of LRRK2 in SNc dopamine neurons and detected a faint LRRK2 signal in dopamine axons (Figure 1C, Supplementary Figures 1H and 1I).

Notably, quantification of our immunolabeling experiments indicated that LRRK2 protein expression was higher in the most vulnerable dopamine neuron populations in the mouse brain, specifically those of the ventral tier SNc, marked by Aldh1a1 (Figure 1D). Additionally, single-nucleus RNA sequencing datasets from humans show that the *LRRK2* gene is enriched in specific midbrain dopamine neuron subtypes marked by *SOX6* (Figure 1E). These particularSOX6+ populations, including the ALDH1A1*+* subcluster, are known to selectively degenerate in human PD.^49^

Altogether, our findings indicate that the LRRK2 protein is enriched in the most vulnerable populations of dopamine neurons in both mice and humans, supporting the idea of a cell-autonomous LRRK2 dysfunction in these populations. neurons in PD.

### LRRK2 phosphorylates RAB3 isoforms in vivo

We and others have previously reported that knock-in mice expressing the pathogenic hyperactive LRRK2^G2019S^ mutation exhibit deficits in striatal dopamine release.^35,37^ To investigate the underlying molecular mechanisms, we conducted quantitative phosphoproteomic Liquid Chromatography-Mass Spectrometry (LC-MS) studies on striatal synaptosomes from LRRK2^G2019S^ mice. The mice were treated with either vehicle control or the potent and specific LRRK2 kinase inhibitor MLi-2.^50^ We focused on this comparison because pharmacological inhibition of the already hyperactive LRRK2^G2019S^ kinase increases the likelihood of identifying in vivo changes in phosphoregulation. First, we confirmed effective target engagement under our experimental conditions by assessing RAB12 phosphorylation at Ser106, a well-established readout of LRRK2 kinase activity. ^51^ In extracts from LRRK2^G2019S^ mice, RAB12 phosphorylation was higher compared to wildtype, but it returned to wildtype levels upon MLi-2 treatment (Supplementary Figure 2A, B). We used our previously described protocols for subcellular fractionation to isolate striatal synaptosomes,^52^ which were subsequently subjected to MS for both total and phosphoproteomic analyses (Figure 2A). When comparing vehicle-to MLi-2 treated LRRK2^G2019S^ mice (unadjusted p value < 0.05 and |log2FC|>0.58), we identified 289 significantly altered phosphopeptides (Figure 2B). To further categorize these changes, we performed pathway enrichment analyses using gene ontology (GO) to annotate the differentially expressed phosphoproteins with at least one altered phosphopeptide. The top 15 enriched pathways showcased key synaptic biological processes, including signal release from the synapse and neurotransmitter secretion (Figure 2C), as well as presynaptic membrane and axonal localizations (Supplementary Figure 2C). These findings suggest that increased LRRK2 kinase activity causes presynaptic alterations in phosphoproteins, which may contribute to reduced nigrostriatal dopamine release in LRRK2^G2019S^ mice. In contrast, total protein analysis from the same samples revealed only 12 differentially expressed proteins (Supplementary Figure 2D).

**Figure 2:**
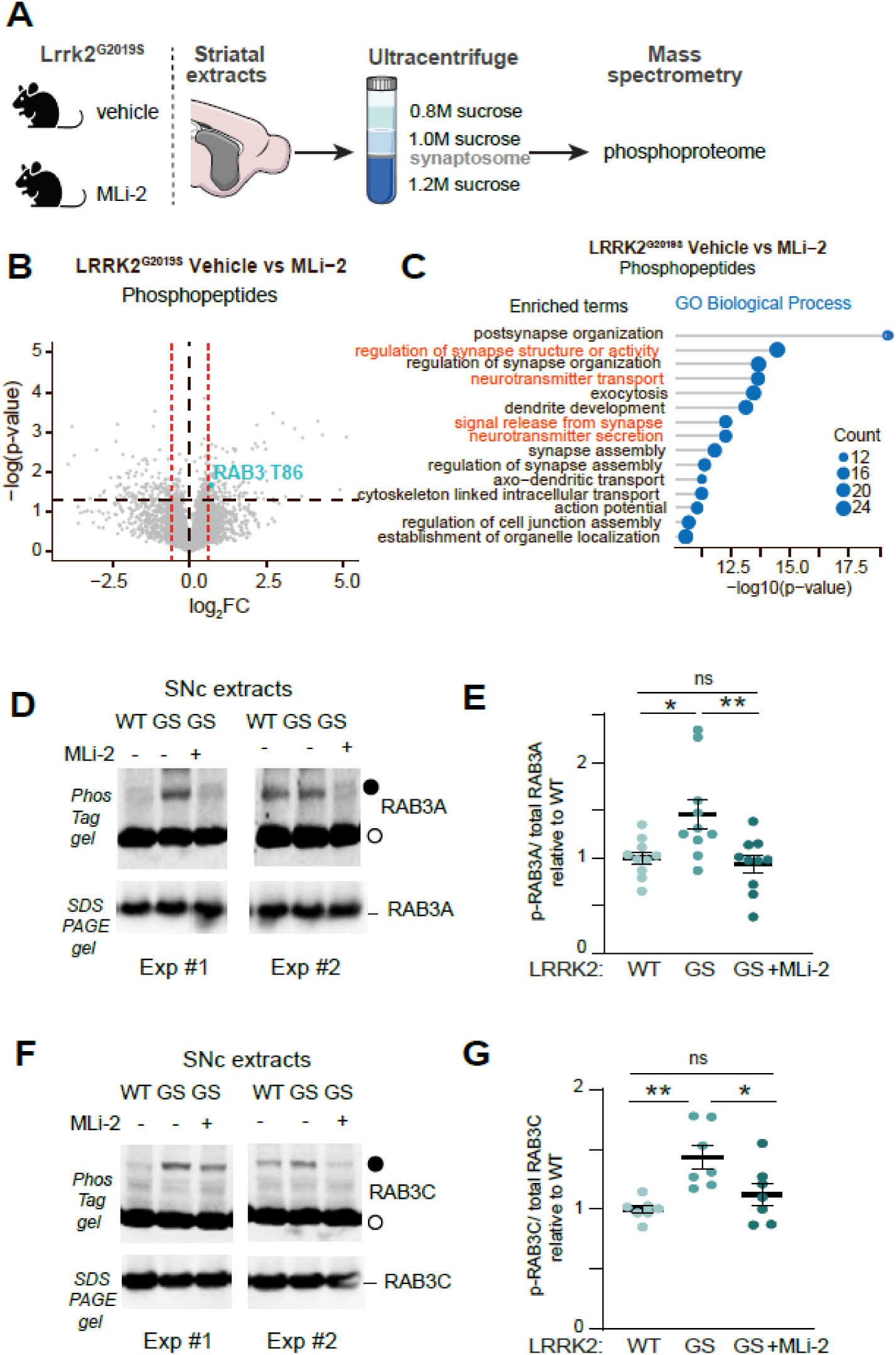
LRRK2 phosphorylates RAB3 proteins in vivo. **A.** Experimental workflow of LC/MS-MS analysis in striatal synaptosomes isolated by subcellular fractionation from 3 technical replicates, with each technical replicate performed using 3 LRRK2^G2019S^ mice treated with either MLi-2 or the corresponding vehicle for 2 hours. Treatment conditions: Vehicle for MLi-2, 40% 2-hydroxypropyl-β-cyclodextrin; MLi-2, 10_mg/kg. This treatment reduces LRRK2 kinase activity in vivo (see Supplementary Figure 2). **B.** Volcano plot comparing the significantly altered phosphopeptides between vehicle and MLi-2-treated LRRK2^G2019S^ mice (p≤_0.05 by multiple unpaired t-tests, |Log_2_FC|> 0.58). **C**. Gene Ontology analysis of proteins with at least one differentially regulated phosphopeptide in striatal synaptosomes from vehicle vs MLi2-treated LRRK2^G2019S^ mice. The top 15 pathways significantly enriched in the biological process term analysis. All enriched pathways have been uploaded to Zenodo (link in the key resources table) and presented in Supplementary File 1. **D.** Representative PhosTag and SDS PAGE gels (upper and lower panels, respectively) from two independent experiments of LRRK2^WT^ and LRRK2^G2019S^ mice (indicated WT, GS, respectively) with or without MLi-2 treatment (10 mg/kg, 2 hours). SNc extracts from these mice were probed for RAB3A (closed/open circles = phosphorylated (p)-RAB3A/unphosphorylated RAB3A species, respectively). **E.** p-RAB3A/total (phosphorylated +unphosphorylated) RAB3A quantification. n=10 mice/treatment. All samples were normalized to the average signal of all WT samples in the same blot. **F.** Same as D, but probing for RAB3C. **G.** Same as E, but for RAB3C. n=7 mice/treatment. In **E** and **G,** data represent mean±SEM. Asterisks indicate statistical significance, as determined by Šídák’s multiple comparisons test following a two-way ANOVA. E: Treatment factor F(1,27)=11.33 p=0.0023, Genotype factor F(1,27)=8.593 p=0.0068. G, Treatment factor F(1,18)=7.398 p=0.0140, Genotype factor F(1,18)=14.76 p=0.0012. *p<0.05, **p<0.01.

One notable finding from our phosphoproteome analysis was the decreased phosphorylation at the T86 site of the RAB3 GTPase in LRRK2^G2019S^ mice treated with MLi-2, compared to the vehicle group (Figure 2B). Given that RAB3 proteins were previously identified as targets of the LRRK2 kinase in cell lines,^42,43^ our phosphoproteomic results suggest that LRRK2 may phosphorylate RAB3 proteins in the mouse brain. There are four RAB3 proteins—RAB3A, RAB3B, RAB3C, and RAB3D—that are involved in the synaptic vesicle cycle at the presynaptic site.^53,54^ To examine the relative expression of these four proteins in dopamine neurons, we first analyzed a dataset from single-cell RNA sequencing studies in mice.^9^ This analysis indicated that RAB3A and RAB3C had the highest expression levels in dopamine neurons (Supplementary Figure 2E). In addition, the examination of a human dopamine neuron single-nucleus dataset revealed that RAB3B is predominantly found in resilient dopamine neuron subclusters,^49^ rather than in the vulnerable ones, which show higher LRRK2 expression (Supplementary Figure 2F). Taken together, these findings suggest that the availability of LRRK2 kinase activity targets differs between vulnerable and resilient dopamine neuron subpopulations.

To evaluate the presence of RAB3A and RAB3C proteins in vulnerable dopamine axons, we labeled Anxa1+ ventral tier dopamine neurons through Cre-dependent viral delivery of APEX2 in the SNc, followed by MS. ^47^ This proteomics approach confirmed high levels of RAB3A and RAB3C in these vulnerable dopamine axons (Supplementary Figures 2G and H). Next, employing our APEX2-based biochemical approach, we further validated the presence of RAB3A and RAB3C in dopamine neurons and their axons. We observed their apparent presence in dopamine axons (striatum), consistent with their presynaptic roles (Supplementary Figure 2I). Additionally, the higher levels of synaptophysin detected in pulldowns from striatal extracts, compared with those from SNc extracts from the same mouse, corroborate the compartment specificity of our method.

Considering these expression patterns, we focused on RAB3A and RAB3C proteins, examining their phosphorylation in mouse brain SNc extracts and how this is influenced by LRKK2 kinase activity. Due to the lack of specific antibodies for phosphorylated RAB3, we used phosTag gels to separate phosphorylated RABs from their non-phosphorylated counterparts by SDS-PAGE^42,55^ and then we blotted the membranes with RAB3 isoform-specific antibodies (Figure 2D-G) Given the relative homology of RAB3 isoforms, to determine the isoform specificity of the antibodies we analyzed extracts from cells transiently transfected with different RAB3 isoforms (Supplementary Figure 2J). The slower migrating RAB3A band (corresponding to phosphorylated species, closed circle, Figure 2D) was increased in the SNc of LRRK2^G2019S^ mice as compared to littermate controls. This increase returned to wild-type levels when the LRRK2^G2019S^ mice were treated with MLi-2 (Figure 2D, E). Similarly, an increase in phosphorylated RAB3C protein was observed in the mutant LRRK2 compared to the wild type, which was reversed upon MLi-2 treatment (Figure 2F, G).

Overall, our findings indicate that the phosphorylation of RAB3A and RAB3C is aberrantly increased in the SNc of LRRK2^G2019S^ mice, and this elevation depends on LRRK2 kinase activity.

### Phosphorylation of RAB3A decreases its binding to the effector proteins RIM1 and RIM2 from mouse brain

RAB3 proteins play an important role in coordinating presynaptic vesicle trafficking and release.^56^ They achieve this by binding to effector proteins such as RIM1 and RIM2.^54^ Prior studies conducted in vitro and in cells^42,43,57^ indicate that LRRK2-induced hyperphosphorylation of RAB proteins disrupts their interaction with regulatory and effector proteins, while at the same time promoting interactions with phospho-specific effectors of the RILP family through their conserved RH2 domains.

To investigate the interactome of phosphorylated (p-) versus unphosphorylated RAB3A in the mouse brain, we employed a biochemical strategy combined with MS (Figure 3A). We purified Flag-RAB3A and phosphorylated it at T86 by MST3 kinase, which phosphorylates RAB proteins at the same site as LRRK2 and is easier to purify in milligram amounts for biochemical studies.^58,59^ Phosphorylated Flag-RAB3A was purified using MonoQ chromatography, and stochiometric phosphorylation was confirmed by PhosTag gel analysis. The site of phosphorylation was established as Thr86 through MS experiments (Supplementary Figure 3A-B). Recombinant p-Flag-RAB3A or unmodified Flag-RAB3A were then used as ‘bait’ for an MS affinity purification experiment using homogenized whole brain extracts from mice that received MLi-2 via subcutaneous injection (30 mg/kg) 2 hours prior to termination (to induce dephosphorylation of endogenous RAB3 phosphorylation). Proteins that interacted with p-RAB3A or RAB3A were detected and quantified using a data-independent acquisition (DIA) approach. This analysis revealed a total of 3,480 proteins, with 69 proteins preferentially interacting with RAB3A compared to p-RAB3A, while another set of 16 proteins exhibited a higher interaction with p-RAB3A as compared to RAB3A (p-value<0.05, |log2FC|=1) (Figure 3D). These data can be further explored using the interactive CURTAIN volcano plot^60^ using the weblink provided in the figure legend. The RAB3A effector proteins RIM1 and RIM2 showed a marked preference for binding to unphosphorylated RAB3A (Figure 3D, E). This finding aligns with the observation that LRRK2-mediated phosphorylation of other RAB proteins, occurring within the effector-binding motif, induces steric clashes that hinder their interactions with respective effector proteins.^43^ RIM1 and RIM2 are essential components of the presynaptic active zone, which serves as the structural and functional platform for dopamine release (Figure 4A).^61^A precise arrangement of active zone proteins is critical for effective neurotransmission. Key components s, such as ERC1/2 (ELKs) and UNC13 (MUNC13), which are known to bind to RIM1 and RIM2, also exhibited differential binding to unphosphorylated RAB3A (Figure 3D). This likely reflects the decreased interaction between RIM1, RIM2, and p-RAB3A. Conversely, we found that p-RAB3A preferentially associated with RILPL1 and RILPL2 (Figure 3D,F), known phospho-specific effectors that recognize phosphorylated RABs and have been linked to the regulation of ciliogenesis in neuronal cells.^42,62,63^

**Figure 3:**
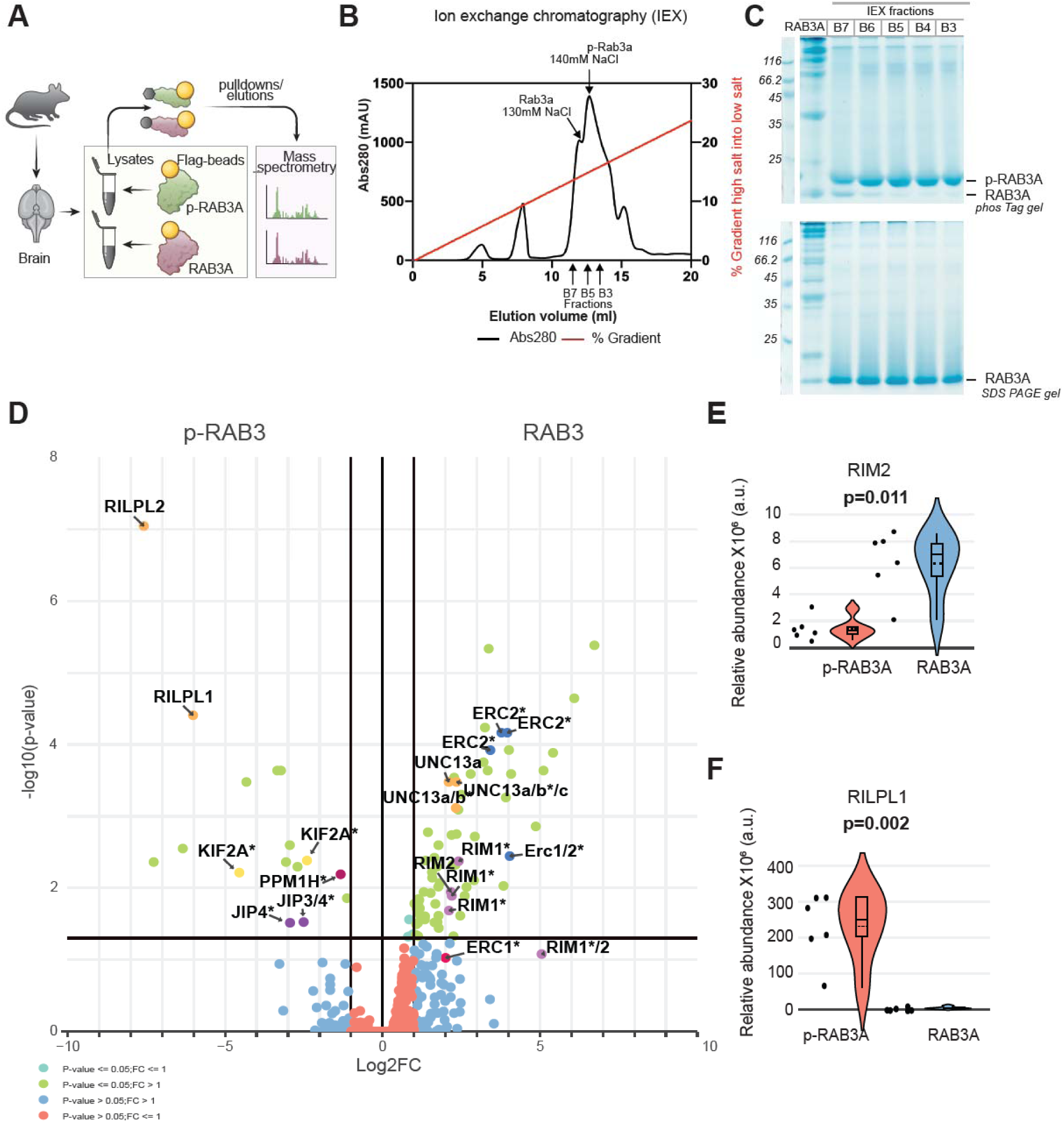
Differential interactome between p-RAB3A and unphosphorylated RAB3A in extracts from mouse brain. **A**. Experimental workflow. Purified Flag-tagged RAB3A or p-RAB3A were incubated with mouse brain lysates. The pulldown samples containing proteins that bound to either RAB3A or p-RAB3A were subjected to MS analysis (Orbitrap Astral). **B.** For purification, p-Flag-RAB3A was subjected to ion exchange chromatography on a MonoQ column and eluted with a 35% 1M NaCl gradient followed by gel filtration. **C.** Ion exchange chromatography fractions from p-RAB3A and RAB3A were resolved on either phos-Tag (upper panel) or SDS-PAGE gels (lower panel). **D**. Volcano plot of proteins with differential affinity for p-RAB3A as compared to RAB3A. The RAB3A effectors RIM1 and RIM2, along with other active zone proteins such as MUNC13 (UNC13) and ELKs (ERC1/2), preferentially interact with unphosphorylated RAB3A. In contrast, other proteins, such as RILPL1 and RILPL2, preferentially bind to p-RAB3A. In the volcano plot, proteins quantified by peptides matching the canonical sequence and a combination of their known isoforms are denoted by *, while a slash (/) indicates protein groups quantified through peptides shared between proteins. Different color codes indicate proteins of interest. For RAB3A: purple, RIM1, RIM2; dark blue, ERC; orange, UNC isoforms. For p-RAB3A, orange: RILPL1, RILPL2. Data were visualized using the interactive Curtain tool and can be accessed through the CURTAIN link: https://curtain.proteo.info/#/7eb667fc-a390-4484-8894-31e8ba78cbb2 **E, F.** Violin plots showing the relative protein abundances of RIM2 and RILPL1 (DIA-NN 1.9.2) through interactions between RAB3A (RIM2; p-value 0.011) and p-RAB3A (RILPL1; p-value 0.002). Univariate statistical testing conducted via paired Student’s t-test (Scipy version 1.11.1). N=6 replicates per condition.

**Figure 4:**
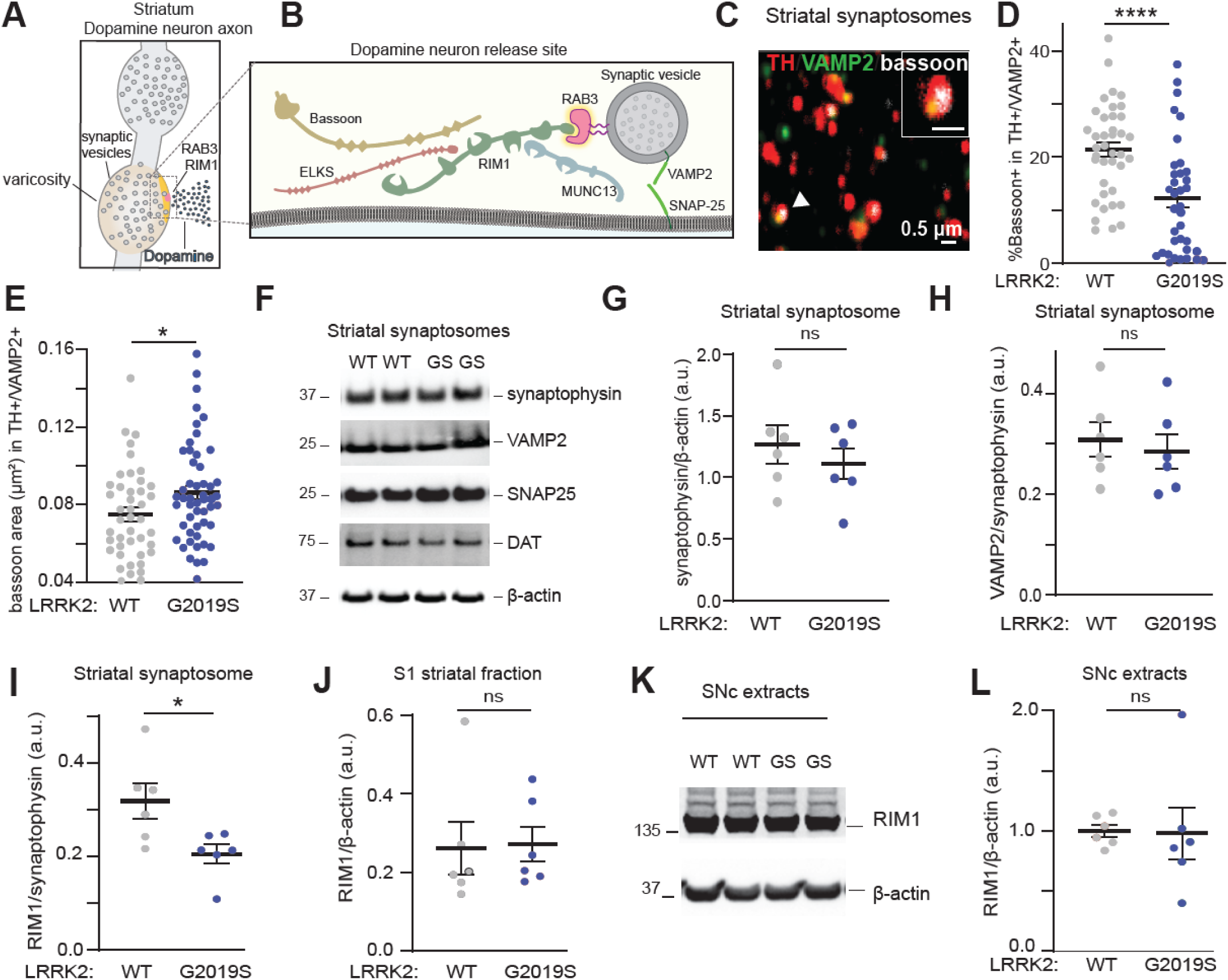
LRRK2 affects the composition of active zone sites in dopaminergic striatal synaptosomes. **A.** Schematic illustrating varicosities in dopamine axons, with active zone-like release site in orange. **B.** Dopaminergic axonal release site, with key molecular components as indicated**. C.** Confocal image of purified striatal synaptosomes stained with bassoon, VAMP2, and TH. The arrowhead indicates a TH+ striatal synaptosome (also shown in the inset), positive for both bassoon and VAMP2. Scale bar, 0.5μm **D..** Quantification of the percentage of TH+/VAMP2+ synaptosomes that contain bassoon from LRRK2^WT^ and LRRK2^G2019S^ mice. **E**. Quantification of area of bassoon staining in TH+/VAMP2+ synaptosomes. For data in D and E, each circle represents the average result of ∼1,000 synaptosomes scored per imaging area. N=10-14 areas/3 mice per genotype. Data are mean± SEM. Asterisks indicate statistical significance after unpaired t-tests. *p <0.01, **** p<0.0001. **F.** Purified striatal synaptosome fractions from LRRK2^WT^ and LRRK2^G2019S^ mice (indicated WT, GS, respectively) were subjected to WB analysis and probed for the antibodies shown on the right. **G**. Quantification of synaptophysin/β-actin ratio in the striatal synaptosome preparations shows no difference between LRRK2^WT^ and LRRK2^G2019S^ mice. Synaptophysin was used as a loading control for further comparisons across genotypes. **H**-**I** Quantification of indicated proteins in striatal synaptosome fractions from LRRK2^WT^ and LRRK2^G2019S^ mice (RIM1 representative blot is shown in Supplementary Figure 5F). **J.** Quantification of RIM1 in the S1 striatal fraction. **K.** WB analysis of SNc extracts from LRRK2^WT^ and LRRK2^G2019S^ mice using RIM1 and β-actin antibodies. **L.** Quantification of RIM1 signal in SNc extracts across different genotypes. Equal amounts of proteins were loaded in each lane, and duplicate lanes represent biological replicates. Data represent mean±SEM. Asterisk shows significance after the unpaired t-test. *p <0.01. Ns = no significant difference. N=6 mice/genotype.

These data indicate that increased phosphorylation of RAB3A by LRRK2 disrupts its interaction with its effectors RIM1 and RIM2 in brain extracts. This suggests a potential regulatory mechanism by which RAB3 phosphorylation may impact dopamine release in mutant LRRK2 mice.

### LRRK2 alters the composition and organization of active zone release sites in dopaminergic striatal synaptosomes

Dopaminergic neuronal release sites have unique characteristics, with only about 25–30% of axonal varicosities containing active zones that facilitate transmitter release.^64–66^ RIM1 and RIM2 proteins are important scaffolding molecules that organize these release sites (Figure 4B). A dopamine neuron-specific knockout of RIM1 and RIM2 results in structural disorganization of active zones and almost completely abolishes evoked dopamine release in the nigrostriatal pathway.^64,67^

Our phosphoproteomic analysis revealed that several pathways involved in the organization of presynaptic release sites are altered in response to changes in LRRK2 kinase activity (Figure 2). Furthermore, we observed a decrease in the interaction between phosphorylated RAB3A and various active zone proteins in the mouse brain (Figure 3). Therefore, we next examined whether alterations in the number and/or organization of active zones may form the cellular basis for the observed deficits in dopamine release in LRRK2^G2019S^ mice. To do so, we prepared striatal synaptosomes from LRRK2^WT^ and LRRK2^G2019S^ mice. (Supplementary Figure 4A, B). These synaptosomes were stained using antibodies against TH, the vesicular SNARE protein synaptobrevin-2 (VAMP2), and the active zone marker bassoon, and imaged using confocal microscopy (Figure 4C, Supplementary Figure 4C). The LRRK2^G2019S^ mutation did not affect the percentage of TH+ synaptosomes that contained VAMP2+ clusters (Supplementary Figure 4D). However, we found a significant decrease in the percentage of TH+/VAMP2+ synaptosomes that contained the active zone marker bassoon (Figure 4D). In addition, the remaining active zone clusters in LRRK2^G2019S^ TH+/VAMP2+ synaptosomes had an increased average area of bassoon staining when compared to wildtype (Figure 4E), a difference which was not observed in TH-/VAMP2+ synaptosomes (Supplemental Figure 4E).

To determine whether LRRK2 specifically impacts dopaminergic active zone sites rather than overall synaptic integrity in the striatum, we analyzed the levels of several synaptic markers in striatal synaptosomes. We found comparable levels of synaptophysin, VAMP2, synaptosomal-associated protein 25 (SNAP25), and the dopamine transporter (DAT) in LRRK2^G2019S^ as compared to those of wild-type synaptosomal extracts (Figure 4F-H, Supplementary Figure 4F-L). These data are in line with previous reports^34^ and indicate that LRRK2 does not affect the number of synaptic vesicles. Similarly, the levels of RAB3 proteins and of GDP dissociation inhibitor 1 and 2, which regulate RAB3 function68, remained unchanged across genotypes (Supplementary Figure 4F-L). However, we observed a decrease in RIM1 levels in the LRRK2^G2019S^ synaptosome fraction compared to controls (Figure 4I and Supplementary Figure 4F). In contrast, RIM1 levels in other striatal fractions or SNc lysates did not change (Figure 4J-L), suggesting that LRRK2 specifically affects the synaptic targeting of RIM proteins. Since RIM1 and Bassoon co-cluster to form the active zones,^64^ the reduction in synaptic RIM1 corresponds with our imaging analysis, which shows fewer bassoon+ clusters in TH+/VAMP2+ synaptosomes (Figure 4D). Overall, these findings suggest that dopamine axons in LRRK2^G2019S^ mice have fewer varicosities containing active zones, and the remaining active zones are disorganized.

### LRRK2 disrupts the number and composition of active zone release sites in vulnerable dopamine axons

Since vulnerable dopamine neurons show higher LRRK2 expression (Figure 1), we wondered whether changes in the active zone may be prominent in vulnerable Aldh1a1+ as compared to resilient Aldh1a1-dopaminergic axons in LRRK2^G2019S^ mice. To accomplish this, we used a previously described intersectional and subtractive approach^13,17^ that allows simultaneous labeling of Aldh1a1+ neurons with EYFP and Aldh1a1-neurons with mCherry (Figure 5A, Supplementary Figure 5A). Striatal brain sections were then stained with antibodies against bassoon to analyze the number and organization of active zone sites in Aldh1a1+ (EYFP+) or Aldh1a1-(mCherry+) axons from the same mouse (Figure 5B). Given the high abundance of bassoon signal in the striatum, we employed structured illumination microscopy (SIM) to achieve improved resolution, followed by 3D reconstruction analysis (Figure 5C). Using this imaging and analysis pipeline, along with the intersectional genetic approach, we were able to investigate active zone sites within vulnerable versus resilient dopamine axons. We observed a decrease in the density of bassoon+ clusters in vulnerable Aldh1a1+ but not resilient Aldh1a1-axons of LRRK2^G2019S^, compared to LRRK2^WT^ mice (Figure 5D). Additionally, we noted an increase in the volume of the remaining bassoon clusters in Aldh1a1+ but not Aldh1a1-dopamine axons in mutant LRRK2 compared to wild-type control mice (Figure 5E). We confirmed these findings using an independent mouse line, Anxa1Cre (Figure 5F-H). To further confirm that we are measuring bassoon clusters within the dopamine axons, we conducted additional analysis. First, we assessed the density of bassoon clusters associated with Anxa1+ dopaminergic axons. Next, we rotated the bassoon signal image by 180 degrees, keeping the GFP channel (TH axon) unchanged, and reassessed the density of bassoon clusters. If our measurements captured primarily bassoon clusters located outside of the axons, their density would not have changed with rotation. Instead, we observed a significant decrease in density (Supplementary Figure 5B), which suggests that our analysis mainly captures bassoon clusters within dopaminergic axons.

**Figure 5:**
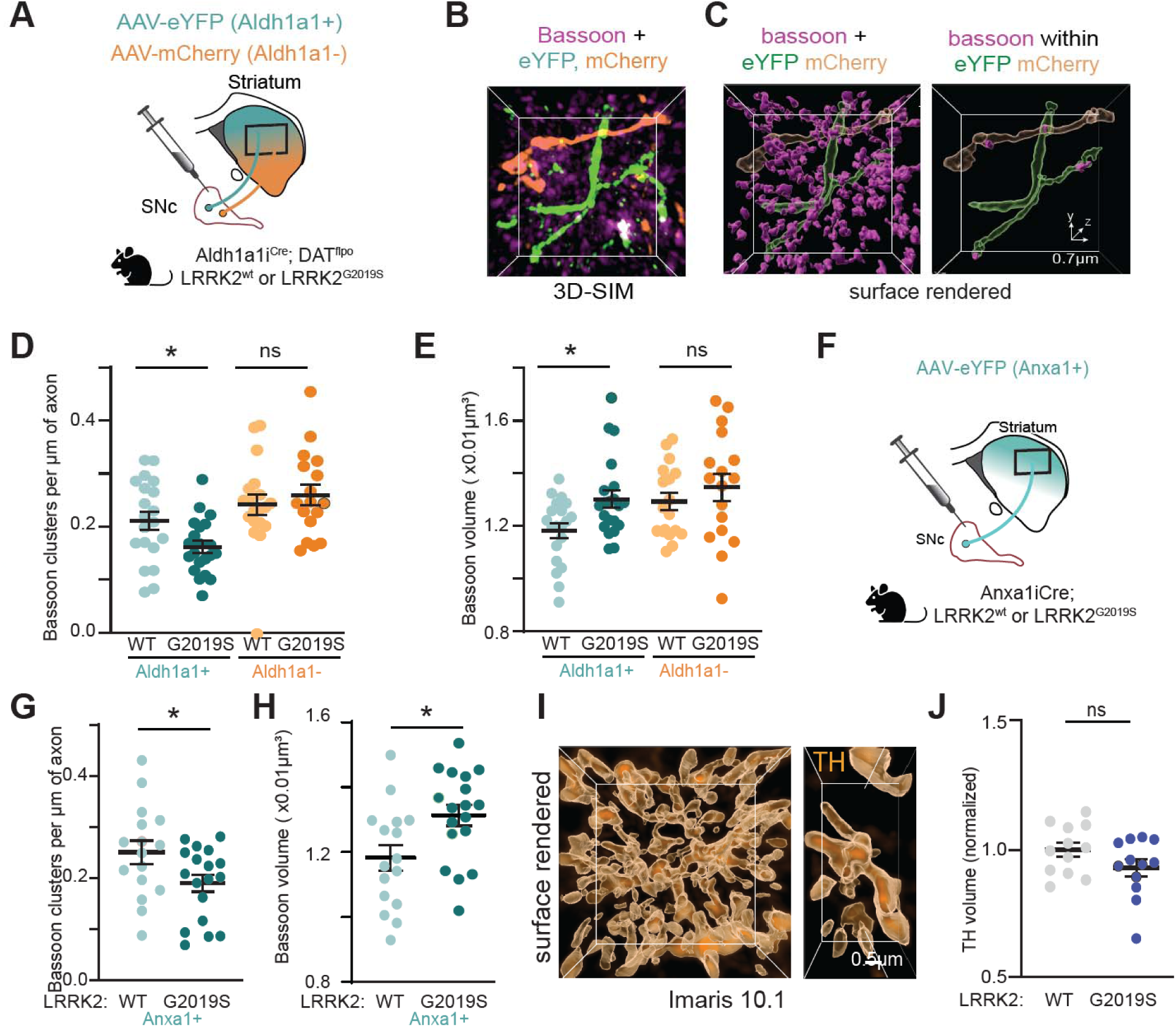
LRR K2 preferentially imp acts active zone release sites in vulnerable dopamine neurons. **A.** Simultaneous intersectional/subtractive viral strategy to label SNc Aldh1a1+ and Aldh1a1-dopamine neuron subtypes and their projections in the striatum of Aldh1a^Cre^; DAT^flpo^ LRRK2^WT^ and LRRK2^G2019S^ mice. **B, C.** Representative 3D-SIM image and its surface rendering in Imaris 10 showing distribution of bassoon clusters (magenta) in the whole field and within Aldh1a1+ (green) and Aldh1a1-(orange) axons in the dorsal striatal slices. **D.** Quantification of bassoon cluster density in Aldh1a1+ and Aldh1a1-dopamine axons. **E.** Quantification of bassoon cluster volume in Aldh1a1+ and Aldh1a1-dopamine axons. In D and E, each circle represents the average value from a section containing 25,000-35000 bassoon clusters; n=17-21 (4-5 sections/mouse, 4-5 mice/group). Data are represented as mean±SEM. Asterisk indicates statistical significance with Šídák’s multiple comparisons test following a two-way ANOVA test. \**p*<0.05. (D: Cell type factor F(1, 71)=12.75 p=0.0006, Genotype factor F(1,71)=1.277, p=0.2622, interaction F(1,71)=4.838 p=0.0311). E: Cell type factor F(1, 69)=4.607, p=0.0354, Genotype factor F(1,69)=1.277, p=0.0252, interaction F(1,69)=0.8688, p=0.3545 **F.** Strategy used to label SNc Anxa1i^Cre^ dopamine neuron subtypes and their projections in the striatum of Anxa1i^Cre^ LRRK2^WT^ and LRRK2^G2019S^ mice. **G.** Quantification of bassoon cluster density in Anxa1i^Cre^ dopamine axons. **H.** Quantification of bassoon cluster volume in Anxa1i^Cre^ dopamine axons. Each circle represents the average value from a section containing 15,000-40000 bassoon clusters; n=16-18 (4-5 sections/mouse, 4-5 mice/group). Data are represented as mean±SEM. Asterisk shows unpaired t-test. \**p*<0.05 **I.** Surface-rendered 3D image (Imaris 10.1) acquired with Nikon SoRa confocal instrument in the dorsal striatum showing TH immunostaining. **J.** Quantification of the total volume occupied by TH staining (axon fibers) in the dorsal striatum. Data represent mean±SEM. Each circle is the average of a section; n=12 (8 images/section, 3-4sections/mouse, 3-4 mice/group). No statistical significance after the unpaired t-test.

The observed active zone changes may result from the loss of TH fibers or alterations in dopamine vesicle clustering. To investigate this, we employed confocal imaging and 3D reconstructions to assess TH+ fibers in the dorsolateral striatum. We found no differences in the volume (Figure 5I, J, Supplementary Figure 5C), area, and intensity of TH signal in immunopositive ALDH1A1 fibers (Supplementary Figure 5D, E) in the dorsal striatum. We also did not observe any changes in the area or intensity of vesicular monoamine transporter 2 (VMAT2) vesicle clusters in LRRK2^G2019S^ mice compared with controls (Supplementary Figure 5D, F). These results, along with the unchanged percentage of TH+ striatal synaptosomes and unchanged levels of synaptic markers (Supplementary Figure 4), suggest that there is no significant loss of dopaminergic axons in young adult LRRK2^G2019S^ mice. Instead, the LRRK2^G2019S^ mutation disrupts the number and composition of active zones necessary for dopamine release, particularly in vulnerable dopaminergic axons. This aligns with previously reported primarily ex vivo decrease in dopamine release in the dorsolateral striatum,^35^ which is the primary projection site of these vulnerable neurons. ^13^

To identify the molecular factors underlying these structural alterations in mutant LRRK2 dopamine axons, we used a Cre-dependent virus to drive APEX2 proximity labeling within genetically targeted Anxa1 neurons in LRRK2^WT^ and LRRK2^G2019S^ mouse brains—with or without MLi-2—and coupled this with MS.^48^ Cre-negative mice injected with Cre virus were controlled for non-specific binding (Supplementary Figure 6A, B). We confirmed that our APEX2 dataset is enriched for dopamine neurons (compared to the publicly available APEX2 DA neuron proteome database).^48^ First, we validated the compartment specificity of our approach by comparing the enrichment of established somatodendritic versus axonal markers between the SNc and the striatum (Supplementary Figure 6C, D). Overall, the number of total proteins altered across different comparisons was limited. Specifically, 2% and 3% of all identified proteins in our dataset showed significant changes when comparing LRRK2^G2019S^ with LRRK2^WT^ and LRRK2^G2019S^ MLi-2-treated versus vehicle control, respectively (p-value < 0.05, |Log2FC| > 0.58). This pattern aligns with the limited number of proteins altered in a similar comparison in our striatal synaptosome-based experiments (Supplementary Figure 2D). Notably, among the significantly altered proteins were known components of active zones and presynaptic function (Supplementary Figure 6E). This is consistent with the observed structural changes (Figure 5G, H) and likely highlights potential compensatory mechanisms downstream of RAB3 phosphorylation that maintain presynaptic structure and function. In either case, the magnitude of effects in proteomic methods underscores that the primary impact of LRRK2 mutations in dopamine axons occurs through phosphoregulation, consistent with LRRK2 function as a kinase.

### In vivo dopamine release is reduced in LRRK2^G2019S^ compared to control mice

We next asked whether the changes in synaptic release machinery described here for Anxa1+ dopamine neurons in LRRK2^G2019S^ mice lead to any detectable changes in dopamine release in vivo. In LRRK2^WT^ or LRRK2^G2019S^ mice, we expressed the optogenetic actuator CHRmine^69^ in Anxa1+ dopamine neurons in the SNc and expressed the dopamine sensor GRAB-DA3m^70^ widely and non-specifically across the dorsal striatum (Figure 6A, Supplementary Figure 7A). We used fiber photometry to record dopamine release in the dorsal striatum evoked by optogenetic activation of Anxa1+ dopamine neuron cell bodies in awake, head-fixed mice on a treadmill (Fig. 6B, C).^13,71^ We found that evoked dopamine release was significantly reduced in LRRK2^G2019S^ compared to LRRK2^WT^ mice (Figure 6D, E). Specifically, 7 of 8 LRRK2^G2019S^ mice showed a statistically significant reduction in dopamine release compared with the control group (Figure 6E). This reduction was observed across a range of optogenetic stimulation powers (Supplementary Figure 7B, C). Importantly, we found that spontaneous locomotion-driven dopamine release was also significantly lower in LRRK2^G2019S^ compared to LRRK2^WT^ mice (Figure 6F). Taken together, these findings indicate that presynaptic anatomical changes in LRRK2^G2019S^ vulnerable dopamine release sites (Figure 5) correspond to a functional decline in striatal dopamine release. These results represent the first evidence of in vivo dopamine signaling deficits in the most vulnerable neuronal populations of LRRK2^G2019S^ mice.

**Figure 6.**
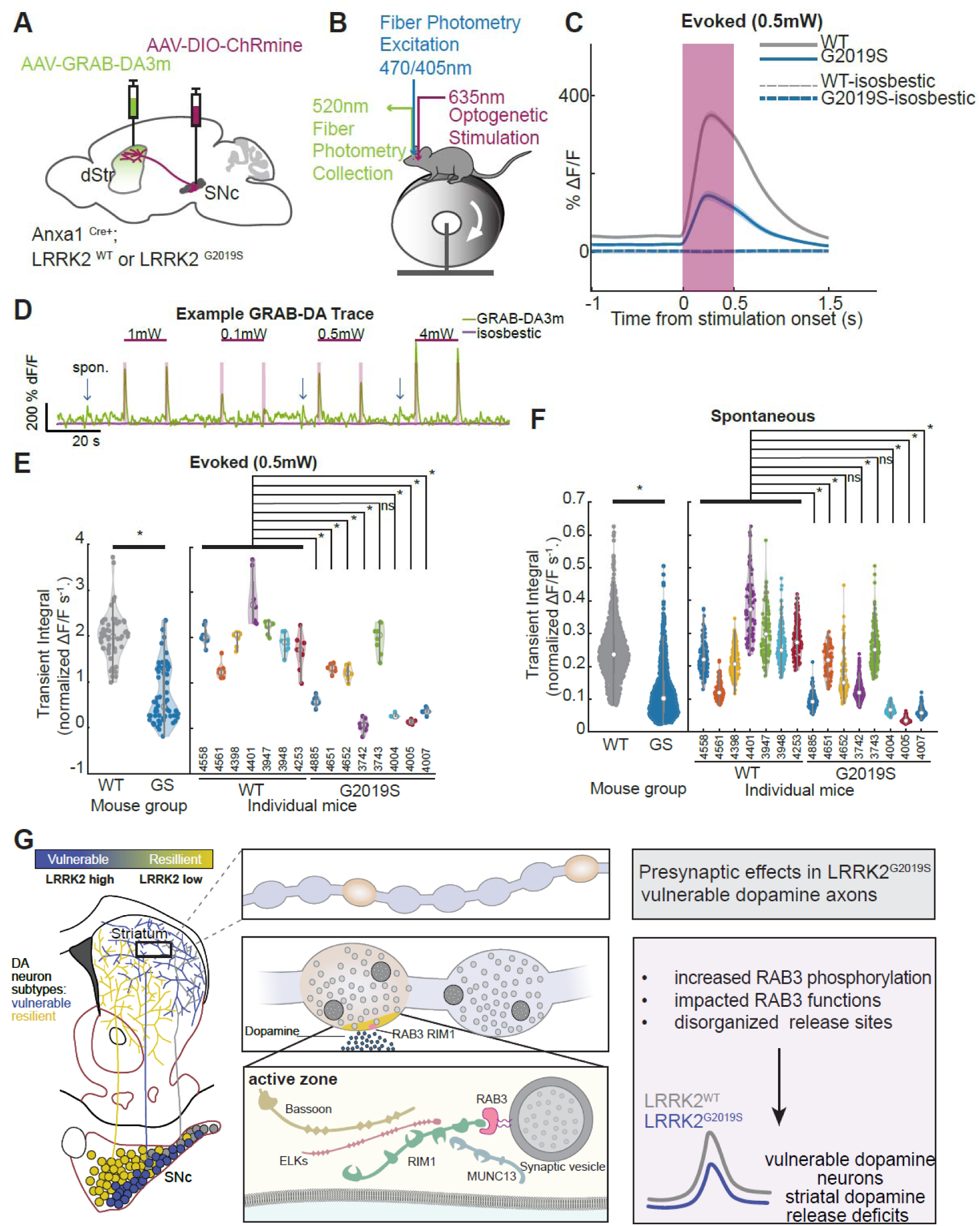
In vivo dopamine release is reduced in LRRK2^G2019S^ compared to control mice. **A.** Strategy used to label Anxa1+ dopamine neurons in the SNc of LRRK2^WT^ or LRRK2^G2019S^ mice with a red-shifted channelrhodopsin, ChRmine, and to express the fluorescent dopamine sensor, GRAB-DA3m, in the dorsal striatum (dStr) for measuring dopamine release (ΔF/F). **B.** Schematics of optogenetic stimulation (in SNc) and fiber photometry recording (dStr) setup. **C.** Evoked dopamine release triggered on optogenetic simulation: Average GRAB-DA3m ΔF/F triggered on optogenetic stimulation light onset with 0.5mW light power (blue trace, LRRK2^G2019S^ mice = 8, n = 64 stimulations; grey trace, WT mice = 7, n = 56 stimulations; red shade, light stimulation time): dashed lines, isosbestic control. Shaded regions denote meanl±lSEM across stimulations. **D.** Example GRAB-DA3m trace during ChRmine activation (shaded in red). The blue arrow indicates example spontaneous dopamine transients. **E.** Violin plots of optogenetically evoked (integrated) dopamine release response with 0.5mW stimulation light power (normalized integral of GRAB-DA3m ΔF/F during the optogenetic-stimulation light-on period). White dots denote group means and black bars denote 25th to 75th percentile. Left, group-level comparisons (LRRK2^G2019S^ n = 64 stimulations, WT n = 56 stimulations; p-value = 2.20E-20, unpaired t-test). Right, Violin plots of optogenetically evoked (integrated) dopamine release response in individual mice. Numbers denote mouse ID (n = 8 stimulations/mouse; p-value (4885) = 5.14E-10, p-value (4651) = 0.0008, p-value (4652) = 0.0001, p-value (3742) = 8.28E-15, p-value (3743) = 0.96, p-value (4004) = 5.62E-13, p-value (4005) = 3.44E-14, p-value (4007) = 4.72E-12). **F.** Violin plots of integrated spontaneous dopamine transients (normalized integral of GRAB-DA3m ΔF/F during significant dopamine transients selected by double-thresholding). Left, group-level comparisons (LRRK2^G2019S^ n = 1064 transients, WT n = 798 transients; p-value = 5.49E-17, unpaired t-test). Right, Violin plots of spontaneous (integrated) dopamine transients in individual mice. Asterisks (*) on the individual plots denote p-value < 0.00625 from Welch’s t-test with Bonferroni correction (α = 0.05/8 = 0.00625) comparing individual LRRK2^G2019S^ mice to the WT group. **G.** Summary graphical abstract of this study.

## Discussion

This study provides new mechanistic insights into how pathogenic LRRK2 mutations impair dopaminergic synaptic functions in PD. Using in vivo approaches, we show that the mutant LRRK2 disrupts the presynaptic architecture, specifically the active zone release sites that are essential for efficient neurotransmission.^64^ This disruption leads to reduced striatal dopamine release. By combining intersectional genetic strategies with real-time, behaviorally driven measurements of dopamine signaling in locomoting mice, our work goes beyond prior ex vivo studies ^35,37,72^ that lacked subtype specificity. We reveal a previously unrecognized deficit in both evoked and spontaneous dopamine release in awake and locomoting mice. We also uncover cell-type–specific mechanisms governing presynaptic dysfunction, highlighting that LRRK2-mediated effects are specific to defined subpopulations of dopamine neurons that are selectively vulnerable in PD. To our knowledge, this is the first demonstration of cell-autonomous LRRK2-mediated deficits in these vulnerable subpopulations, establishing a mechanistic framework for LRRK2-driven synaptic dysfunction in PD.

Recent single-cell transcriptomic analyses, together with studies in postmortem human tissue and preclinical models, have identified Aldh1a1_/Anxa1_ dopamine neurons as among the most vulnerable in PD.^11–13,16^ Our findings show that this ventral-biased subtype expresses elevated LRRK2 protein levels, further underscoring its relevance to disease pathophysiology. To investigate these implications further, we used targeted knock-in mice expressing the G2019S pathogenic mutation and applied intersectional labeling to study vulnerable subsets of dopamine neurons in the SNc, focusing on the Aldh1a1+ and Anxa1+ (a subset of Aldh1a1+) subpopulations. Super-resolution imaging revealed a selective reduction and disorganization of active zones in these neurons, effects not observed in less vulnerable subtypes, demonstrating a striking example of cell-type specificity.

Previous studies highlight unique features of dopamine neurons’ release sites compared with those of other synapses. Only 25-30% of dopamine neuron varicosities contain active zones, limiting presynaptic platforms for dopamine release.^64,66^ Functional studies confirm that about 20% of dopamine varicosities can release dopamine.^65^ Consequently, the precise regulation of these sparse release sites becomes increasingly crucial for dopaminergic synapses. Furthermore, dopamine synapses rely more heavily on RIM1 and RIM2 proteins than non-dopaminergic synapses,^64^ likely due to a lack of functional redundancy with other synaptic proteins (e.g., ELKS), as reported in hippocampal synapses.^73,74^ This suggests that active zone-like structures in dopamine neurons are molecularly and functionally specialized. Of note, the alterations in active zones observed in our study resemble, albeit to a lesser extent, those reported in conditional dopamine neuron–specific RIM1 and RIM2 knockout models.^64^ This raises the possibility that LRRK2 may directly or indirectly modulate RIM-dependent processes to influence release site integrity, thereby impacting fast dopamine signaling in vulnerable dopamine neurons. Consistent with the notion that active zone disruption pathology is a key component of PD, RIM1 and RIM2 have now been identified as PD genome-wide association (GWAS) loci. ^75,76^

Increased kinase activity is the pathogenic mechanism linked to LRRK2 mutations. Our findings in mice and humans show higher LRRK2 levels in vulnerable dopamine neurons, suggesting that enhanced phosphorylation of downstream LRRK2 targets may lead to presynaptic dysfunction. To investigate the molecular basis of these presynaptic defects, our phosphoproteomic analysis of striatal synaptosomes identified differentially phosphorylated RAB3 presynaptic proteins associated with distinct LRRK2 phosphorylation states, consistent with prior work indicating that RAB proteins are robust LRRK2 substrates across various cell types (non-dopamine neurons).^42,43^ We further found that increased phosphorylation of RAB3 in LRRK2^G2019S^ mice, which is reduced by LRRK2 kinase inhibition, decreases its interaction with its effector proteins RIM1 and RIM2, providing a mechanistic link between LRRK2 kinase activity and impaired active zone organization. This disruption likely affects synaptic vesicle trafficking ^54,56,77–79,^ thereby altering the probability of synaptic vesicle release. Support for this comes from studies in Drosophila showing that disruption of RAB3 function reduces the number of active zones while enlarging the remaining ones, a pattern that resembles our findings (Figures 4 and 5). Additionally, a recent study using induced pluripotent stem cell (iPSC)-derived LRRK2 neurons indicates that LRRK2-mediated RAB3 phosphorylation may hinder the axonal trafficking of synaptic vesicle precursors, ^80^ which could reduce the targeting of RIM1 and RIM2 to axonal varicosities. We found decreased RIM1 levels in striatal synaptosomes, but not in substantia nigra extracts from LRRK2^G2019S^ mice, indicating disrupted axonal targeting. Altogether, our studies propose that LRRK2-dependent phosphorylation of RAB3 alters the composition of active zones in vulnerable dopamine axons. Considering the necessity of striatal dopamine release on RIM1/2 proteins, ^64^ these changes likely underlie the in vivo deficits in dopamine release in these LRRK2mice (Figure 6). Impaired RAB-RIM interaction occurs only with phosphorylated RAB3 (Figure 3) and not with overexpressed phosphomimetic RAB3 constructs (Supplementary Figure 8), which have been used in various studies to probe the effects of altered phosphorylation.^80^ These findings, together with previous observations ^42,63^ underscore the significance of using phosphorylated RAB proteins (Figure 3) to better understand how phosphorylation status affects RAB-protein interactions.

Although LRRK2 is broadly expressed and influences various synapses ^81,82^, dopamine neurons may be uniquely vulnerable to presynaptic perturbations due to their extensive axonal arborization, high number of release sites, and reliance on a limited number of specialized release sites.^3,83^ In this context, our findings support a predominantly cell-autonomous mechanism, consistent with previous studies in mice in which mutant LRRK2 was selectively expressed in dopamine neurons and produced release deficits.^40^ Nevertheless, non–cell-autonomous influences within striatal circuits, such as astrocytes, interneurons, or striatal projection neurons^52,84–86^ cannot be ruled out. Further work should determine how these mechanisms interact across distinct neuronal populations.

By the time the symptoms appear in patients, it is estimated that dopamine terminal loss can be ∼60%, whereas dopamine cell bodies loss is approximately 30% ^87,88.^ Furthermore, imaging studies in non-manifesting LRRK2 mutation carriers ^89,90^ and in prodromal PD cohorts reveal presynaptic deficits prior to motor symptom onset ^21,91^. These findings in humans are also reflected in multiple genetic and toxin-based PD models, which show impairments in dopamine release and other presynaptic abnormalities before overt neuronal loss, supporting the idea that presynaptic dysfunction represents an early step in disease progression. For example, α-synuclein overexpression or auxilin knockout causes synaptic deficits before dopamine neuron loss. ^92,93^ Despite consistent evidence for synaptic alterations, the temporal sequence of events alone does not establish a causal link between synaptic dysfunction and dopamine neuron loss, which remains to be experimentally demonstrated. Most mouse models with LRRK2 mutations do not show neurodegeneration in the absence of additional stressors, reflecting the multifactorial vulnerability of dopamine neurons in the human condition. However, these models show in vivo deficits in dopamine signaling, which our data suggest may arise from LRRK2-dependent regulation of axonal substrates, such as RAB3 (Figure 2). The eventual loss of dopamine neurons may require additional somatic factors, including pathways regulating autophagy, lysosomal function, or mitochondrial integrity. Consistent with this possibility, our compartment-specific proteomic analyses reveal distinct changes in axons versus somata (ProteomeXchange dataset PXD065003), supporting the idea of compartment-specific substrates or LRRK2 effects. While our findings highlight the presynapse as a critical site of LRRK2 action, they do not directly address mechanisms of neuronal loss; rather, they provide mechanistic insight into how pathogenic LRRK2 disrupts dopamine signaling, capturing early pathogenic processes relevant to the initial stages of Parkinson’s disease.

Our results also have immediate translational relevance, as several LRRK2-targeted therapies are in clinical development as disease-modifying interventions. Elevated LRRK2 kinase activity has also been observed in idiopathic PD^29^, highlighting its broader role beyond familial forms of the disease. This mechanistic relevance is further supported by evidence of synaptic deficits in sporadic cases. ^91^ Importantly, ongoing clinical trials are evaluating LRRK2 inhibition in patients with sporadic PD, suggesting that LRRK2 targeting may have broad applicability. Future studies should determine whether LRRK2 kinase inhibition can restore dopamine release in vulnerable axons and examine how aging—a major risk factor for PD—modulates synaptic vulnerability and therapeutic responsiveness.

## Acknowledgments

This research was funded in whole or in part by Aligning Science Across Parkinson’s [D.A.D., R.A., L.P ASAP-020600] and [D.R.A ASAP-000463] through the Michael J. Fox Foundation for Parkinson’s Research (MJFF) and the UK Medical Research Council [MC_UU_00018/1] (D.R.A). A.R.K. was funded by Research Ireland through a Research Frontiers grant, 20/FFP-A/8446, and R.A. by R01NS119690. For the purpose of open access, the author has applied a CC BY public copyright license to all Author Accepted Manuscripts arising from this submission. Imaging work was performed at the Northwestern University Center for Advanced Microscopy (RRID: SRC_020996), generously supported by NCICCSGP30 CA060553, which was awarded to the Robert H. Lurie Comprehensive Cancer Center. Structured illumination microscopy was performed on a Nikon N-SIM system, purchased with NIH 1S10OD016342-01 support. Super-resolution spinning disk microscopy was performed on a Nikon SoRa system, purchased with NIH 1S10OD032270-01 support. Brain tissue from MLi-2-treated mice for the pRAB3A/RAB3A affinity MS experiment was kindly provided by Dr. Francesca Tonelli. We thank Dr. Matei Azcorra for providing some of the schematics.

## Authors contributions

Conceptualization, C.C., L.P.; scientific input, E.G., Y.K., D.A., A.K., D.R.A., D.A.D., S.H., and R.A.; methodology, C.C., E.H.G.T., E.N., M.J., O.R.M., H.S., Y.N., A.S., V.P., K.Q., C.N.; imaging, proximity labeling methods, C.C. Y.K., intersectional genetic strategies development, O.A.M.R, R.A.; in vivo evoked dopamine release, Q.H., K.Q., D.A.D.; biochemical methods, C.C., Y.N., S.H., in vitro phosphorylation of RAB3 protein, E.N., and A.K., mass spectrometry, M.J., and D.A.; data curation, formal analysis, C.C., Q.H., G.T., M.J., H.S., K.Q; writing, review and editing, all authors; supervision, L.P.

## Data sharing

The mass spectrometry proteomics data have been deposited in the ProteomeXchange Consortium via the PRIDE partner repository, with dataset identifiers PXD065003, PXD065006, and PXD0651255. Raw tubular Data from all experiments, all images, and the codes used in this study are available via the open-access option on the Zenodo repository (https://doi.org/10.5281/10.5281/zenodo.15635558) or GitHub. The protocols used in this study were uploaded to protocols.io; their DOIs are listed in the key resources table.

## Materials and methods

### 1. Animals

All experiments complied with the guidelines set by the National Institutes of Health and were reviewed by the Northwestern Animal Care and Use Committee. Adult male and female mice, aged 3 to 7 months, were used in all experiments. Structural studies were conducted in mice aged 3-5 months, while analyses of TH fibers were performed in mice aged 5-6 months. The in vivo dopamine release experiments included mice aged 6-7 months. The mice were group-housed on a standard 12-hour light/dark cycle and provided with standard feed. Littermates were randomly assigned to experimental procedures. Researcher blinding was implemented across experiments.

C57BL/6 mice (JAX:000664), homozygous KI mice expressing the pathogenic mutation LRRK2^G2019S^ (JAX:030961), and homozygous *Lrrk2^KO^* mice (JAX:016121) were purchased from Jackson Laboratory. The Aldh1a1i^Cre^ and Anxa1i^Cre^ lines were generated by the Transgenic and Targeted Mutagenesis Laboratory at Northwestern University and were characterized in a previous study;^13^ these lines were maintained in a heterozygous state. The DAT^Flpo^ line was generated by knocking in Flpo to the 3’ end of Dat coding sequences, following a P2A peptide, and will be described in detail in an upcoming study.

The Cre-dependent APEX2 mouse line (DIO-APEX2.NES-P2A-EGFP) was characterized in a prior study. ^46^ These APEX2-EGFP mice, bred to homozygosity, were crossed with the Dat^Cre^ (JAX:006660) to achieve expression of APEX2 in dopamine neurons of the substantia nigra compacta (SNc). All mice were backcrossed for several generations and maintained on a C57BL/6J background with either the LRRK2^WT^ or LRRK2^G2019S^ allele. Standard genotyping primers are available on the Jackson Laboratory website or in referenced studies. Further details on the mouse strains can be found in the resources table.

### 2. Immunofluorescence and confocal imaging in brain sections

Mice were perfused with 50 ml of PBS, followed by 50 ml of 4% paraformaldehyde in PBS. The brains were dehydrated using 30% sucrose in PBS for 48 hours and then sectioned coronally at a thickness of 30 μm with a cryostat (Leica Biosystems, CM305). The slices were collected in PBS containing 0.1% sodium azide and stored at 4°C for subsequent immunohistochemical analysis.

Striatal sections were incubated in 5% goat serum with 0.2% Triton X-100/PBS for 2 hours. Antigen retrieval for LRRK2 staining was performed by incubating the sections in antigen-unmasking solution (Vector, H-3300) at 100°C for 30 minutes, followed by cooling to room temperature for an additional 30 minutes. The sections were then washed with PBS for 5 minutes.

Next, the sections were incubated overnight at 4°C in 5% goat serum with 0.2% Triton X-100 in PBS. Then, they were incubated for an additional 48 to 72 hours at 4°C with primary antibodies: anti-TH (1:1000, SYSY), anti-ALDH1A1 (1:300, Abcam), anti-LRRK2/Dardarin clone N241A/34 (1:200, NeuroMab), anti-GFP (1:1000, Invitrogen). After washing with PBS, the sections were incubated in the same buffer used for primary antibodies with secondary antibodies—Alexa Fluor™ 488, Alexa Fluor™ 568, and Alexa Fluor™ 647 (1:300, Invitrogen)—for 3 hours. Subsequently, all slices were washed with PBS and mounted using ProLong™ Diamond Antifade Mountant. Detailed catalog information and identification numbers can be found in the resources table.

Confocal images of the fixed 30 μm-thick striatal and midbrain sections were obtained using the Nikon A1R microscope. Fluorescence images were captured using a 20x objective for LRRK2 expression, TH area, and intensity (Supplementary Figure 6), and a 100x objective for Th fluorescence at a resolution of 1,024 x 1,024 pixels, with a constant laser power maintained across all genotypes for each channel.

### 3. Stereotactic injections and viruses

For the intersectional labeling of Aldh1a1 and Anxa1 populations, adult mice expressing Aldh1a1i^Cre^;DAT^Flpo^ or Anxa1i^Cre^ were anesthetized with 3% isoflurane, which was maintained at approximately 1.5% during the procedure. Pain management included subcutaneous injections of Rimadyl (3 mg/kg) at the beginning of the surgery, followed by a slow-release injection of Buprenex (0.25 mg/kg) after the surgery, and an additional dose of Rimadyl (3 mg/kg) the following day.

The surgery commenced with the exposure of the skull, using the bregma as the reference point. A 0.5-1 mm hole was drilled into the skull at the stereotactic coordinates of the substantia nigra pars compacta (SNc) (RC: −3.16 mm, ML: - 1.50 mm, DV: −4.00 mm). A 0.5 µl Hamilton Neuros syringe was employed for precise delivery, with the needle left in place for 5 minutes after reaching the target depth. A volume of 0.3 µl of the viral vector was slowly injected into the SNc over 5 minutes, followed by another 5-minute wait to allow for proper viral diffusion within the brain parenchyma.

Two viral cocktail options were utilized: (i) a 1:1 mixture of AAV8 hSyn-Con/Fon-YFP (Addgene #55650, titer: 2.4x10¹³ GC/ml) and AAV8 Ef1a-Coff/Fon-mCherry (Addgene #137134, titer: 2.2x10¹³ GC/ml) delivered unilaterally for imaging experiments in Figure 5, or (ii) a pure injection of the AAV5-CAG-DIO-APEX2-NES (Addgene plasmid #79907, virus packaged by VectorBiolabs; titer: 3.8x10¹² GC/ml), delivered bilaterally for APEX2 proximity labeling. The viral constructs are detailed in the resources table. Following the injection, the cranial wound was closed with staples. The mice were perfused 4 weeks post-injection, and their brains were isolated for subsequent analysis.

### 4. APEX2 biotinylation in acute brain slices and tissue dissections

Mice were anesthetized with ketamine and underwent transcardial perfusion with 10 ml of ice-cold cutting solution consisting of 110 mM choline, 2.5 mM KCl, 1.25 mM monosodium phosphate, 10 mM glucose, 7 mM MgCl_2_, 0.5 mM CaCl_2_, 1.3 mM NaH_2_PO_4_, and 25 mM sodium bicarbonate, all saturated with 95% O_2_ and 5% CO_2_. The brains were quickly extracted and placed in ice-cold cutting solution, also saturated with 95% O_2_ and 5% CO_2_.

Coronal slices of 300 µm were prepared using a Leica VT1200S. After the slices were made in the cold cutting solution, they were transferred to a glass dish containing artificial cerebrospinal fluid (aCSF), which included 2.5 mM KCl, 10 mM glucose, 125.2 mM NaCl, 0.3 mM NaH_2_PO_4_, 1.3 mM MgCl_2_, 2.4 mM CaCl_2_, 26 mM NaHCO_3_, and 0.3 mM KH_2_PO_4_, supplemented with 0.5 mM biotin-phenol. The aCSF was continuously saturated with 95% O_2_ and 5% CO_2_. The slices were allowed to recover at room temperature for 60 minutes.

Following recovery, APEX2 labeling was initiated by adding 1 mM H_2_O_2_ to the aCSF at room temperature for 5 minutes. To quench the labeling, the slices were rapidly transferred to a separate glass dish containing quenching aCSF, which consisted of the aforementioned aCSF supplemented with 10 mM Trolox, 20 mM sodium ascorbate, and 10 mM sodium azide, for 5 minutes.

The slices were then rapidly dissected in ice-cold quenching aCSF. The tissues were flash-frozen in liquid nitrogen and stored at −80°C until further use in WB or proteomic experiments.

### 5. Protein extraction and streptavidin enrichment of APEX2 samples

Frozen APEX2-labeled tissues were homogenized on ice using a glass Dounce homogenizer with 30 strokes of both A and B pestles. The lysis was performed in 0.75 ml of ice-cold tissue lysis buffer, which contained 50 mM Tris (pH 8.0), 150 mM NaCl, 10 mM EDTA, 1% Triton X-100, 5 mM Trolox, 10 mM sodium ascorbate, 10 mM sodium azide, and Halt protease and phosphatase inhibitor cocktail (Thermo Fisher Scientific). After adding 39 μl of 10% SDS to achieve a final concentration of 0.5%, the lysates were rotated for 15 minutes at 4°C. The lysates were then clarified by centrifugation at 21,000 × g for 10 minutes at 4°C. The supernatants were transferred to a new prechilled Eppendorf tube for trichloroacetic acid (TCA) precipitation (for LC-MS) or stored at −80°C (for WB).

To precipitate proteins from the lysates, an equal volume of ice-cold 55% trichloroacetic acid (TCA) was added. The samples were incubated on ice for 30 minutes, followed by centrifugation at 21,000 × g for 10 minutes at 4°C. The protein pellets were then resuspended in 1 ml of acetone prechilled to −20°C and centrifuged again under the same conditions. This resuspension and centrifugation process was repeated three additional times using 1 ml of acetone prechilled to −20°C, for a total of four washes. After removing any residual acetone, the protein pellets were resuspended in Urea Dissolve Buffer, which contained 8 M urea, 1% sodium dodecyl sulfate (SDS), 100 mM sodium phosphate (pH 8), and 100 mM ammonium bicarbonate (NH_4_HCO_3_). The pellets were dissolved by sonication for 1 minute, followed by gentle agitation on an orbital shaker for 1 hour at room temperature. A small aliquot (5%) of the resuspended protein was flash-frozen and stored at −80°C. The samples were diluted with equal volumes of water to achieve a final concentration of 4 M urea and 0.5% SDS.

Streptavidin magnetic beads (Thermo Fisher #88817) were resuspended and washed three times in Urea Detergent Wash Buffer (4 M urea, 0.5% SDS, 100 mM sodium phosphate, pH 8) for 15 minutes at 4°C. After washing, the streptavidin beads were resuspended in ice-cold Urea Detergent Wash Buffer, and 50 μl containing 0.5 mg of beads was added to each sample. Proteins were incubated overnight with streptavidin beads on a rotary wheel at 4°C. After 14 to 18 hours, the unbound supernatant was discarded, and the beads were resuspended in 1 ml of Urea Detergent Wash Buffer and transferred to a new tube. The beads were washed three times for 10 minutes in 1 ml of Urea Detergent Wash Buffer at room temperature.

After the third wash, the beads were resuspended in 1 ml of Urea Wash Buffer (4 M urea, 100 mM sodium phosphate, pH 8) and transferred to a new tube. This washing step was repeated three times for 10 minutes in 1 ml of Urea Wash Buffer at room temperature. The beads were then resuspended in 200 μl of Urea Wash Buffer and transferred to a new tube. The buffer was removed using a magnetic stand, and the beads were flash-frozen and stored at −80°C.

### 6. Western blot analysis

#### Brain extracts

Tissues were extracted and homogenized in 1x cell lysis buffer (Cell Signaling Technologies) supplemented with the Halt protease and phosphatase inhibitor cocktail (Thermo Fisher Scientific) using pellet pestles for 30 seconds. The protein concentration was determined using BCA Protein Assays (Thermo Scientific). Equal amounts of protein (30 μg of total tissue lysate) were separated by 4–12% NuPAGE Bis-Tris PAGE (Invitrogen) and transferred to membranes using the iBlot2 nitrocellulose membrane blotting system (Invitrogen) according to the manufacturer’s protocol.

#### Synaptosomes

Subcellular fractionation of mouse striatum was performed as previously described.^64^ Specifically, mouse striata were dissected and rapidly homogenized in 1ml iced-cold homogenizing buffer (4mM HEPES, 320 mM Sucrose, and Halt protease supplemented with phosphatase inhibitor cocktail (Thermo Fisher Scientific)) using a Teflon homogenizer (15 strokes). The homogenized brain extract was centrifuged at 1,000×g for 10 min at 4_. The supernatant (S1) was collected and centrifuged at 12,500× g for 15 min at 4 °C. The supernatant (S2) was removed, and the pellet (P2) was re-homogenized in 1 ml homogenizing buffer with 10 strokes in a Teflon homogenizer. After the addition of 1 ml homogenizing buffer, the P2 homogenate was added to the top of a sucrose gradient made of 5ml 1.2M sucrose and 5ml 0.8M sucrose, and was centrifuged at 69,150×g (SW41) for 70 min at 4 °C. The synaptic plasma membrane fraction (SPM) in the interphase between two sucrose fractions was collected using a syringe and transferred to clean ultracentrifuge tubes, and the samples were centrifuged in a swinging bucket rotor (SW55) at 200,000×g for 30 min. The pellet was resuspended in SDS lysis buffer (1% SDS, 10mM EDTA, 50mM Tris) supplemented with the Halt protease and phosphatase inhibitor cocktail (Thermo Fisher Scientific) or in 1x Phosphate-buffered saline (Sigma-Aldrich) for imaging. The protein concentration was determined using the BCA Protein Assay (Thermo Scientific), and 5 μg were separated by 4–12% NuPAGE Bis-Tris PAGE (Invitrogen) and transferred to membranes using the iBlot2 nitrocellulose membrane blotting system (Invitrogen) according to the manufacturer’s protocol.

#### APEX2 based experiments

The protein concentration of frozen tissue lysates was determined using the Bradford assay (Thermo Scientific) and 10 μg APEX2 labeled tissue lysate were loaded into 4–12% NuPAGE Bis-Tris PAGE (Invitrogen) as input. Frozen streptavidin beads were resuspended in ∼20 μl of 1× SDS sample buffer supplemented with 20 mM DTT and 2 mM biotin. Samples were boiled for 5 min at 95°C to elute biotinylated proteins. Beads were immediately placed onto a magnetic rack, and the entire sample was loaded into 4–12% NuPAGE Bis-Tris PAGE (Invitrogen) and transferred to membranes using the iBlot2 nitrocellulose membrane blotting system (Invitrogen) according to the manufacturer’s protocol.

The membranes were incubated with primary antibodies specific for streptavidin-HRP (1:1000, Abcam), LRRK2/Dardarin clone N137/6 (1:1000, NeuroMab), LRRK2/Dardarin clone N241A/34 (1:1000, NeuroMab), TH (1:1000, Millipore), RAB3A (1:1000, Sigma), RAB3C (1:1000, Proteintech), synaptophysin (1:1000, Cell Signaling), VAMP2 (1:1000, R&D Systems), SNAP25 (1:1000, Proteintech), DAT (1:1000, Millipore), pS1292 LRRK2 (1:100, Abcam), pS935 LRRK2 (1:100, Abcam), RIM1 (1:1000, Proteintech), GDI1 (1:1000, Sigma), GDI2 (1:1000, Novus), phospho-RAB12 (1:1000, Abcam), total Rab12 (1:1000, Proteintech), and β-actin (1:3000, Sigma) overnight at 4°C. Following this, the membranes were incubated with secondary anti-mouse or anti-rabbit antibodies (1:2000, Thermo Scientific) for 1 hour at room temperature. Afterward, they were incubated with Immobilon ECL Ultra Western HRP Substrate (Millipore) for 3 minutes prior to image acquisition.

Chemiluminescent blots were imaged using the iBright imaging CL1500 system (Thermo Fisher Scientific). Further details on the chemicals and antibodies can be found in the resource table.

#### Phos Tag gels

SNc tissues extracted and homogenized in EDTA-free Lysis buffer (1% Triton X-100, 1% glycerol, 150 mM NaCl, 25 mM HEPES, 1.5 mM MgCl_2_) using pellet pestles (30 seconds). The lysis buffer was supplemented with Halt protease and phosphatase inhibitor cocktail (Thermo Fisher Scientific). The protein concentration was determined using BCA Protein Assays (Thermo Scientific). A total of 10 μg of lysate were separated by 12.5% SuperSep^TM^ Phos-tag^TM^ (Wako). Gels were washed 3 times with 1x TG transfer buffer (Fisher BioReagents) supplemented with 10 % Methanol and 10 mM EDTA, with each wash lasting 10 minutes. After the washes, the gels were incubated with 1x TG transfer buffer with 10% Methanol for an additional 10 minutes and then transferred to Nitrocellulose membranes (Thermo Scientific) using the Mini Gel Tank and Blot Module (ThermoFisher) at 15V for 3 hours.

The membranes were incubated in 3% BSA in 1x Tris buffered saline (Sigma) supplemented with 0.1% Tween 20 for 1 hour, followed by incubation with primary antibodies specific for RAB3A (1:1000, Sigma) and RAB3C (1:1000, Proteintech) at 4 °C overnight, followed by probing with secondary anti-rabbit antibodies (1:2000, Thermo Scientific), as described above. Chemicals and antibodies are detailed in the resource table.

### 7. Striatal synaptosome preparation for imaging

Synaptosomes were diluted 5,000 times in 1x PBS, spun down at 4,000 rpm for 10 min at 4°C on #1.5 poly-d-lysine coated coverslips and fixed for 10 minutes in 4% PFA in PBS. Synaptosomes were blocked in 5% donkey serum in PBS for 1 hour and permeabilized in 0.5% Triton X-100 in PBS for 10 minutes. Incubation with primary antibodies (VAMP2 1:200, R&D; TH 1:500, SYSY; bassoon 1:300, Enzo) was carried out in 0.1% Triton X-100, 1% donkey serum PBS overnight at 4°C. After three 5-min washes in 1X PBS, synaptosomes were incubated with the secondary antibodies (Donkey anti-Rabbit Alexa Fluor™ 488; Donkey anti-goat Alexa Fluor™ 647; Cy™3 AffiniPure™ Donkey Anti-Guinea Pig) in 0.1% Triton X-100, 1% donkey serum PBS for 90 min and mounted (ProLong™ Diamond Antifade Mountant) onto Superfrost™ slides. A single optical section for each area of synaptosomes was acquired with the confocal Nikon A1 laser scanning microscope system using a 100X 1.49NA objective.

### 8. Imaging analysis

#### LRRK2 signal in SNc sections

The protein signals for TH, ALDH1A1, and LRRK2 were measured using Imaris 10.1 software (Bitplane, Concord, USA). The surface rendering function was used to segment TH/ALDH1A1 cells. Background subtraction was enabled; the diameter for the largest sphere was set to 10 μm and automatically thresholded, with a smoothing surface set to 3 μm. Mean intensity of ALDH1A1 protein within the TH surface above 1.5 times the average TH protein channel mean intensity was considered positive. The LRRK2 intensity within TH-positive cells was measured automatically by the software.

#### Synaptosome analysis

Synaptosome imaging analysis was performed using FIJI software.^94^ To define synaptosomes that are positive for a specific protein, background was subtracted from all channels using the “rolling ball” FIJI plugin with a radius of 33.0 pixels. Automatic segmentation was applied with the following parameters: default threshold, particle size ranging from 0.04 to 0.40, and circularity between 0.30 and 1.00. This process allowed the definition of regions of interest (ROIs) first in the VAMP2 channel. The average intensity of the signal was measured within each ROI, and only those ROIs that were positive for VAMP2 were retained. An ROI is classified as containing the target protein if the average fluorescence intensity of that protein within the ROI is more than twice the average intensity of all pixels in the image. Conversely, an ROI was considered negative for the target protein if the intensity within the ROI fell below this threshold. To identify synaptosomes that contained both VAMP2 and TH, the intensity of the TH signal within the VAMP2-positive ROIs was measured. The intensity of the bassoon signal was measured in the double-positive ROIs, retaining only those that were triple-positive.

To assess the bassoon area, automatic segmentation was used to define the ROIs set both for bassoon and TH (threshold: default; analyze particles: size 0.04-0.40 for bassoon and size 0.06-0.60 for TH; circularity: 0.30-1.00). These bassoon and TH ROI sets were then combined to define the overlapped ROIs, using the “AND” function in the ROI Manager. The intensity of the VAMP2 signal helped define the bassoon/TH overlapped ROIs that were positive for VAMP2. Finally, the average area value of bassoon localized within TH and VAMP2 double-positive synaptosomes was calculated using the “Set Measurement” and “Measure” FIJI functions. The percentage of bassoon positive clusters within VAMP2 or double TH/VAMP2-positive was calculated based on the intensity signal within ROIs as described above.

#### Striatal TH expressing fibers analysis

After performing perfusion and immunofluorescence as described above, dorsal striatal sections were stained using the primary antibodies: anti-TH (1:1,000, SYSY), anti-ALDH1A1 (1:300, Abcam), and anti-VMAT2 (1:200, Immunostar). The resource table provides detailed catalog and identification numbers for these antibodies.

Analysis was performed in fluorescence images acquired using different microscopes and settings as follows: <u>a.Volume</u> <u>measurements of TH fibers.</u> Nikon CSU-W1 SoRa with a 60x objective at 0.1 mm intervals at 1,024 x 1,024 pixel resolution. TH protein signal was measured using Imaris 10.1 software (Bitplane, Concord, USA). Surface rendering function was used to segment the TH fibers. Background subtraction was enabled, the diameter of the largest sphere was set at 0.805 μm and threshold set at 2,000. Segments were filtered with “Number of voxels Img=1” set above 50. TH volume was measured automatically by software.

b. <u>Intensity and area of TH expressing axons</u>. To eliminate any confounding factors related to magnification settings, we measured TH axons using both high and low magnification. For the high magnification analysis (Figure 5), we utilized a Nikon A1R microscope with a 100x oil immersion objective (Numerical Aperture = 1.45) and a resolution of 1,024 x 1,024 pixels to assess the intensity of TH axons in the striatum across different genotypes. Rolling ball background subtraction was performed using Nikon Elements software. The protein signals were analyzed using Imaris 10.1 software (Bitplane, Concord, USA), utilizing the surface rendering function to segment TH, ALDH1A1, and VMAT2 proteins. Background subtraction was disabled and automated thresholds were applied with manual adjustment made as needed. Segments were filtered with “Number of voxels Img=1” set above 50. For the ALDH1A1 positive TH axon, filtering was applied based on the “overlapped area ratio to surface surface=TH-ALDH1A1” above 0.4. Similarly, VMAT2 within TH axonal terminals was filtered with “overlapped area ratio to surface surface= TH-VMAT2” above 0.4. Area and intensity were measured automatically by software. For the low magnification analysis (Supplementary Figure 6), images were captured using the same Nikon A1R microscope with a low magnification (20x) objective (NA=0.75), with a 1,024 x 1,024 pixel resolution. Stitched images of the whole striatum were automatically generated with Nikon Element. The TH protein signal was analyzed with Imaris 10.1 software (Bitplane, Concord, USA) employing the surface rendering function to segment TH fibers. Background subtraction was disabled and the threshold was set to 1,500 with manual adjustments made if needed. Segments were filtered with “Number of voxels Img=1” set above 50. TH area and intensity were measured automatically by software.

### 9. LC-MS sample preparation

#### Synaptosomes

Mouse striata were dissected two hours after administering a dose of MLi-2 (10 mg/kg) or control vehicle via oral gavage. The tissues were flash frozen and stored at −80°C until use. Subcellular fractionation of the mouse striatum was performed as previously described.^95,96^

In this process, three mouse striata were pooled and rapidly homogenized in four volumes of ice-cold Buffer A (0.32 M sucrose, 5 mM HEPES, pH 7.4, 1 mM MgCl_2_, 0.5 mM CaCl_2_) supplemented with a Halt protease and phosphatase inhibitor cocktail (Thermo) using a Teflon homogenizer with 12 strokes. The homogenized brain extract was then centrifuged at 1,400 g for 10 minutes. The supernatant (S1) was saved, and the pellet (P1) was homogenized in Buffer A with a Teflon homogenizer (five strokes). After centrifugation at 700 g for 10 minutes, the supernatant (S1’) was pooled with S1.

Next, the pooled S1 and P1 were centrifuged at 13,800×g for 10 minutes, resulting in a crude synaptosomal pellet (P2) and its corresponding supernatant (S2). The P2 pellet was resuspended in Buffer B (0.32 M sucrose, 6 mM Tris, pH 8.0), also supplemented with the protease and phosphatase inhibitor cocktail, using a Teflon homogenizer (five strokes). It was then carefully loaded onto a discontinuous sucrose gradient (0.8 M/1 M/1.2 M sucrose solution in 6 mM Tris, pH 8.0) with a Pasteur pipette, followed by centrifugation in a swinging bucket rotor at 82,500×g for 2 hours.

The synaptic plasma membrane fraction (SPM) located at the interphase between the 1 M and 1.2 M sucrose fractions was collected using a syringe and transferred to clean ultracentrifuge tubes. Samples were then centrifuged in a swinging bucket rotor at 200,000×g for 30 minutes. The supernatant was removed and discarded, while the SPM pellet was flash frozen and stored at −80°C.

The SPM pellet samples were processed by Tymora Analytical Operations in West Lafayette, Indianapolis. For the lysis process, 200 µL of phase-transfer surfactant lysis buffer (PTS), which contains 12 mM sodium deoxycholate, 12 mM sodium lauroyl sarcosinate, 10 mM TCEP, and 40 mM CAA, was added to each tissue sample. A phosphatase inhibitor cocktail 3 (Millipore-Sigma) was also included.

After the initial lysis, the samples were incubated for 10 minutes at 95°C, pulse-sonicated several times, and then incubated for an additional 5 minutes at 95°C. The lysed samples were centrifuged at 16,000 × g for 10 minutes to remove debris, and the supernatant was collected. The samples were diluted fivefold with 50 mM triethylammonium bicarbonate, and a BCA assay was performed to determine protein concentration, normalizing all samples by protein amount.

The samples were then normalized to 300 mg protein in each and digested with 6 mg Lys-C (Wako) for 3 h at 37°C. 6 mg trypsin was added for overnight digestion at 37°C. The supernatants were collected and acidified with trifluoroacetic acid (TFA) to a final concentration of 1% TFA. Ethyl acetate solution was added at a 1:1 ratio to the samples. The mixture was vortexed for 2 min and then centrifuged at 16,000 × g for 2 min to obtain aqueous and organic phases. The organic phase (top layer) was removed, and the aqueous phase was collected, dried completely in a vacuum centrifuge, and desalted using Top-Tip C18 tips (Glygen) according to manufacturer’s instructions. The samples were dried completely in a vacuum centrifuge and subjected to phosphopeptide enrichment using PolyMAC Phosphopeptide Enrichment kit (Tymora Analytical) according to manufacturer’s instructions, and the eluted phosphopeptides dried completely in a vacuum centrifuge.

The full phosphopeptide sample was dissolved in 10.5 μl of 0.05% trifluoroacetic acid with 3% (vol/vol) acetonitrile and 10 μl of each sample was injected into an Ultimate 3000 nano UHPLC system (Thermo Fisher Scientific). Peptides were captured on a 2-cm Acclaim PepMap trap column and separated on a 50-cm column packed with ReproSil Saphir 1.8 μm C18 beads (Dr. Maisch GmbH). The mobile phase buffer consisted of 0.1% formic acid in ultrapure water (buffer A) with an eluting buffer of 0.1% formic acid in 80% (vol/vol) acetonitrile (buffer B) run with a linear 90-min gradient of 6–30% buffer B at flow rate of 300 nL/min. The UHPLC was coupled online with a Q-Exactive HF-X mass spectrometer (Thermo Fisher Scientific). The mass spectrometer was operated in the data-dependent mode, in which a full-scan MS (from m/z 375 to 1,500 with the resolution of 60,000) was followed by MS/MS of the 15 most intense ions (30,000 resolution; normalized collision energy - 28%; automatic gain control target (AGC) - 2E4, maximum injection time - 200 ms; 60sec exclusion).

#### APEX2

This analysis was performed in Biognosys AG (Schlieren, Switzerland). Samples (beads with proteins attached via streptavidin pulldown as described in section 5) were solubilized and digested overnight with sequencing grade trypsin (Promega) in a urea-containing denaturation buffer. Each sample is the eluted proteins from one mouse: 4-8 mice/genotype, treatment combination as shown in Supplementary Figures 1 and 6 except Cre-control mice n=3 mice/condition. Beads were collected on a magnetic rack, supernatant was transferred to a new tube and used for the clean-up. Purification for mass spectrometry was carried out using Oasis HLB μElution Plate 30μm plate (WATERS) according to the manufacturer’s instructions. Peptides were dried down to complete dryness using a SpeedVac system and dissolved in LC solvent A (1 % acetonitrile in water with 0.1 % formic acid) containing Biognosys’ iRT-peptide mix for retention time calibration. Peptide concentrations in mass spectrometry ready samples were measured using the mBCA assay (Thermo Scientific Pierce). For DIA LC-MS/MS measurements, peptides were injected on an in-house packed reversed phase column on a Thermo Scientific EASY-nLC 1200 nano-liquid chromatography system connected to a Thermo Scientific Orbitrap Exploris 480 mass spectrometer equipped with a Nanospray Flex ion source and a FAIMS Pro ion mobility device (Thermo Scientific). LC solvents were A: water with 0.1 % FA; B: 80 % acetonitrile, 0.1 % FA in water. The nonlinear LC gradient was 1 – 50 % solvent B in 171 minutes followed by a column washing step in 90 % B for 7 minutes, and a final equilibration step of 1 % B for 0.5 column volumes at 64 °C with a flow rate set to a ramp between 450 to 271 nl/min (min 0: 450 nl/min, min 172: 271 nl/min, washing at 400 nl/min). The FAIMS-DIA method consisted per applied compensation voltage of one full range MS1 scan and 34 DIA segments, as adapted from previous work.^97,98^

### 10. MS Data analysis

#### Synaptosomes

The raw files were searched directly against the mouse database with no redundant entries, using Byonic (Protein Metrics) and Sequest search engines loaded into Proteome Discoverer 2.3 software (Thermo Fisher Scientific). The data from the two search engines were combined. In most cases, the same peptides were identified by both. The final data is from both search engines, including identifications reported by only one search engine. MS1 precursor mass tolerance was set at 10 ppm, and MS2 tolerance was set at 20 ppm. Search criteria included a static carbamidomethylation of cysteines (+57.0214 Da), and variable modifications of phosphorylation of S, T and Y residues (+79.996 Da), oxidation (+15.9949 Da) on methionine residues and acetylation (+42.011 Da) at N terminus of proteins. Search was performed with full trypsin/P digestion and allowed a maximum of two missed cleavages on the peptides analyzed from the sequence database. The false-discovery rates of proteins and peptides were set at 0.01. All protein and peptide identifications were grouped, and any redundant entries were removed. Only unique peptides and unique master proteins were reported. All data were quantified using the label-free quantitation node of Precursor Ions Quantifier through the Proteome Discoverer v2.3 (Thermo Fisher Scientific). For the quantification of phosphoproteomic data, the intensities of phosphopeptides were extracted with initial precursor mass tolerance set at 10 ppm, minimum number of isotope peaks as 2, maximum ΔRT of isotope pattern multiplets – 0.2 min, PSM confidence FDR of 0.01, with hypothesis test of ANOVA, maximum RT shift of 5 min, pairwise ratio-based ratio calculation, and 100 as the maximum allowed fold change. For calculations of fold-change between the groups of proteins, total phosphoprotein abundance values were added together, and the ratios of these sums were used to compare proteins within different samples. This analysis identified 2,155 unique phosphoproteins, 8,008 unique phosphopeptides that map to 2,155 unique phosphoproteins, 24,238 unique peptides that map to 3,130 unique proteins. The threshold for significance was set to p<0.05 (unadjusted p-value) & FC (difference on the log2 intensity values) ≥ 0.58. To capture a greater phosphorylation for GO-term and kinase-substrate analysis, the statistical cut-off for inclusion was set at uncorrected p > 0.05.

#### APEX2 study

The DIA mass spectrometric data were analyzed using Spectronaut software (Biognosys, version 19.0).^93^ The false discovery rate on peptide and protein level was set to 1 %. A mouse UniProt .fasta database (Mus musculus, 2024-07-01) was used for the search engine, allowing for 2 missed cleavages, carbamidomethylation of cysteine as fixed modification and up to 5 variable modifications (N-terminal acetylation, methionine oxidation, deamidation of asparagine or glutamine and phosphorylation at serine or threonine or tyrosine).

HRM mass spectrometric data were analyzed using Spectronaut software (Biognosys, version 19.0). The false discovery rate on peptide and protein level was set to 1 %, and data were filtered using row-based extraction. The directDIA+ library generated in this project was used for the analysis. The HRM measurements analyzed with Spectronaut were normalized using global median normalization.

Proteins that were not enriched (enriched: q-value < 0.05, |log2FC| > 0) in Anxa1^Cre^; LRRK2^G2019S^ vehicle vs. non-transgenic control were removed from the dataset of samples from SNc and striatum, respectively. The filtered dataset was used for further analysis. For testing of differential protein abundance, protein intensities for each protein were analyzed using a two-sample Student’s t-test. The following thresholds were applied for candidate identification: p-value < 0.05; absolute average log2 ratio > 0.58 (fold-change > 1.5). Distance in heat maps was calculated using the “manhattan” method, and clustering was performed using “ward.D” for both axes. Principal component analysis was conducted in R using prcomp and a modified ggbiplot function for plotting.

### 11. SIM imaging and analysis

Aldh1a1^Cre^-DAT^flop^ or Anxa1^Cre^ mice crossed with LRRK2^G2019S^ mice or control littermates were labeled with viruses as described in sterotactic injections and viruses sections and processed as in Immunofluorescence and confocal imaging in brain sections. They were probed with the primary antibodies anti-EGFP (1:1000, Invitrogen), anti-mCherry (1:1000, Invitrogen), and anti-bassoon (1:300, Enzo). Detailed catalog and identification numbers are shown in the resource table.

Multichannel SIM images were obtained with a Nikon N-SIM Structured Illumination super-resolution microscope in the Center for Advanced Microscopy at the Nikon Imaging Center at Northwestern University. The imaging was conducted with a 100x objective lens with a numerical aperture (NA) of 1.4. ^62^ The acquisition settings were set to 10 MHz, and 16 bit depth with electron multiplication (EM) gain and no binning. Exposure times ranged between 200 and 900 ms, with the EM gain multiplier kept below 1,500. The conversion gain was maintained at 1x. The laser power was adjusted to keep LUTs within the first quarter of the scale. LUTS were kept similar across comparisons.

Single images were processed and analyzed using Nikon Elements software. 3D reconstructions in a single plane used nine images captured with 2D SIM and reconstructed with 3D SIM to increase xy and z resolution, were generated by Nikon Element, and the illumination modulation contrast was set automatically by the software. Images were cropped using 3D crop function to remove the edges, and a resolution of 1,950x1,950 was applied for further analysis with Imaris 10.1. The segmentation of bassoon particles and axons was conducted using the surface creation tool. In the surface wizard, the source channel for axons was selected, corresponding to either Alexa488 (EGFP) or Alexa568 (mCherry), with object-object statistics activated.

To create a clean surface of axon surface, the smoothing surface detail option was enabled (Surface Grain Size = 0.0643 µm). Background subtraction was performed using local contrast with a diameter of Largest Sphere set to 0.241 µm for better separation of the background. The auto-threshold value was applied, with user adjustments made as required. The segments were then filtered by setting the “Number of Voxels Img=1,” above the lower automatic threshold. The settings were saved and manually checked and validated with 10 different images before they were applied to all the images in the dataset. The fluorescence signal outside the surface was removed by using the “Mask channel” function, selecting the corresponding channel for either the Alexa488 (EGFP) or Alexa568 (mCherry), and voxel intensity outside the surface was set to 0.

Axon terminal segments’ length was measured with the filaments tool. In the tool wizard, the source channel for axonal terminal segments was again selected, corresponding to the masked Alexa488 (EGFP) or masked Alexa568 (mCherry), with the seed points threshold set at 6,000. Machine learning pixel classification was performed by training Imaris to discard unwanted segments (not overlapping with the surface) while keeping segments that overlapped with the surface until they covered over 99% of the visible surface. The settings were again saved and verified using 10 randomly chosen images before being applied to all images without further user adjustments.

The source channel for bassoon particles was selected based on Alexa647, with object-object statistics on. To create a clean surface of bassoon particles, the smoothing surface detail was activated (Surface Grain Size = 0.06 µm). Background subtraction (local contrast) (Diameter of Largest Sphere =0.15 µm) was employed to separate the background. An auto-threshold value was applied with user adjustments as required. Additionally, the Region Growing Estimated Diameter was set to 0.100 µm based on intensity to help split touching objects. The segments were then filtered by “quality” between lower and upper automatic thresholds, followed by a further “volume” filter between 0.003 and 0.04 µm³. Bassoon clusters within axonal terminals were filtered based on “overlapped volume ratio to surface,” where the ratio of EGFP/mCherry was set above 1.0. These settings were saved and, as before, checked with 10 different randomly chosen images before being applied to all the images.

The number of bassoon objects, their volume, and the length of axon terminal segments were automatically measured and calculated. Density calculations were performed in Excel using the number of bassoon objects relative to the total length of axonal terminal segments, with bassoon object number and axonal terminal segment length determined in Excel.

### 12. p-RAB3A vs. RAB3A interactome experiment

#### Preparation of recombinant p-RAB3A (pT86)

The cDNA for His-FLAG-RAB3A (residues 18-190, Q81L) and MST3 kinase (residues 1-431), codon optimized for *E.coli* expression, was ordered from Genscript and cloned into the pET28a vector at the NdeI/BamH1 sites. FLAG-RAB3A and MST3 were expressed in 2xYT and Lysogeny Broth (LB) respectively, supplemented with 30 mg/mL kanamycin at 37 °C. At an OD600 of 0.6, FLAG-RAB3A was induced with 0.5 mM IPTG, and MST3 was induced with 0.1 mM IPTG. The cells were then grown overnight at 18°C. Cells were harvested by centrifugation and lysed by sonication in extraction buffer (20 mM Tris-Cl, pH 8, 300 mM NaCl, 20 mM imidazole, 10 mM β-mercaptoethanol, 5 mM MgCl_2_). Following centrifugation at 20,000 x g for 40 minutes at 4_ to remove cellular debris, the supernatants were applied to a gravity flow column containing Ni^2+^-agarose resin. The resin was washed with an excess of extraction buffer and wash buffer (extraction buffer supplemented with 40 mM imidazole) and then eluted in extraction buffer supplemented with 200 mM imidazole. The His-tag was removed from FLAG-RAB3A by overnight dialysis at 4_ in gel filtration buffer (20mM Tris-HCl, pH 8.0, 100 mM NaCl, 1 mM DTT, 5 mM MgCl_2_) with thrombin protease, followed by a second Ni^2+^-column. FLAG-RAB3A, not intended for phosphorylation, was further purified by gel filtration chromatography using a Superdex 75 column (10/300 GL) equilibrated in gel filtration buffer. To generate phospho-FLAG-RAB3A, purified FLAG-RAB3A was incubated with MST3 (8:1 wt/wt ratio) in phosphorylation buffer (50 mM Tris–HCl, 150 mM NaCl, 10 mM MgCl_2_, 2 mM ATP, pH 7.5), similar to previous protocols for other RABs at room temperature for 40 minutes.^58,59^ The reaction generated more than 50% phosphorylated RAB3A, and the mixture was subsequently exchanged into a low-salt buffer (10 mM Tris-HCl, pH 7, 10 mM NaCl, 1 mM DTT, 5 mM MgCl2) for separation by ion-exchange chromatography on a Mono Q 5/50 GL column (GE Healthcare). Phospho-FLAG-RAB3A was separated from the non-phosphorylated species using a 0-35% gradient from low-to-high salt buffer (low salt buffer supplemented with 1M NaCl) and confirmed by PhosTag gel electrophoresis. Finally, phospho-FLAG-RAB3A was further purified by gel filtration chromatography as previously described.

#### Immunoprecipitation of p-RAB3A and RAB3A interactors

p-FLAG RAB3A and FLAG-RAB3A were used as ‘bait’ for the immunoprecipitation of proteins from mouse brain lysate to identify binders that interact differentially. Brains from mice administered a subcutaneous injection of MLi-2 (30 mg/kg, 2 hours). Whole brain was isolated snap frozen in liquid nitrogen and stored at −80° C. The brains were homogenized in lysis buffer (50 mM Tris-HCl pH 7.4, 150 mM NaCl, 10% [w/v] glycerol, 10 mM sodium-glycerophosphate, 10 mM sodium pyrophosphate, 0.5% [v/v] NP40 - supplemented with Complete Mini [EDTA-free] protease inhibitor (Merck), PhosSTOP phosphatase inhibitor (Merck), 5 mM MgCl_2_). The lysate was clarified by centrifugation at 17,000×g for 15 minutes at 4°C and quantified using the Bradford assay. Phospho-FLAG-RAB3A and FLAG-RAB3A were bound to PierceTM Anti-DYKDDDDK Magnetic Agarose at 4°C on a rotary wheel for 1 hour. The supernatant was removed, and the beads were washed twice with PBS-T (PBS supplemented with 0.01% Tween-20, 5 mM MgCl_2_). Brain tissue lysate (750 mg) was added to the beads in a 500 ml volume and incubated on a rotary wheel for 1 hour at 4°C. The supernatant was removed, and the beads were washed twice with high-salt lysis buffer (lysis buffer supplemented with 500 mM NaCl), followed by regular lysis buffer, and then frozen at −80°C for mass spectrometry analysis, as described below.

#### Animals

For affinity MS pulldown experiments of pRAB3A/RAB3A interactors, 12 C57BL/6j male mice at 10 weeks of age were used. Mice were maintained under specific pathogen-free conditions at the University of Dundee (UK). All animal studies were ethically reviewed and carried out by the Animals (Scientific Procedures) Act 1986 and regulations set by the University of Dundee and the U.K. Home Office. Animal studies were approved by the University of Dundee ethical committee and performed under a U.K. Home Office project license. Mice were housed at an ambient temperature (20–24°C) and humidity (45–55%) and were maintained on a 12 hr light/12 hr dark cycle, with free access to food and water. Mice were injected subcutaneously with vehicle (40% [w/v] 2-hydroxypropyl)-β-cyclodextrin (Sigma-Aldrich #332607) or MLi-2 dissolved in the vehicle at a 30 mg/kg final dose. Mice were sacrificed by cervical dislocation 2 hours following treatment, and the collected tissues were rapidly snap frozen in liquid nitrogen for downstream analysis.

#### Sample preparation for quantitative proteomics

Immunoprecipitation samples were eluted from magnetic beads in 40 μL of lysis buffer containing 2% SDS (w/v), 20 mM HEPES (pH 8), with a complete EDTA-free protease inhibitor cocktail (Roche) and PhosSTOP phosphatase inhibitor cocktail tablets (Roche). Samples were reduced with 10 mM TCEP for 30 minutes at 60°C, followed by alkylation of cysteine residues with 40 mM IAA for 30 minutes at 25°C in the dark. 20% SDS was then added to achieve a final concentration of 5% SDS in the samples, which were subsequently acidified by the addition of Trifluoroacetic acid to a final concentration of 1%. Samples were subsequently processed for on-column tryptic digestion using a micro S-trap (Protifi, USA). Briefly, samples were diluted 6-fold with wash buffer (90% methanol, 10% 100 mM TEABC) and loaded onto microcolumns, with centrifugation at 1000g for 1 minute, and the flow-through discarded. After sample loading, the S-Trap columns were washed four times with 150 μL wash buffer, followed by centrifugation at 1000g for 1 minute. On-column digestion was performed by incubating 40 μl (1 μg) of Trypsin/Lys-C mix (MS grade, Promega, UK) in 50 mM TEABC solution (pH 8) at 47°C for 1 hour and 20 minutes, followed by incubation at 22°C overnight. The samples were then eluted with the addition of 40 μl 50mM TEABC (pH 8), 40 μl 0.15% (v/v) formic acid (FA), and 3x 40 μl 80% acetonitrile (ACN), 0.15% FA. Peptides were then dried using a vacuum centrifuge at room temperature and stored at −20°C until mass spectrometry analysis.

#### Quantitative Proteomic Analysis

Peptides were resuspended in 0.1% formic acid supplemented with 0.015% N-Dodecyl-β-D-Maltoside (DDM). Approximately 200 ng of peptides were analyzed with a Vanquish Neo nano-liquid chromatography system in-line with an Orbitrap Astral mass spectrometer (ThermoScientific). Peptides were trapped and eluted using an Acclaim™ PepMap™ 100 C18 HPLC column (3 μm particle size, 75 μm diameter, 150 mm length) and separated using an EASY-Spray™ PepMap™ Neo UHPLC column (2 μm C18 particle, 75 μm diameter, 150 mm length) with buffer A: 0.1% formic acid and buffer B: 80%ACN, 0.1% formic acid. Peptides were separated across a 13-minute gradient as follows: 0-0.7 min, 1% buffer B, 1.8 μl/min; 0.7-1.0 min, 4% buffer B, 1.8 μl/min; 1.0-7.7 min, 8% buffer B, 1.8 μl/min; 7.7-11.4 min, 22.5% buffer B, 1.8 μl/min; 11.4-11.8 min, 35% buffer B, 1.8 μl/min; 11.8-12.3 min, 55% buffer B, 2.5 μl/min; 12.3-13 min, 99% buffer B, 2.5 μl/min.

Samples were analyzed in DIA mode, with MS1 scans performed at a resolution of 240,000 across an m/z range of 380-980, using a normalized AGC target of 500% and a maximum injection time of 3 milliseconds. DIA scans were performed on the precursor mass range with an isolation window of 4 m/z, a scan range of 150-2,000 m/z, and a normalized collision energy of 25%.

#### Raw Mass Spectrometry Data Analysis

Raw data were searched using DIA-NN (version 1.9.2) against a predicted spectral library generated from the reviewed mouse UniProt database (downloaded January 2023; 17,124 entries with isoforms). Spectra were searched with strict Trypsin specificity (cleavage at K or R residues), allowing a maximum of one missed cleavage. Peptides with an amino acid length of 7-30 were considered using default settings, with cysteine carbamidomethylation enabled as a fixed modification.

#### Statistical Analysis and Data Visualization

Statistical analysis and visualization of DIA-NN output files were performed with Python (version 3.11.5, Jupyter Notebook; Project Jupyter) using packages pandas (2.0.3), matplotlib (3.7.2), seaborn (0.12.2), numpy (1.26.1), and then uploaded to the CURTAIN web tool (https://curtain.proteo.info/#/).^60^ Proteins identified by a single peptide were excluded, with log2-transformed protein intensities then filtered by 100% detection in at least one sample condition. Missing values were imputed by random draws from a Gaussian distribution centered at the 1% quantile of global protein group intensities per sample. The Limma package (version 3.56.2; R, version 4.3.1) was used to fit the data to a linear model using the lmfit function, with eBayes correction to compute moderated t-statistics and false-discovery rate controlled by Benjamini-Hochberg correction.

#### Confirmation of in vitro p-RAB3A phosphorylation site with MS

1 µg of purified phosphorylated FLAG-Rab3a protein was diluted 1:39 in 50 mM triethylammonium bicarbonate (TEABC, pH 8) in triplicate. Triplicate samples of the non-phosphorylated FLAG-RAB3A protein were also prepared as controls. Protein was reduced by adding Tris(2-carboxyethyl) phosphine (TCEP) to a final concentration of 10 mM at 60°C with mixing at 1,100 rpm (ThermoMixer C, Eppendorf) for 30 minutes. Samples were then alkylated by the addition of Iodoacetamide (IAM) to a final concentration of 40 mM at 25°C with mixing at 1,100 rpm for 30 minutes in the dark. 100 ng of Trypsin/Lys-C MS grade protease (Pierce, ThermoScientific) was then added to samples for tryptic digest at 37°C overnight (approximately16 hours), followed by acidification with 1% formic acid to a final concentration of 0.1%. Samples were then dried at room temperature and resuspended in 0.1% formic acid with 0.015% n-dodecyl-β-d-maltoside (DDM) in LC-grade water, with agitation at 1,800 rpm for 30 minutes at room temperature. Peptides were desalted using C18 EvoTips (EvoSep) and eluted in 40% acetonitrile. Samples were again dried using a vacuum centrifuge and stored at −20°C. For mass spectrometry analysis, samples were resuspended in 0.1% formic acid with 0.015% n-dodecyl-β-d-maltoside (DDM) in LC-grade water. 5 ng of peptides from each sample were injected into a Vanquish Neo nano-liquid chromatography system, in-line with an Orbitrap Astral mass spectrometer (Thermo Scientific), as previously described.

Raw data were searched against a fasta file generated from the Flag-RAB3A protein sequence using DIA-NN (version 1.9.2), with phosphorylation set as a variable modification (UniMod:21 with mass delta 79.9663 at STY, maximum number allowed=1) and precursor FDR set at 0.01. DIA-NN matched the expected phosphorylation site of peptide: YRT(UniMod:21)ITTAYYR, in the 2+ charge state. This was confirmed by manual inspection of the precursor and fragment peptide traces in Skyline (idotp=0.99, Skyline version 24.1). The DIA-NN spectral database search also identified another phosphorylation site in the peptide TSFLFRY(UniMod:21)ADDSFTPAFVSTVGIDFK, in the 4+ charge state. However, manual inspection of this peptide trace in Skyline confirmed this to be a false positive, as no precursor ion was found within the retention time window to match the observed fragmentation spectra traces (idotp=0).

### 13. Experiments in HEK-293 cells

#### Cell culture

HEK-293 cells were cultured in low glucose DMEM (Gibco, #11885-084) supplemented with 10% fetal bovine serum (Gibco, #A5256801), 1% MEM non-essential amino acids (Gibco, #11140050), and 1% penicillin/streptomycin (Gibco, #15140122). At 90% confluency, cells were trypsinized with 0.25% trypsin/EDTA (Gibco, #25200072) and plated at a 1:4 dilution into a 100 mm tissue culture dish.

RAB3, RIMS1, and pCMV plasmids were transformed into competent bacteria (One Shot TOP10 chemically competent E. coli, Invitrogen, #C404003). Single colonies were selected and cultured in 3 mL of LB medium supplemented with the corresponding antibiotics for 8 h at 37 °C. After that, 1 mL of bacterial culture was grown in 200 mL of LB medium plus antibiotics overnight at 37°C. Plasmids were purified using ZymoPURE II Plasmid MidiPrep kits (Zymo Research, #D4201) according to the manufacturer’s instructions.

#### Transfection

For cotransfection of RAB3 plasmids and RIMS1, HEK-293 cells at 80% confluency were plated into 6-wells at a 1:20 split ratio. The following day, cells were co-transfected with HA-RAB3 constructs and myc-flag-RIMS1, or with myc-flag-RIM1 alone. For each 6-well, 200 ng of HA-RAB3 plasmids and 2 μg of myc-flag-RIM1 plasmid were mixed in a 1.5 ml centrifuge tube in 50 μl high glucose DMEM (Gibco, #11965092), and 6 μl of LipoD293 (Signagen Laboratories, #SL100668) was mixed in another 1.5 ml centrifuge tube in 50 μl high glucose DMEM. Three minutes later, the contents were mixed, briefly vortexed, and incubated at room temperature for 15 minutes. The media was exchanged to fresh complete medium, and the plasmid/LipoD293 mix was added dropwise to the wells. 24 hours after transfection, cells were transferred into 100 mm dishes and grown for an additional 24 hours.

For the determination of isoform-specificity of the RAB3 antibodies, HEK-293 cells at 80% confluency were plated into 6-well plates. The following day, they were co-transfected with HA-RAB3 constructs (200 ng) and pCMV (2 μg), or with pCMV alone (2 μg) as described above. After 24 hours, the transfected cells were transferred into 100 mm dishes and grown for an additional 24 hours.

#### Immunoprecipitation

For HA-RAB3 and myc-flag-RIM1 co-transfections, HEK293 cells were collected 48 h after transfection. For each 100 mm dish, cells were washed with ice-cold 1 x PBS, and lysed with 1 ml of IP lysis buffer (20 mM Tris-HCl pH 7.5, 150 mM NaCl, 1 mM EDTA pH 8.0, 0.3% Triton-X100, 10% glycerol, and 1X cOmplete, Mini, EDTA-free protease inhibitor cocktail) in a 1.5 ml low protein binding tube for 30 min on a rotary wheel at 4 °C. Lysates were centrifuged at 10,000 rpm for 10 minutes at 4 °C, and 50 μl of the supernatant set aside as immunoprecipitation input. Magnetic anti-HA beads (25 μl) (Pierce) were washed for 30 min with IP wash buffer (20 mM Tris-HCl pH 7.5, 150 mM NaCl, 1 mM EDTA pH 8.0, 10% glycerol) on a rotary wheel at 4°C. Beads were pulled down with magnet, followed by equilibration in IP lysis buffer for 3 min on ice. Beads were pulled down again and incubated with 1 ml of lysed supernatant for 1h at 4 °C on a rotary wheel. Beads were washed three times for 5 min with 1 mL of IP wash buffer, and proteins were eluted with 40 μL of 2x Laemmli sample buffer and boiled at 95°C for 5 min to release bound proteins.

For the determination of RAB3 antibody specificity, cells were washed and collected 48 h after transfection in 1 ml of IP lysis buffer. Lysates were incubated for 30 minutes on a rotary wheel at 4°C, and then centrifuged for 10 minutes at 10,000 rpm at 4 °C. For each transfection, five μl of supernatant was resolved on SDS-PAGE gels and analyzed by WB as described above.

### 14. In vivo evoked dopamine release

#### Surgery

Anxa1iCre; LRRK2^WT^ and Anxa1iCre; LRRK2^G2019S^ mice were used for virus injection and fiber implantation at 6 months of age. Mice were anesthetized with isoflurane (5% at induction and 1–2% for maintenance). Two craniotomies, 0.5–1mm in diameter, were made over the right substantia nigra (−3.20_mm caudal, +1.6_mm lateral from bregma) and the right rostral striatum (+0.5_mm caudal, +1.8_mm lateral from bregma). A small volume (0.4_μl total) of virus (pAAV-Ef1a-DIO-ChRmine-mScarlet-WPRE, Addgene, 130998-AAV5, titer 2.20E+13 vg/mL), diluted 1:1 in PBS, was pressure injected through a pulled glass micropipette into the SNc at four depths (−3.8, −4.1, −4.4, and −4.7_mm ventral from dura surface, 0.1_μl per depth). Similarly, 0.1_μl of virus (pAAV-hSyn-GRAB-gDA3m, Addgene, 208698-AAV1, titer ≥ 7×10¹² vg/mL), diluted 1:2 in PBS, was injected into the striatum at −1.9 mm ventral from the dura surface. Following injections, a 4mm long, 400 um diameter optic fiber (MFC_400/430-0.66_4.0mm_TS3.0_C60, Doric Lenses) for opto-stimulation was implanted through the same craniotomy just above the SNc, and a 2mm long, 200-diameter optic fiber (MFC_200/250-0.66_2.0mm_ZF1.25(G)_FLT, Doric Lenses) for fiber photometry was implanted in the dorsal striatum. The fibers and a custom-made metal headplate were secured on the skull with metabond (Parkell).

For details, see the online protocol at dx.doi.org/10.17504/protocols.io.e6nvwd6qdlmk/v1.

#### Fiber photometry and optogenetics setup

A custom-made photometry setup, based on a previously published design^13^ was used for recording. Blue excitation light (470-nm LED, Thorlabs, M70F3) and purple excitation light (for the isosbestic control) (405-nm LED, Thorlabs, M405FP1) were coupled into the optic fiber such that a power of 0.5_mW emanated from the fiber cannula tip. Then, 470-nm and 405-nm excitation were alternated at 100_Hz using a waveform generator, each filtered with a corresponding filter (Semrock, FF01-406/15-25 and Semrock, FF02-472/30-25) and combined with a dichroic mirror (Chroma Technology, T425lpxr). The excitation light was collimated and coupled to a fiber-optics patch cord (MBP_400/430/3000-0.57_1m_FCM-2xZF1.25(F), Doric Lenses), which connects to the fiber cannula implant. Green fluorescence was separated from the excitation light by a dichroic mirror (Chroma Technology, T505lpxr) and further filtered (Semrock, FF01-540/50-25) before collection using a GaAsP PMT (H10770PA-40, Hamamatsu; signal amplified using Stanford Research Systems SR570 preamplifier). A red LED light for opto-stimulation (LEDFRJ_635, Doric Lenses) was similarly coupled to a fiber-optic patch cord and connected to implanted fiber-optic cannulas. The frequency, power, and duration were constructed as stimulation trains and controlled by Doric Studio software (Doric Lenses) using complex waveforms. The stimulation trains were triggered by a custom LabVIEW script, which introduces random time delays between every train within a time interval. LED powers at the tip of the cannula were measured with a digital power meter (PM100D, ThorLabs). The rotational velocity of the treadmill during animal locomotion was sampled at 1 kHz by a rotary encoder (E2-5000, US Digital) attached to the treadmill’s axle, using a custom Arduino script. A DigiData data acquisition system (Axon Digidata 1550B, Molecular Devices) was used to record and synchronize fluorescence and optogenetic light triggers at a sampling rate of 2_kHz.

#### In vivo optogenetic activation of Anxa1+ DA neurons

4 weeks after surgery, mice were head-fixed with their limbs resting on a one-dimensional cylindrical Styrofoam treadmill ∼20_cm in diameter by 13_cm wide. A separate red LED light was placed in front of their face to mask the red light delivery to the brain. After 2 days of habituation, GRAB-DA3m signals were recorded for 20 minutes. During recordings, TTL-triggered red LED stimulation was delivered in pseudorandom order at 0.1, 0.5, 1, and 4 mW. Each pulse train consisted of 8-ms on, 8-ms off cycles for 500 ms. Stimuli were spaced at least 20 seconds apart and repeated 8 times per power level.

#### Fiber photometry signal processing

Simultaneous traces of optogenetic stimulation triggers, treadmill velocity, green fluorescence (520 nm), excitation light function generator, and camera triggers were collected as times series at 2kHz from DigiData as ABF files. The files are then loaded into MATLAB and analyzed using custom scripts similar to previously described. ^13^ The 470-nm and 405-nm excited fluorescence was first separated based on function generator signals and then all traces were re-binned at 100 Hz for analysis. The fluorescence generated from 470 nm excitation was used for functional measurements, while the non-functional fluorescence generated from 405 nm excitation (isosbestic point) was used as a control for movement artifacts. Raw fluorescence traces first underwent correction procedures to eliminate background fluorescence and slow drifts. Background fluorescence was estimated from cortex recordings where there is no GRAB-DA labeling and calculated as a percentage of the baseline (defined as the 8th percentile of total fluorescence using a 20-s sliding window). Background fluorescence was subtracted from 470 nm and 405 nm independently. The background-subtracted traces were normalized by baseline (newly defined as the 8th percentile of the background-subtracted trace using a 20-s window) division. To calculate the ΔF/F, the normalized traces were then converted to % ΔF/F by subtracting the baseline (defined as the 8th percentile of the whole trace) and dividing by it. For triggered averages of ΔF/F on opto-stimulation light-on periods, integral values were normalized to a range of −0.25 to 1 for plotting.

#### Spontaneous dopamine transient selection

Spontaneous GRAB-DA3m signals were recorded in the same sessions as evoked responses and isolated by removing time points corresponding to optogenetic stimulation from the ΔF/F trace, including a buffer of 1 s before and 1.5 s after each stimulation. The remaining ΔF/F segments were concatenated and normalized by z-scoring. Significant spontaneous transients were identified using a double-threshold applied to the z-scored signal: a low threshold (z < −1) defined the onset and offset of each transient, and crossing a high threshold (z > 1) was required for each transient selection. Thus, each detected transient began and ended at the low threshold and contained at least one time point exceeding the high threshold. For each identified transient epoch, the area under the curve was calculated using the original (non-z-scored) ΔF/F signal and normalized to the same unit as evoked integrals.

#### Quantification and statistical analysis

Data were analyzed using custom code in MATLAB deposited in Github. Link can be found in the resource table.

For tests of statistical significance, all *P* values reported were calculated using a 2-sample t-test (ttest2 function in MATLAB). For individual-to-group comparisons, the Welch t-test was used because of the assumption of unequal variance. The specific test used is stated in the main text and figure legends where *P* values are reported. All tests were corrected for multiple comparisons using the Bonferroni correction: multiplying the *P* values by the number of comparisons made.

No statistical methods were used to predetermine sample sizes; however, our sample sizes are similar to those reported in previous publications. Surgeries, data collection, and signal processing were performed in a blinded manner, concerning the mouse type (LRRK2^WT^ or LRRK2^G2019S^). Post-hoc histological verification of virus expression and fiber placement were used as exclusion criteria, which led to the exclusion of one G2019S mouse due to the abnormality of GRAB-DA3m and ChRmine expression.

### 15. Pipelines for proteome analysis and data visualization

#### Gene Ontology

(GO) analysis, related to Biological Process, Cellular Components, and Molecular Function, was evaluated using the clusterProfiler 4.16.6 package and the org.Mm.eg.db 3.20.0 database in R 4.4.3. Other packages such as readxl/1.4.5, openxlsx/4.2.8, dplyr/1.1.4, and ggplot2/3.5.1 were used for filtering and visualization. Pathways with an adjusted p-value <0.05, based on a hypergeometric test and a Benjamini-Hochberg correction for multiple testing, are considered significantly enriched for the specified terms.

The functional analysis of significantly altered phosphopeptides was performed in proteins with at least one peptide showing significant differences in its phosphorylation status between synaptosomes of vehicle-treated and MLi2-treated LRRK2^G2019S^ mice. Significance was defined as |log_2_FC| > 0.58 and p ≤0.05 as determined by Student’s t-tests. The top 15 pathways are specified in Figures 2C and Supplementary Figure 2C. The functional analysis of differentially expressed proteins was performed on proteins with a significant difference in the APEX2 proteomes of vehicle-treated vs MLi2-treated LRRK2^G2019S^ mice. Significance was defined as |Log_2_FC| > 0.58 and p ≤0.05 as determined by two-sample Student’s t-tests. Selected pathways relevant to presynaptic functions are shown in Supplementary Figure 6E.

The complete list of analyses will be uploaded to Zenodo and in Supplementary File 1 and Supplementary File 2 for synaptosome experiment (Figure 2) and APEX2 experiment (Supplementary Figure 6), respectively.

#### Volcano plots

All quantified peptides in the phosphopeptidome analysis comparing vehicle-treated and MLi2-treated LRRK2^G2019S^ mice were graphed according to their Log2FC and -log(p-value), as determined by multiple unpaired t-tests. All proteins identified in the proteomics analysis of vehicle- and MLi2-treated LRRK2^G2019S^ mice were plotted according to their Log2FC and log (p-value), as determined by multiple unpaired t-tests. Visualization was performed using the ggplot2 R package in R version 4.4.3. Phosphopeptides and proteins with |Log_2_FC| > 0.58 and the p≤0.05 were considered significantly altered.

Heatmaps were generated using the ComplexHeatmap R package.

#### Plotting human genes in dopamine neurons

Feature plots were generated using R (RRID: SCR_001905) ShinyCell package^98^ (RRID: SCR_022756) to display the expression of LRRK2 and RAB3 genes in molecularly defined subclusters of dopamine neurons from a single-nuclei RNA dataset from Kamath et al.^49^ (GEO Accession # GSE178265)

### 16. Statistics

Group statistical analyses were performed using GraphPad Prism 10.1 software (GraphPad, La Jolla, CA). Sample size (n value) is defined by the number of observations (i.e., neurons, sections, or mice) and is indicated in the legends. All data are expressed as mean ± SEM or individual plots. Unless stated otherwise, statistical significance was determined by two-tailed Student’s t-tests for two-group comparisons. For multiple group comparisons, one-way or two-way analysis of variance (ANOVA) tests were used for normally distributed data, followed by post-hoc analyses. All statistical tests are specified in the figure legends.

## Resources table

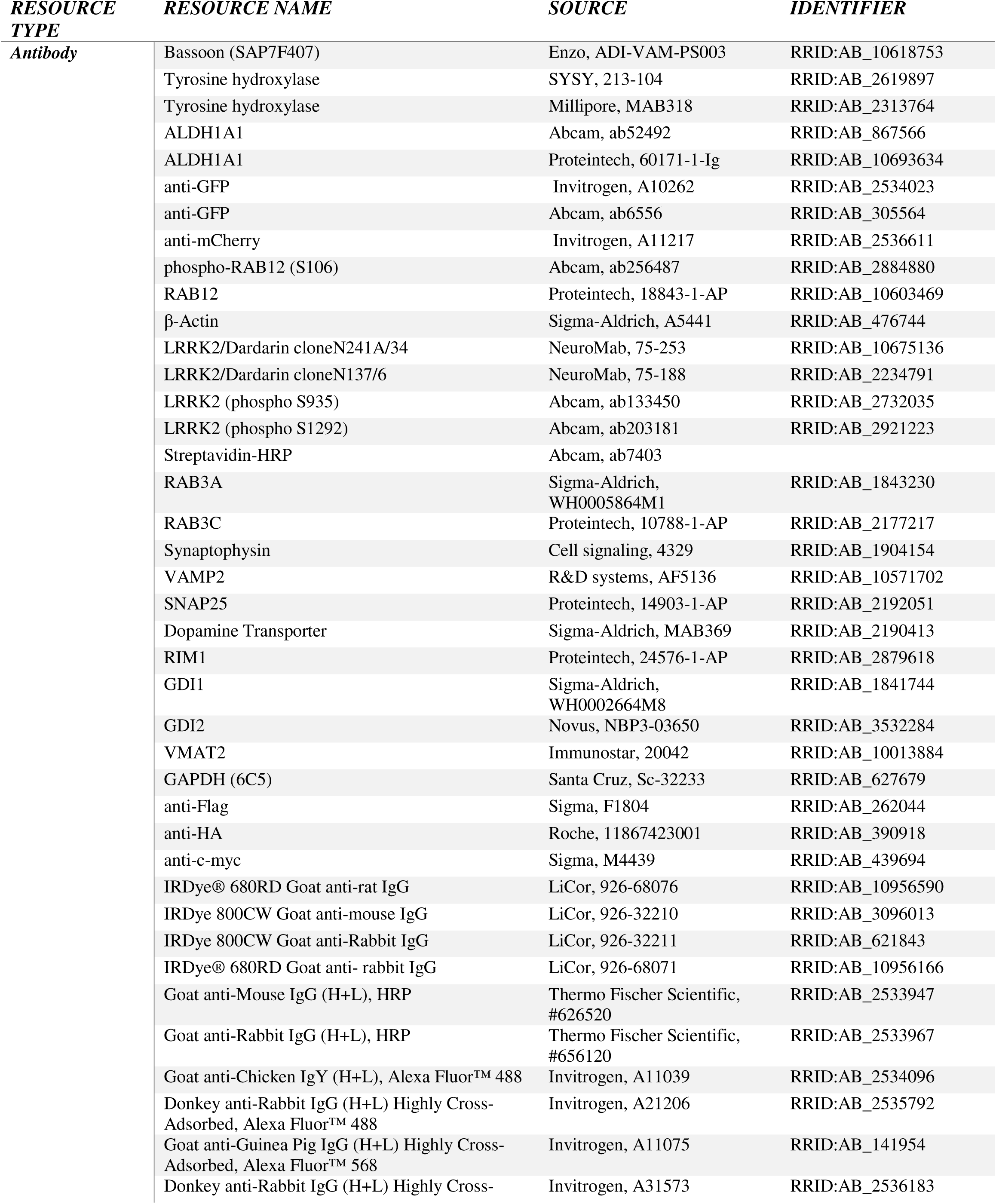
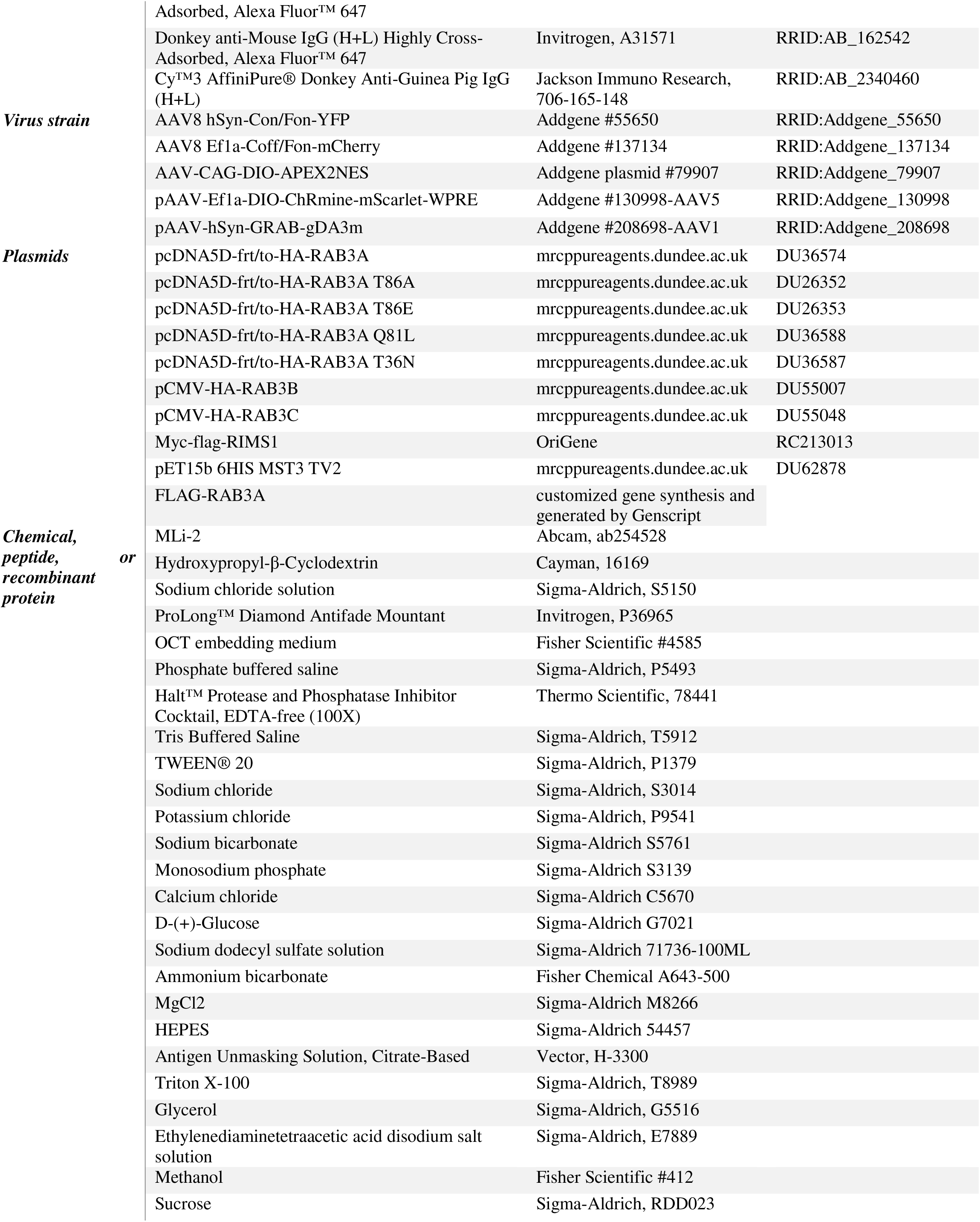
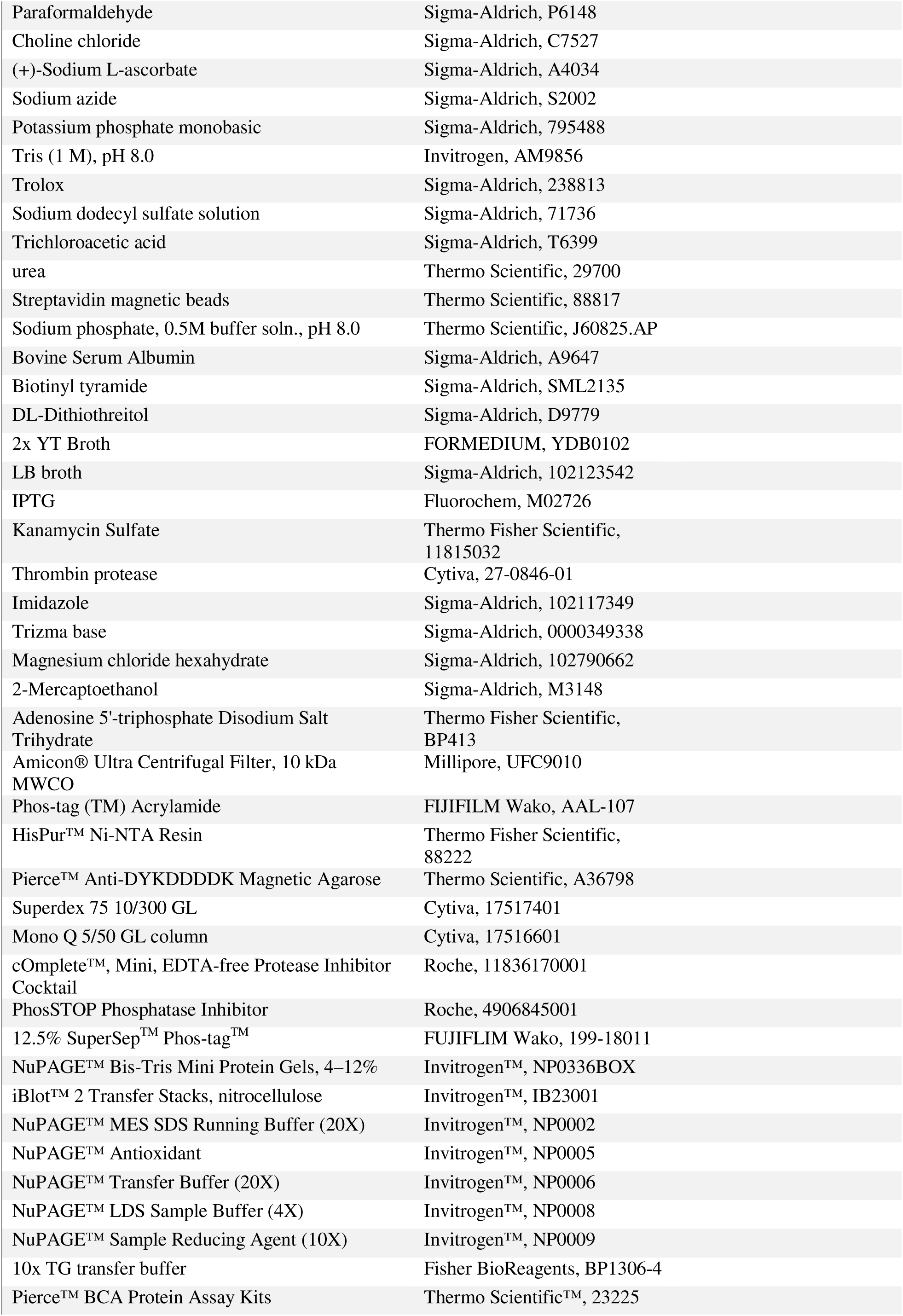
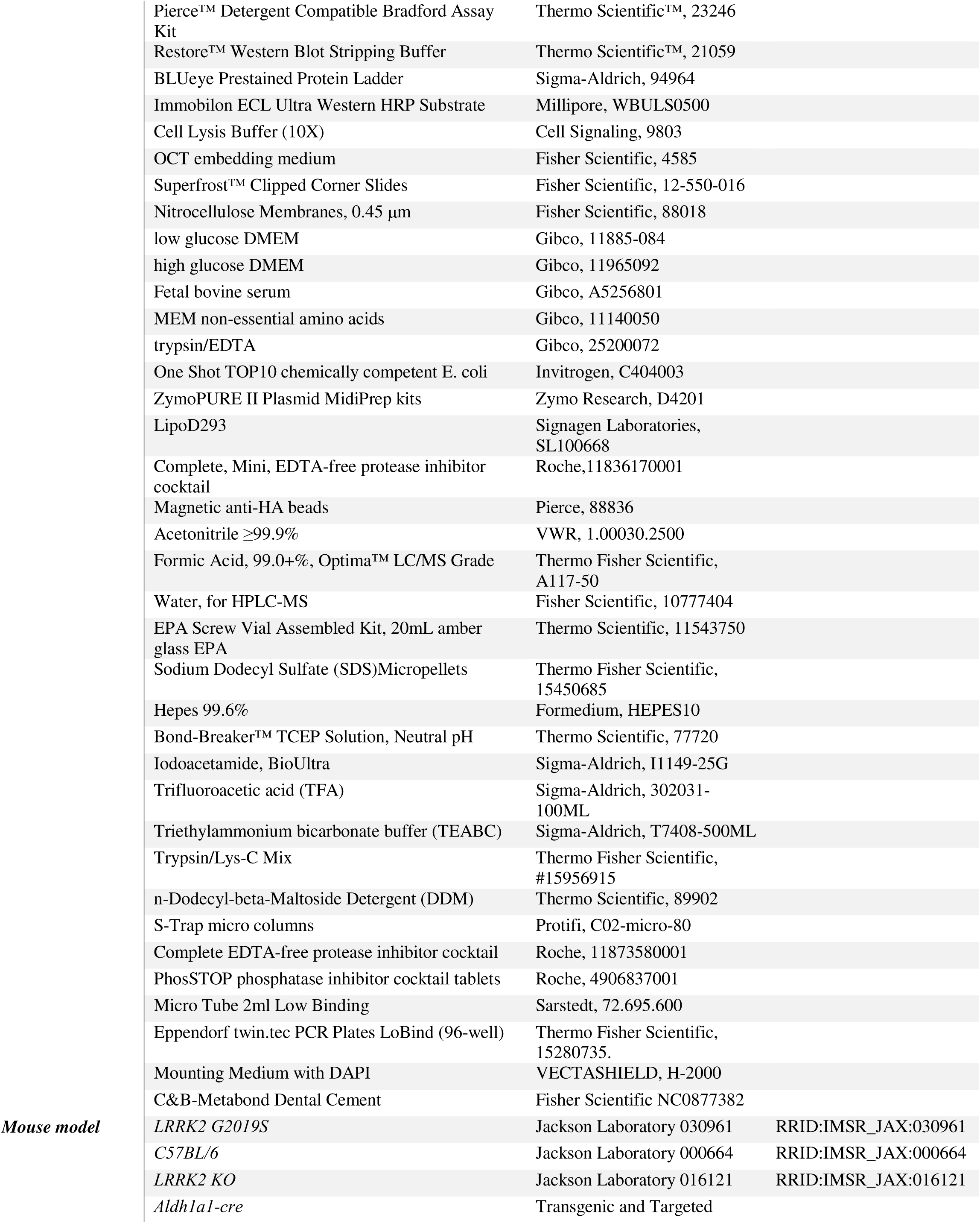
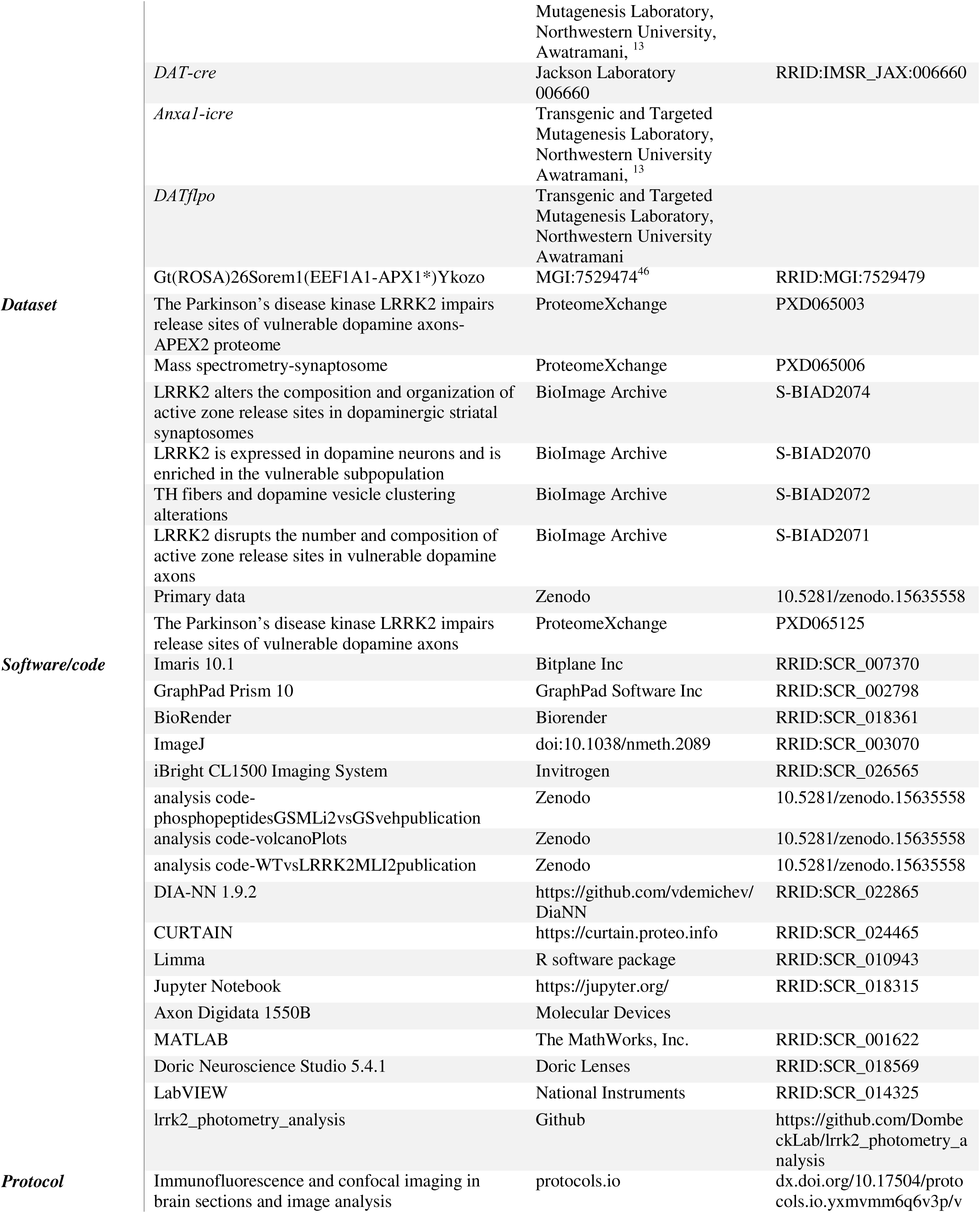
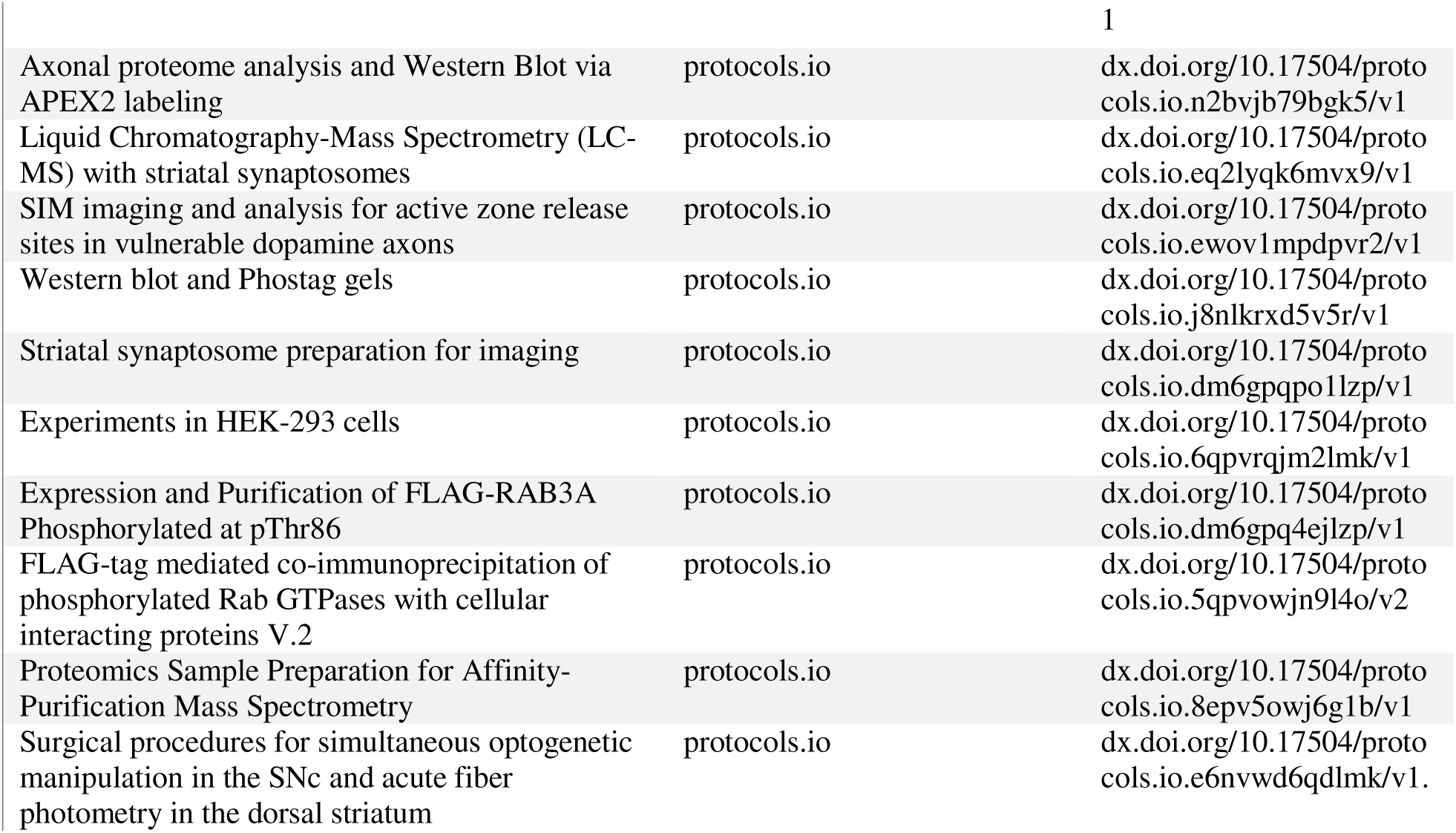

---

**Supplementary Figure 1.**
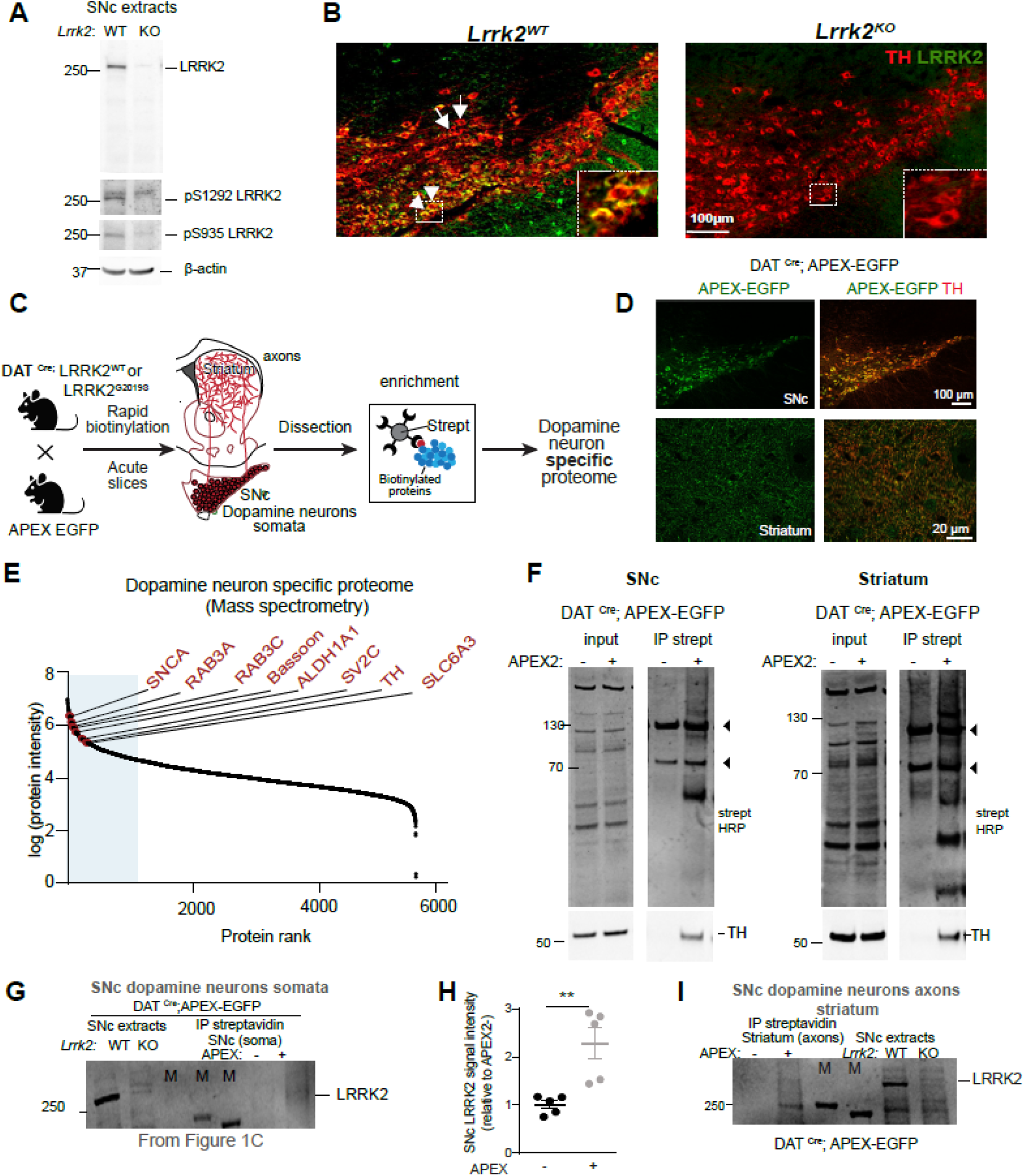
LRRK2 protein is expressed in SNc dopamine neurons (linked to Figure 1). **A.** WB analysis of SNc brain extracts from *Lrrk2^WT^* and *Lrrk2^KO^*mice probed with the antibodies indicated on the right. Note the absence of the LRRK2 signal in the *Lrrk2^KO^* tissues. **B.** Brain sections from *Lrrk2^WT^* and *Lrrk2^KO^* mice were stained with the LRRK2 antibody from A, along with TH. **C.** Workflow of APEX2-based proximity labeling within genetically labeled dopamine neurons in the mouse brain. Acute brain slices from DAT^Cre^; LRRK2^WT^ and DAT^Cre^; LRRK2^G2019S^ mice crossed with the APEX2 EGFP reporter mouse line^46^ were rapidly biotinylated, quenched, and dissected. Tissues were lysed and precipitated to remove free biotin. Tissue proteins were purified using streptavidin beads to enrich biotinylated proteins. On-bead digestion yielded peptides enriched from dopamine neurons, which were analyzed using mass spectrometry. **D.** Representative images showing the Cre-dependent expression of EGFP reporter along with TH immunostaining in the SNc and striatum of a DAT^Cre^; LRRK2^WT^; APEX EGFP mouse. **E.** 85% of the top 55 mDA neuron marker genes (e.g., TH and DAT; SLC6A3) from a publicly available APEX2 proteome dataset (dataset identifier PXD026229 ProteomeXchange Consortium) were detected in all our mass spectrometry samples from DAT^Cre^; APEX EGFP mice. 40% of these markers were among the top 20% (1105/5525) most highly expressed proteins by intensity (shaded light blue). **F.** WB analysis of biotinylated protein inputs and their streptavidin pulldowns from dissections of the indicated brain regions probed with antibodies as indicated on the right. Input: 10 μg of total protein (∼1% of striatal lysate and ∼5% of SNc); an equal volume of eluted proteins was loaded in each lane. **G**. Full gel from experiment presented in Figure 1C showing SNc extracts from *Lrrk2^WT^*and *Lrrk2^KO^* mice for reference of LRRK2 protein size. **H.** Quantification of LRRK2 intensity in DAT^Cre^ (APEX-) and DAT^Cre^; APEX EGFP mice (APEX+) was normalized to the APEX2-condition. Each dot represents an independent experiment with 3–4 mice pooled. **, p < 0.01, unpaired t-test. **I.** WB showing streptavidin pull-downs from equal volumes of striatal proteins from DAT^Cre^ or DAT^Cre^; APEX EGFP mice or DAT^Cre^ controls; SNc lysates from Lrrk2WT and Lrrk2KO are used as references for LRRK2 protein size. M, lanes loaded with molecular size markers. strept=streptavidin.

**Supplementary Figure 2.**
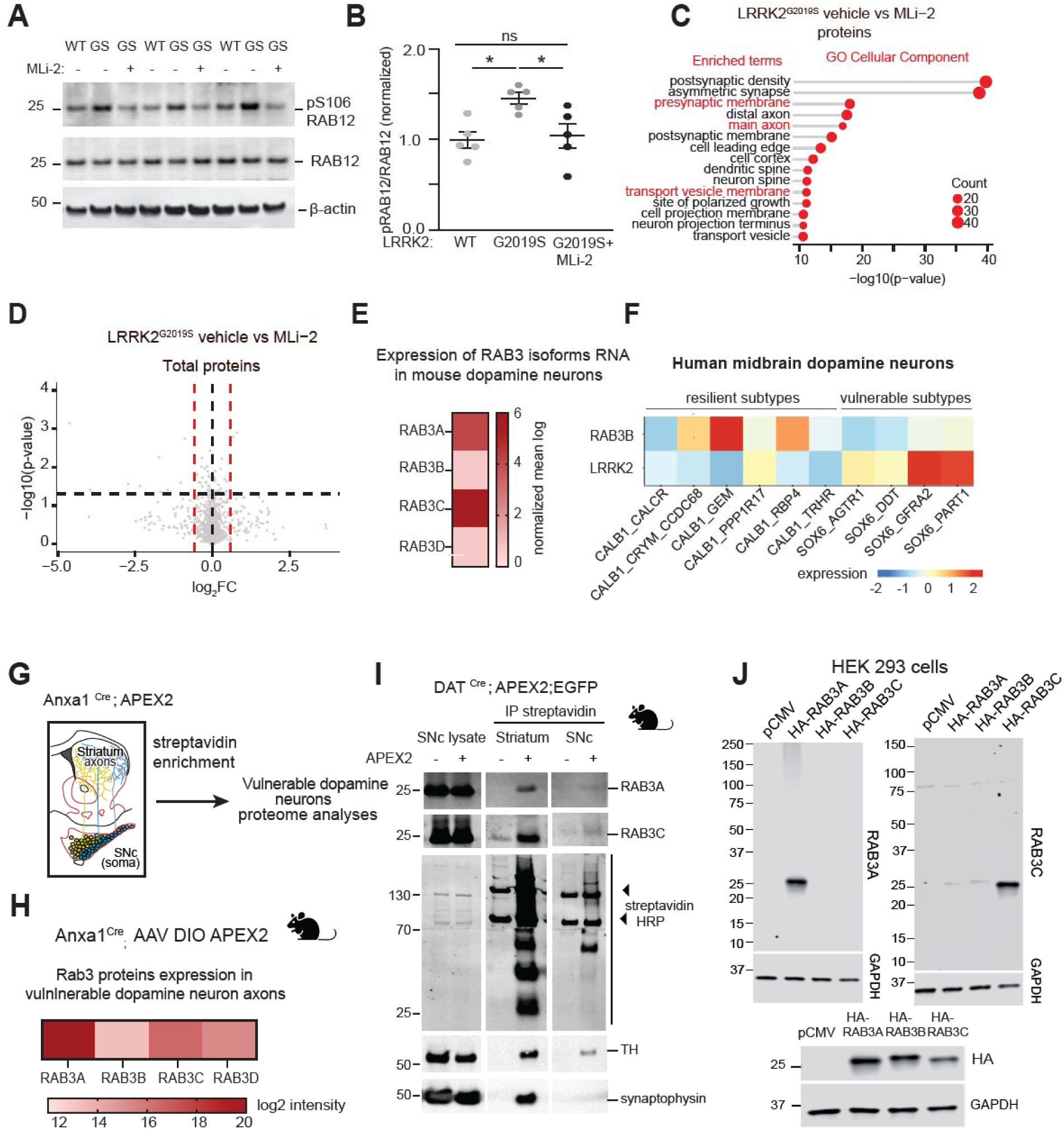
Heterogeneous RAB3 isoform expression across dopamine neuron subtypes (linked to Figure 2). **A**. WB analysis of SNc extracts from mice previously treated with MLi-2 (10 mg/kg) or vehicle for 2 h and probed for pS106 RAB12 (LRRK2 kinase target), total RAB12, and β-actin. **B.** Quantification of p-RAB12 band intensities normalized to total RAB12—n=5 mice/condition. Data are represented as mean±SEM (error bars). Asterisks denote statistical significance for Šídák’s multiple-comparison tests after a two-way ANOVA. *p< 0.05. (Treatment factor F(1,12)=8.614 p=0.0125, Genotype factor F(1, 12)=10.64, p=0.0068) **C.** Gene Ontology analysis of proteins with at least one differentially regulated phosphopeptide in striatal synaptosomes from vehicle vs MLi2-treated LRRK2^G2019S^ mice. The top 15 pathways significantly enriched in the cellular component term analysis (adjusted *p*-value_≤_0.05 by hypergeometric test with HB correction). All enriched pathways have been uploaded to Zenodo. Link in the resources table.**D.** Volcano plot comparing the significantly altered total proteins (|Log2FC|_>_0.58 and unadjusted *p*-value_≤_0.05 by multiple unpaired t-tests) between vehicle and MLi-2-treated LRRK2^G2019S^ striatal synaptosomes. **E**. Heat map of the expression of RAB3 isoforms in a dataset of single-cell RNA sequencing.^9^ The differential expression of RAB3 isoforms in TH+ SNc neurons compared to the remaining non-TH SNc neurons is expressed as normalized mean log values (RAB3A = 4.11, RAB3B = 0.693, RAB3C = 5.59, and RAB3D = 0.693). **F.** Heat map illustrating the expression of LRRK2 and RAB3B isoforms in human dopamine neuron subpopulations from a single nucleus RNA sequencing dataset,^49^ plotted with Shinny Cell.^99^ **G.** Workflow of APEX2 experiment for dopamine neuron subpopulation-specific analysis with subcellular compartment resolution. Cre-dependent APEX2 expressing AAV (AAV5-CAG-DIO-APEX2-NES) was injected into the SNc of Anxa1^Cre^ LRRK2^WT^ mice for Anxa1+ dopamine neuron-specific APEX2 labeling. **H.** Relative expression of RAB3 protein isoforms in our Anxa1+ dopamine axon subcluster proteomic dataset (for further details see Supplementary Figure 6)**. I.** Equal amounts of eluted proteins from streptavidin pulldowns from the striatum and SNc from either a LRRK2^WT^ or a LRRK2^G2019S^ DAT^Cre^ mouse injected with APEX2 AAV**. J.** HEK293T cells were transiently transfected with either pCMV or HA-tagged RAB3 isoforms and blotted with isoform-specific antibodies or anti-HA, and with GAPDH for loading control. Equal expression of all isoforms is shown in the panel below.

**Supplementary Figure 3.**
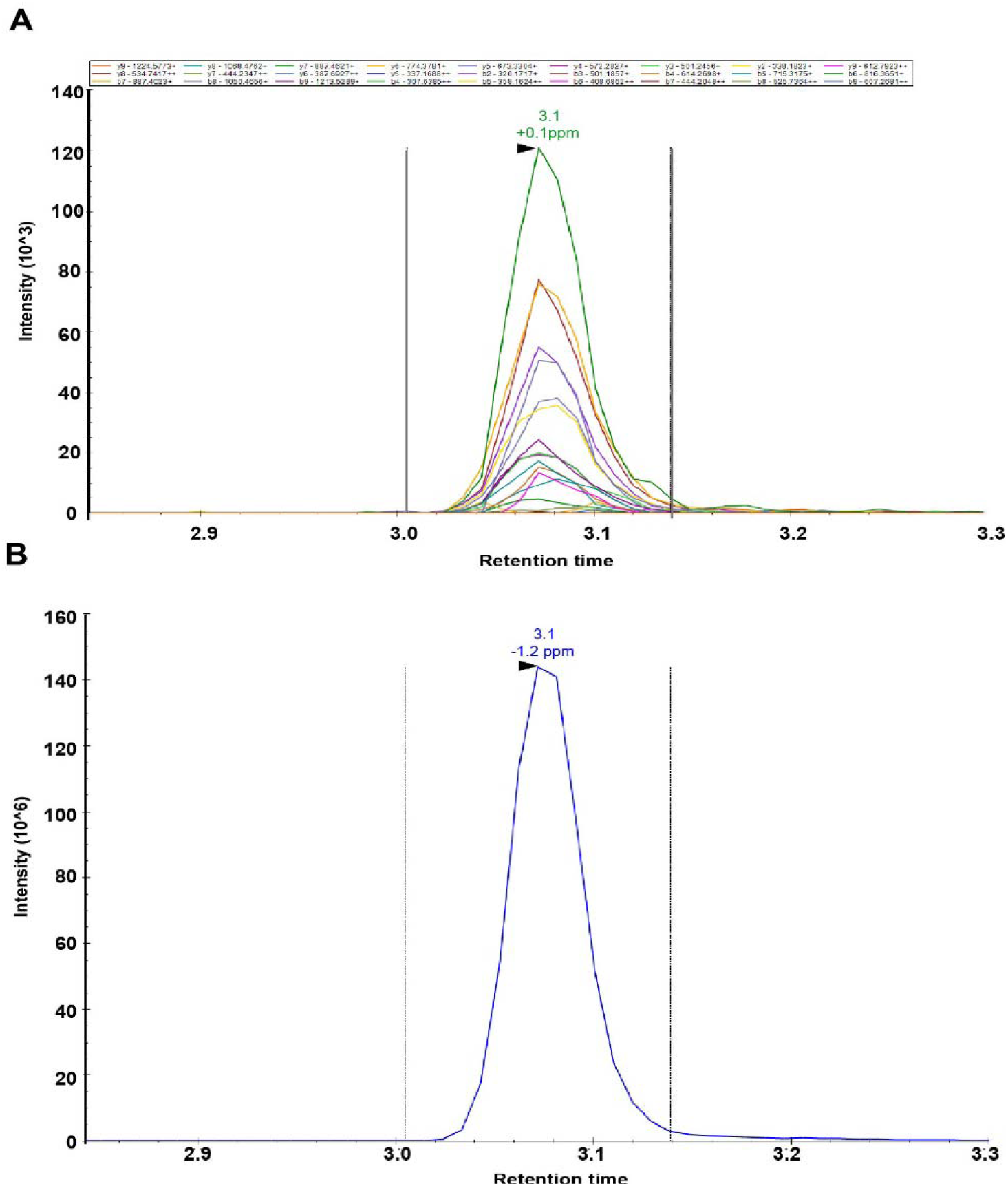
Differential interactome of p-RAB3A and unphosphorylated RAB3A in extracts from mouse brain (linked to Figure 3). **A.** Peptide fragment ions of the phosphopeptide; YR<u>T</u>ITTAYR, of the 2+ precursor ion **(B).** which spans phosphorylation site T86 in recombinant p-Flag-RAB3A protein. All chromatograms elute at the same retention time, and peptide identifications are at a false-discovery rate of <1% (DIA-NN 1.9.2).

**Supplementary Figure 4.**
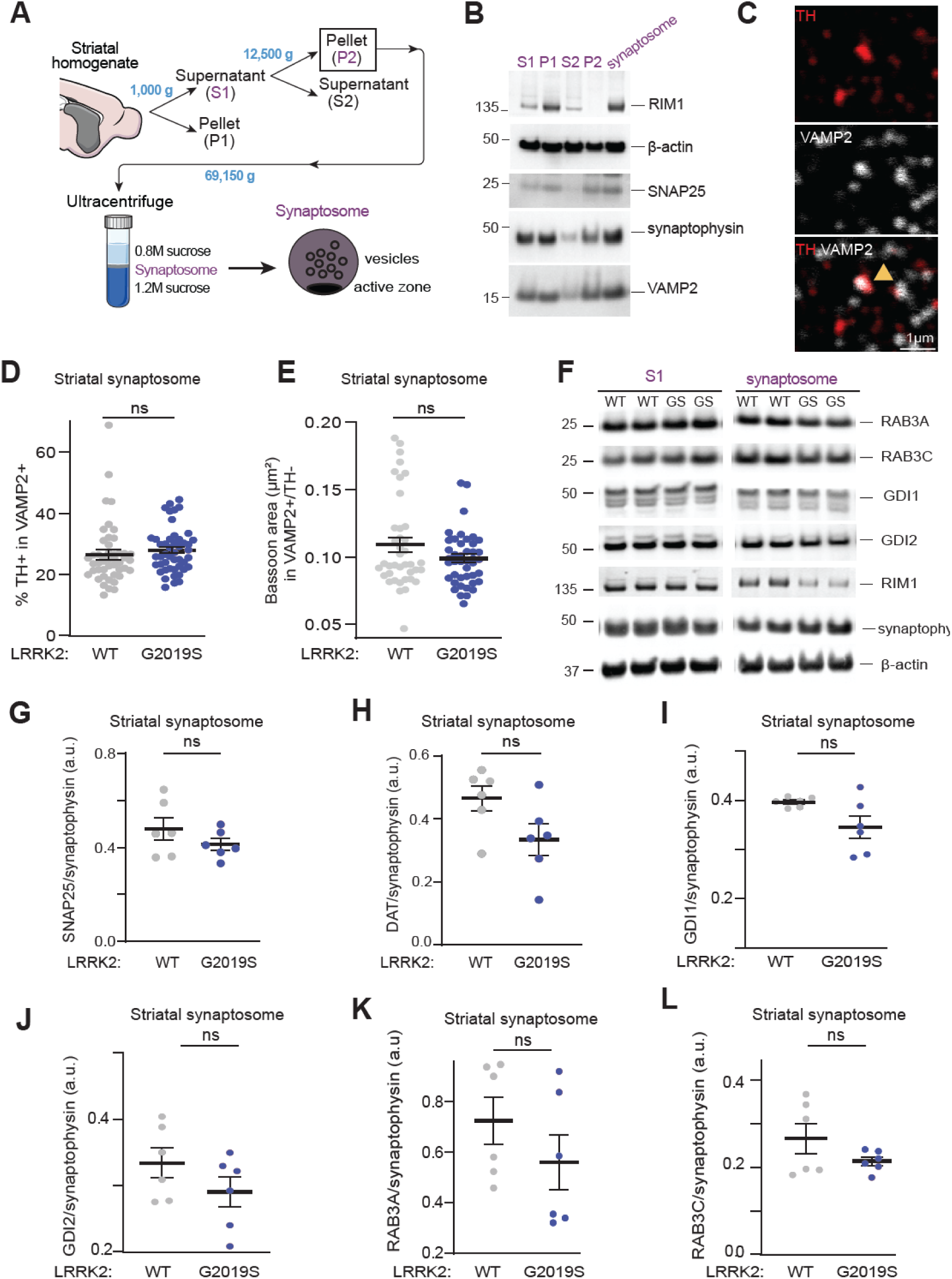
LRRK2 alters the distribution and organization of active zones in striatal synaptosomes (linked to Figure 4). **A.** Schematic representation of the steps for striatal synaptosome fraction preparation**. B.** WB analysis of selective fractions from striatal subcellular fractionation, probed with antibodies as indicated on the right. This analysis shows enrichment of synaptic vesicles and active zone proteins. **C.** Confocal image of striatal synaptosomes stained with TH and VAMP2. A double positive synaptosome is indicated with the arrowhead. **D.** Quantification of the percentage of TH+/VAMP2+ striatal synaptosomes in LRRK2^WT^ and LRRK2^G2019S^ mice (denoted by WT and GS, respectively). Each circle represents the average result for an area containing approximately 1,000 synaptosomes. n = 10-14 areas/4 mice per genotype. **E**. Quantification of bassoon area in VAMP2+ but TH-synaptosomes. Each circle represents the average of an image with ∼1,000 synaptosomes, n=10-14 areas/4 mice per genotype. Data in D and E represent mean± SEM. Ns denotes no statistical significance after unpaired t-tests. **F.** WB analysis of S1 and synaptosome fraction from LRRK2^WT^ and LRRK2^G2019S^ mice using the antibodies shown on the right. **G-L**. Quantification of the indicated protein signals in striatal synaptosomes from LRRK2^WT^ and LRRK2^G2019S^. Data are mean± SEM. Ns =non-significant differences in comparisons after unpaired t-tests. n= 6 mice/genotype. Equal amounts of proteins were loaded in all experiments, and duplicates represent biological replicates.

**Supplementary Figure 5.**
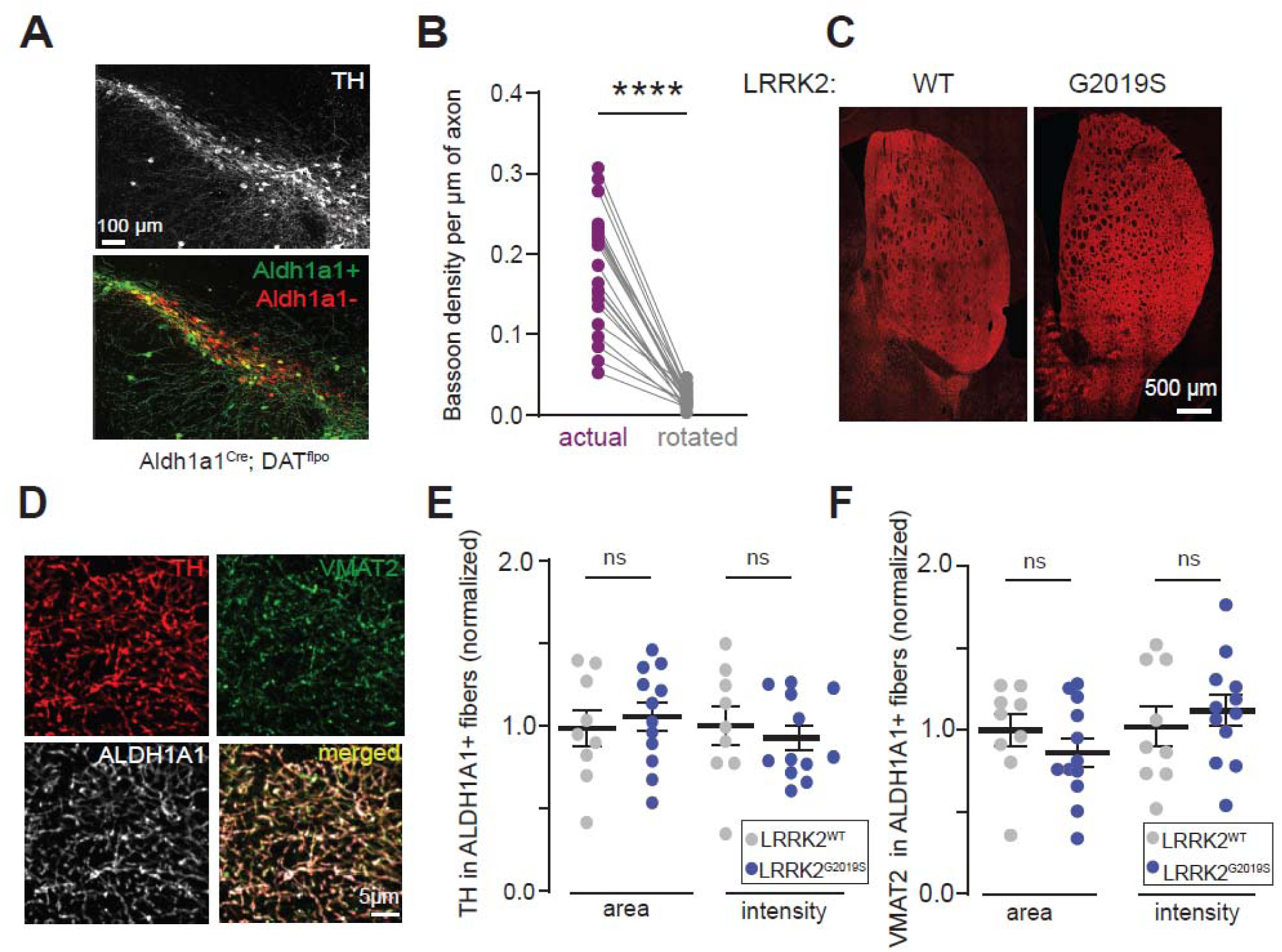
LRRK2^G2019S^ changes the number and composition of active zones without affecting the overall integrity of TH fibers (linked to Figure 5). **A.** SNc sections from Aldh1a1^Cre^; DAT^flpo^ mice injected with viral plasmids stained with EGFP and mCherry to label Aldh1a1+ and Aldh1a1-dopamine neurons, respectively. **B**. Image with bassoon signal in control mice was rotated by 180°, whereas the eGFP channel remained unaltered. Quantification of bassoon density within dopamine axons before and after image rotation. Each circle represents the average result of a region containing 25,000-35,000 bassoon clusters. Data represent mean ±SEM. n=18 (3-6 sections/mouse, 4 mice). Significant differences in comparisons after unpaired t-tests. **** p<0.0001. **C.** Representative low magnification striatal sections from 6-month-old LRRK2^WT^ and LRRK2^G2019S^ mice stained with TH. **D**. Representative high magnification dorsal striatal sections from adult (6 months) LRRK2^WT^ and LRRK2^G2019S^ mice stained with TH, ALDH1A1, and VMAT2 antibodies. **E**. Quantification of TH area and fluorescence intensity in ALDH1A1 immunostained dopamine axons across genotypes in the dorsal striatum. **F.** Average VMAT2 area and intensity in TH+/ALDH1A1+ dopamine axons across genotypes in dorsal striatum. For E and F, each dot represents the mean area or intensity of a striatal section, and data show mean±SEM; n=9-12 (2-3sections/mouse, 4-5mice/group). Ns: not significant with an unpaired t-test.

**Supplementary Figure 6.**
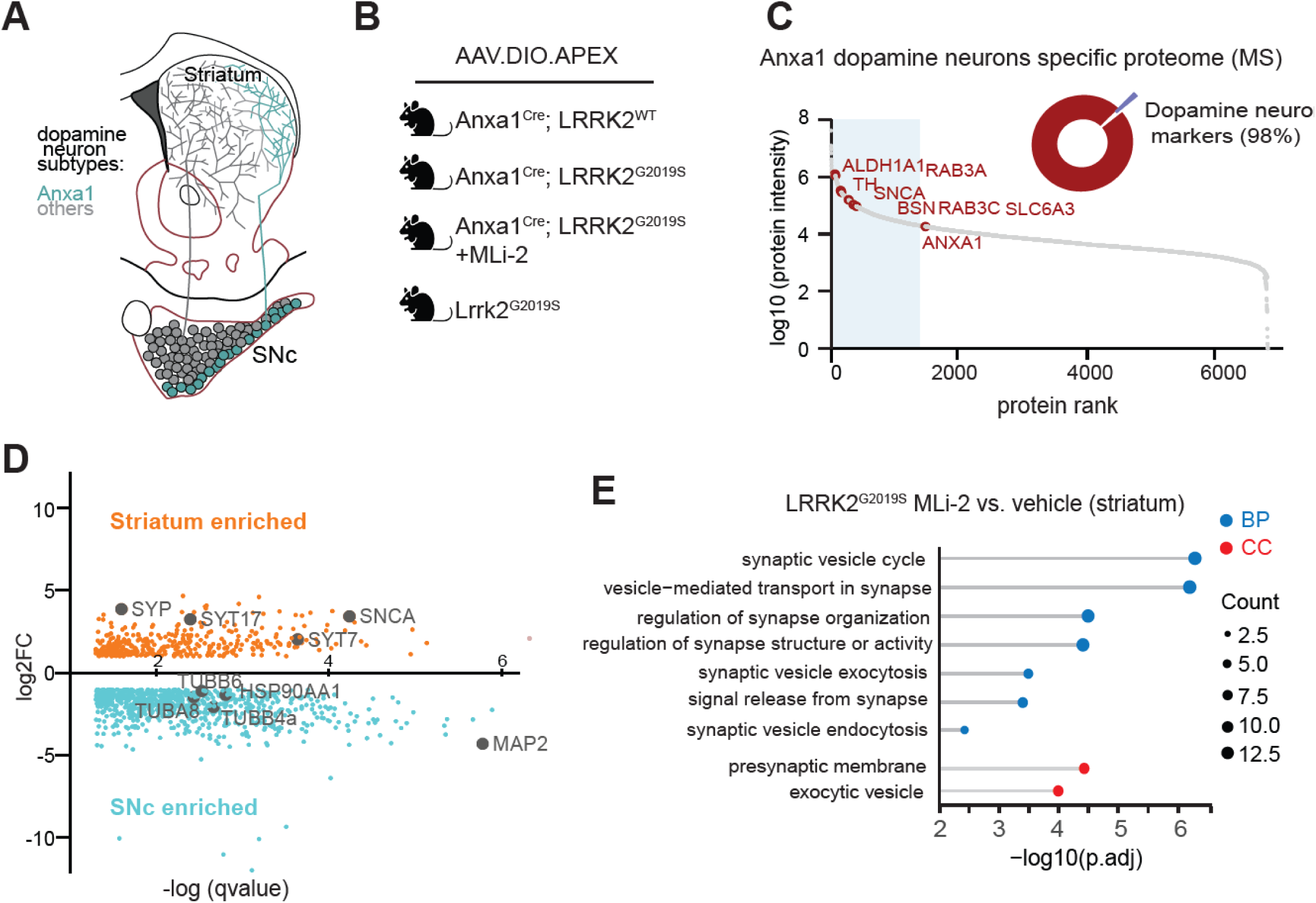
Axonal and somatic proteomics of Anxa1+ neurons (linked to Figure 5). **A.** Schematic illustrating the localization of Anxa1 dopamine neuron subtype in the mouse SNc and their projection patterns in the striatum. **B.** Experiment groups of APEX2-based proximity labeling within virally APEX2-labeled dopamine neurons in the mouse brain. **C**. 98% of the top 55 mDA neuron marker genes (e.g., TH and DAT (SLC6A3)) from a publicly available APEX2 proteome dataset (dataset identifier PXD026229 ProteomeXchange Consortium) were detected in all our mass spectrometry samples. **D.** Differential expression comparison of SNc and Striatum APEX+ streptavidin pulldown samples. Proteins colored orange or blue indicated significant enrichment in the Striatum vs. the SNc, respectively. (|Log2FC|_>_0.58 and unadjusted p-value_≤_0.05 by multiple unpaired t-tests) **E.** Gene Ontology analysis of proteins differentially expressed in striatum from vehicle vs. MLi2-treated Anxa1iCre LRRK2^G2019S^ mice. BP (Biological Process) and CC (Cellular Component) refer to categories in GO. All enriched pathways have been uploaded to Zenodo; the link is available in the key resources table.

**Supplementary Figure 7.**
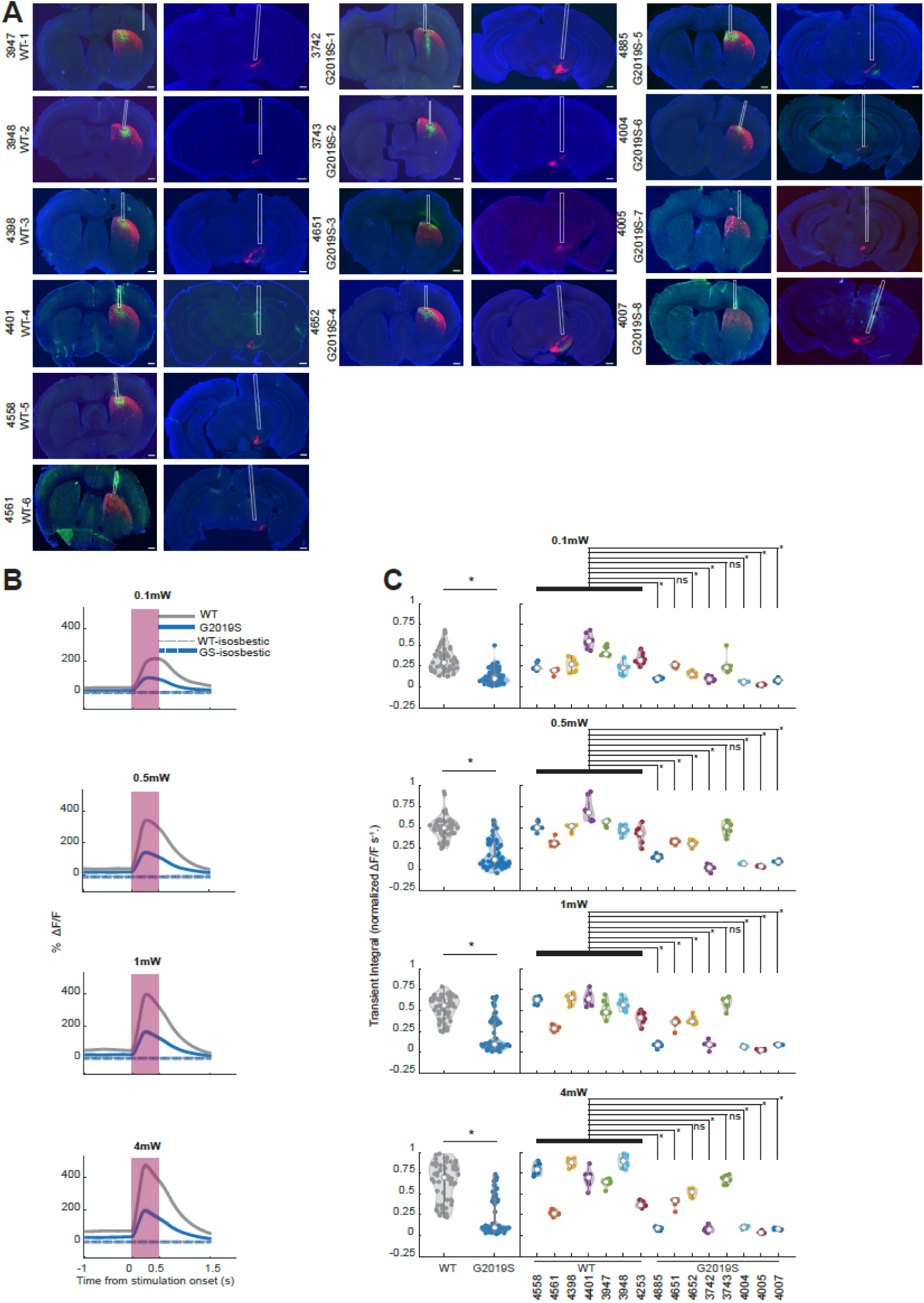
Histology and power dependence for in vivo evoked dopamine measurements (linked to Figure 6). **A.** Histological verification of viral expression and fiber placement in each animal. Left two columns, coronal sections of the dorsal striatum region (dStr) and the Substantia Nigra pars compacta (SNc) of each LRRK2^WT^ mouse used in the analysis. Red fluorescence indicates the expression of ChRmine in the somas (SNc) and the axons (dStr) of Anxa1+ dopaminergic neurons. Green fluorescence indicates the expression of GRAB-DA3m in the dorsal striatum. White rectangles indicate where the photometry (dStr) and optogenetics (SNc) fibers were placed. Right four columns, same as left, but in LRRK2^G2019S^ mice. Scale bar = 0.5 mm. **B.** Same as Figure 6C but with 0.1 mW, 0.5 mW, 1 mW, and 4 mW optogenetic light powers from top row to bottom row (blue trace, LRRK2^G2019S^ mice = 8, n = 64 stimulations; grey trace, LRRK2^WT^ mice = 7, n = 56 stimulations). Shaded regions denote mean_±_SEM across stimulations. **C.** Same as Figure 6D but with 0.1 mW, 0.5 mW, 1 mW, and 4 mW optogenetic light powers from top row to bottom row. White dots denote group means and black bars denote 25th to 75th percentile. Left, group comparisons at each power (LRRK2^G2019S^ n = 64 stimulations per power, LRRK2^WT^ n = 56 stimulations per power; p-value (0.1 mW) = 2.59E-15, p-value (0.5 mW) = 2.20E-20, p-value (1 mW) = 2.45E-17, p-value (4 mW) = 4.78E-16, unpaired t-test). Right, dopamine release response in individual mice. Numbers denote mouse ID. Asterisks (*) on the right denote p-value < 0.00625 from Welch’s t-test with Bonferroni correction (α = 0.05/8 = 0.00625) comparing individual LRRK2^G2019S^ mouse to the LRRK2^WT^ group.

**Supplementary Figure 8.**
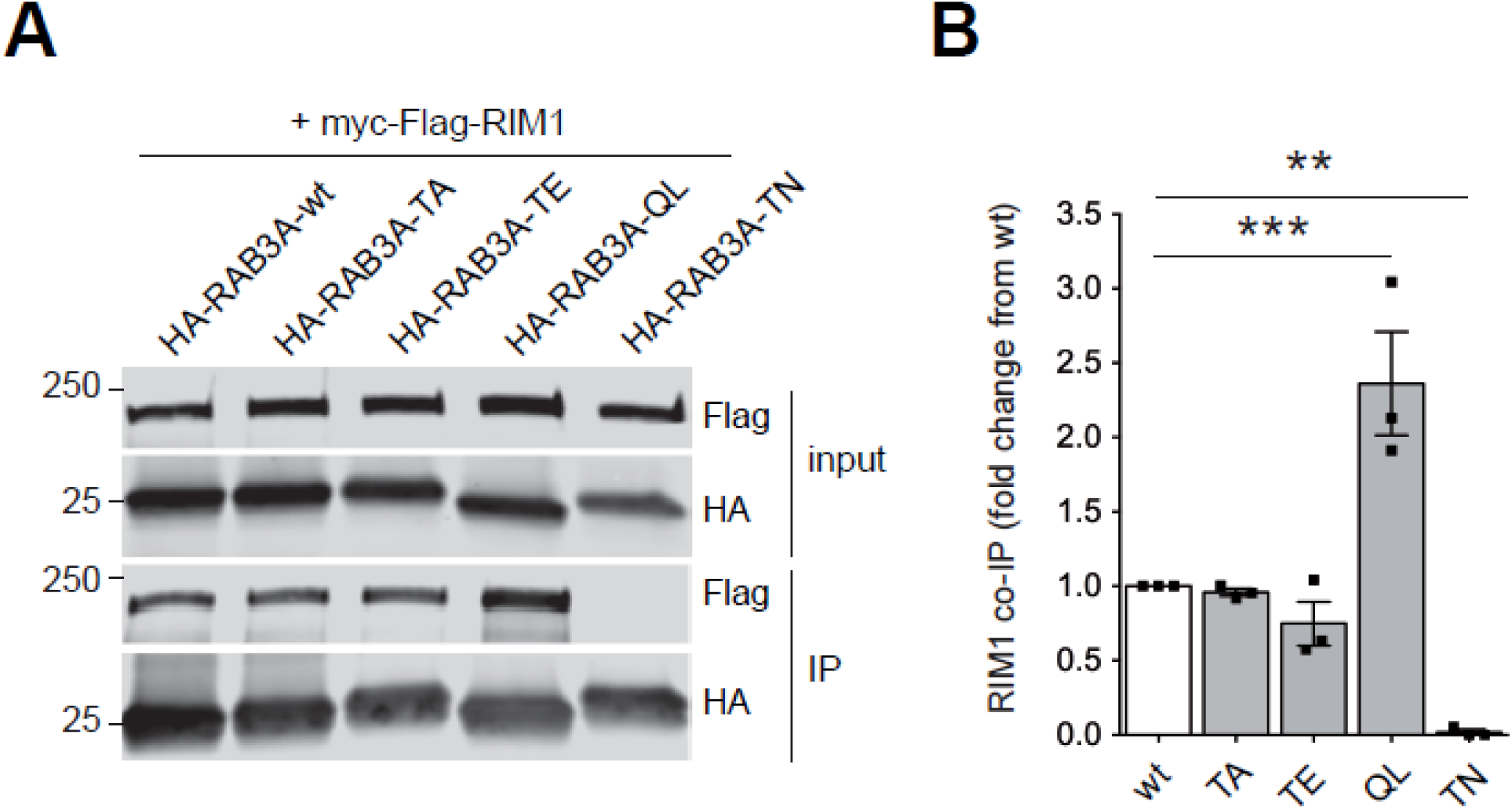
Phosphomimetic RAB3 constructs fail to reproduce the effects of phosphorylated RAB3 proteins (linked to Figure 3). **A.** HEK293 cells were transiently co-transfected with HA-tagged RAB3A constructs and myc-flag-RIM1. Extracts were subjected to pulldown with anti-HA beads, and input and immunoprecipitated material (IP) subjected to blotting with the indicated antibodies. WT, phospho-deficient (RAB3A-T86A) and phospho-mimic (RAB3A-T86E) RAB3A show similar binding to tagged RIM1. In contrast, and as expected, the RAB3A mutant mimicking the GTP-bound state (Q81L) shows increased RIM1 binding, whilst the RAB3A mutant mimicking the GDP-bound state (T36N) shows loss of RIM1 binding. **B.** Quantification of experiments of the type shown in A. Co-immunoprecipitation of myc-flag-RIM1 with the indicated HA-tagged RAB3A constructs was normalized to that obtained with wild-type (wt) RAB3A. n=3 independent experiments (mean±SEM). Asterisk indicates statistical significance as determined by one-way ANOVA with Dunnett’s multiple comparisons. ***p<0.001; **p<0.0.

## References

(1) Damier, P.; Hirsch, E. C.; Agid, Y.; Graybiel, A. M. The Substantia Nigra of the Human Brain: II. Patterns of Loss of Dopamine-Containing Neurons in Parkinson’s Disease. Brain 1999, 122 (8), 1437–1448. 10.1093/brain/122.8.1437.

(2) Fearnley, J. M.; Lees, A. J. Ageing and Parkinson’s Disease: Substantia Nigra Regional Selectivity. Brain 1991, 114 (5), 2283–2301. 10.1093/brain/114.5.2283.

(3) Sulzer, D.; Surmeier, D. J. Neuronal Vulnerability, Pathogenesis, and Parkinson’s Disease. Movement Disorders. January 2013, pp 41–50. 10.1002/mds.25095.

(4) Kish, S. J.; Shannak, K.; Hornykiewicz, O. Uneven Pattern of Dopamine Loss in the Striatum of Patients with Idiopathic Parkinson’s Disease. N. Engl. J. Med. 1988, 318, 876–880.

(5) Fu, Y. H.; Paxinos, G.; Watson, C.; Halliday, G. M. The Substantia Nigra and Ventral Tegmental Dopaminergic Neurons from Development to Degeneration. J. Chem. Neuroanat. 2016, 76, 98–107. 10.1016/j.jchemneu.2016.02.001.

(6) Gibb, W. R. G.; Lees, A. J. Anatomy, Pigmentation, Ventral and Dorsal Subpopulations of the Substantia Nigra, and Differential Cell Death in Parkinson’s Disease. J. Neurol. Neurosurg. Psychiatry 1991, 54 (5), 388–396. 10.1136/jnnp.54.5.388.

(7) Poulin, J. F.; Zou, J.; Drouin-Ouellet, J.; Kim, K. Y. A.; Cicchetti, F.; Awatramani, R. B. Defining Midbrain Dopaminergic Neuron Diversity by Single-Cell Gene Expression Profiling. Cell Rep. 2014, 9 (3), 930–943. 10.1016/j.celrep.2014.10.008.

(8) La Manno, G.; Gyllborg, D.; Codeluppi, S.; Nishimura, K.; Salto, C.; Zeisel, A.; Borm, L. E.; Stott, S. R. W.; Toledo, E. M.; Villaescusa, J. C.; Lönnerberg, P.; Ryge, J.; Barker, R. A.; Arenas, E.; Linnarsson, S. Molecular Diversity of Midbrain Development in Mouse, Human, and Stem Cells. Cell 2016, 167 (2), 566–580.e19. 10.1016/j.cell.2016.09.027.

(9) Saunders, A.; Macosko, E. Z.; Wysoker, A.; Goldman, M.; Krienen, F. M.; de Rivera, H.; Bien, E.; Baum, M.; Bortolin, L.; Wang, S.; Goeva, A.; Nemesh, J.; Kamitaki, N.; Brumbaugh, S.; Kulp, D.; McCarroll, S. A. Molecular Diversity and Specializations among the Cells of the Adult Mouse Brain. Cell 2018, 174 (4), 1015–1030.e16. 10.1016/j.cell.2018.07.028.

(10) Wu, J.; Kung, J.; Dong, J.; Chang, L.; Xie, C.; Habib, A.; Hawes, S.; Yang, N.; Chen, V.; Liu, Z.; Evans, R.; Liang, B.; Sun, L.; Ding, J.; Yu, J.; Saez-Atienzar, S.; Tang, B.; Khaliq, Z.; Lin, D. T.; Le, W.; Cai, H. Distinct Connectivity and Functionality of Aldehyde Dehydrogenase 1a1-Positive Nigrostriatal Dopaminergic Neurons in Motor Learning. Cell Rep. 2019, 28 (5), 1167–1181.e7. 10.1016/j.celrep.2019.06.095.

(11) Liu, G.; Yu, J.; Ding, J.; Xie, C.; Sun, L.; Rudenko, I.; Zheng, W.; Sastry, N.; Luo, J.; Rudow, G.; Troncoso, J. C.; Cai, H. Aldehyde Dehydrogenase 1 Defines and Protects a Nigrostriatal Dopaminergic Neuron Subpopulation. Journal of Clinical Investigation 2014, 124 (7), 3032–3046. 10.1172/JCI72176.

(12) Pereira Luppi, M.; Azcorra, M.; Caronia-Brown, G.; Poulin, J. F.; Gaertner, Z.; Gatica, S.; Moreno-Ramos, O. A.; Nouri, N.; Dubois, M.; Ma, Y. C.; Ramakrishnan, C.; Fenno, L.; Kim, Y. S.; Deisseroth, K.; Cicchetti, F.; Dombeck, D. A.; Awatramani, R. Sox6 Expression Distinguishes Dorsally and Ventrally Biased Dopamine Neurons in the Substantia Nigra with Distinctive Properties and Embryonic Origins. Cell Rep. 2021, 37 (6). 10.1016/j.celrep.2021.109975.

(13) Azcorra, M.; Gaertner, Z.; Davidson, C.; He, Q.; Kim, H.; Nagappan, S.; Hayes, C. K.; Ramakrishnan, C.; Fenno, L.; Kim, Y. S.; Deisseroth, K.; Longnecker, R.; Awatramani, R.; Dombeck, D. A. Unique Functional Responses Differentially Map onto Genetic Subtypes of Dopamine Neurons. Nat. Neurosci. 2023. 10.1038/s41593-023-01401-9.

(14) Mantas, I.; Contestabile, A.; Skara, V.; Loiseau, C.; Santos, I. A.; Cramb, K. M. L.; Filograna, R.; Wade-Martins, R.; Magill, P.; Meletis, K. Anxa1+ Dopamine Neuron Vulnerability Defines Prodromal Parkinson’s Disease Bradykinesia and Procedural Motor Learning Impairment. December 22, 2024. 10.1101/2024.12.22.629963.

(15) Fushiki, A.; Ng, D.; Lewis, Z. R.; Yadav, A.; Saraiva, T.; Hammand, L. A.; Wirblich, C.; Tasic, B.; Menon, V.; da Silva, J. A.; Costa, R. M. A Vulnerable Subtype of Dopaminergic Neurons Drives Early Motor Deficits in Parkinson’s Disease. bioRxiv 2024. 10.1101/2024.12.20.629776.

(16) Hadjas, L. C.; Kollman, G. J.; Linderhof, L.; Xia, M.; Mansur, S.; Saint-Pierre, M.; Lim, B. K.; Lee, E. B.; Cicchetti, F.; Awatramani, R.; Hollon, N. G.; Hnasko, T. S. Parkinson’s Disease-Vulnerable and -Resilient Dopamine Neurons Display Opposite Responses to Excitatory Input. June 7, 2025. 10.1101/2025.06.03.657460.

(17) Poulin, J. F.; Caronia, G.; Hofer, C.; Cui, Q.; Helm, B.; Ramakrishnan, C.; Chan, C. S.; Dombeck, D. A.; Deisseroth, K.; Awatramani, R. Mapping Projections of Molecularly Defined Dopamine Neuron Subtypes Using Intersectional Genetic Approaches. Nat. Neurosci. 2018, 21 (9), 1260–1271. 10.1038/s41593-018-0203-4.

(18) Surmeier, D. J.; Calabresi, P.; Volpicelli-Daley, L. A.; Gcwensa, N. Z.; Russell, D. L.; Cowell, R. M. Molecular Mechanisms Underlying Synaptic and Axon Degeneration in Parkinson’s Disease. 2021. 10.3389/fncel.2021.626128.

(19) Wong, Y. C.; Luk, K.; Purtell, K.; Burke Nanni, S.; Stoessl, A. J.; Trudeau, L. E.; Yue, Z.; Krainc, D.; Oertel, W.; Obeso, J. A.; Volpicelli-Daley, L. A. Neuronal Vulnerability in Parkinson Disease: Should the Focus Be on Axons and Synaptic Terminals? Movement Disorders. John Wiley and Sons Inc. 2019, pp 1406–1422. 10.1002/mds.27823.

(20) Kordower, J. H.; Olanow, C. W.; Dodiya, H. B.; Chu, Y.; Beach, T. G.; Adler, C. H.; Halliday, G. M.; Bartus, R. T. Disease Duration and the Integrity of the Nigrostriatal System in Parkinson’s Disease. Brain 2013, 136 (Pt 8), 2419–2431. 10.1093/brain/awt192.

(21) Cramb, K. M. L.; Beccano-Kelly, D.; Cragg, S. J.; Wade-3 Martins, R. Impaired Dopamine Release in Parkinson’s Disease 2 Here We Review the Evidence for Dopamine Release Deficits Prior to Neurodegeneration In. 2023. 10.1093/brain/awad064/7067886.

(22) Leonard, H. L. Novel Parkinson’s Disease Genetic Risk Factors Within and Across European Populations. March 17, 2025. 10.1101/2025.03.14.24319455.

(23) Chang, D.; Nalls, M. A.; Hallgrímsdóttir, I. B.; Hunkapiller, J.; Brug, M. van der; Cai, F.; Kerchner, G. A.; Ayalon, G.; Bingol, B.; Sheng, M.; Hinds, D.; Behrens, T. W.; Singleton, A. B.; Bhangale, T. R.; Graham, R. R. A Meta-Analysis of Genome-Wide Association Studies Identifies 17 New Parkinson’s Disease Risk Loci. Nat. Genet. 2017, 49 (10), 1511–1516. 10.1038/ng.3955.

(24) Krebs, C. E.; Karkheiran, S.; Powell, J. C.; Cao, M.; Makarov, V.; Darvish, H.; Di Paolo, G.; Walker, R. H.; Shahidi, G. A.; Buxbaum, J. D.; De Camilli, P.; Yue, Z.; Paisán-Ruiz, C. The Sac1 Domain of SYNJ1 Identified Mutated in a Family with Early-Onset Progressive Parkinsonism with Generalized Seizures. Hum. Mutat. 2013, 34 (9), 1200–1207. 10.1002/humu.22372.

(25) Olgiati, S.; Quadri, M.; Fang, M.; Rood, J. P. M. A.; Saute, J. A.; Chien, H. F.; Bouwkamp, C. G.; Graafland, J.; Minneboo, M.; Breedveld, G. J.; Zhang, J.; Verheijen, F. W.; Boon, A. J. W.; Kievit, A. J. A.; Jardim, L. B.; Mandemakers, W.; Barbosa, E. R.; Rieder, C. R. M.; Leenders, K. L.; Wang, J.; Bonifati, V. DNAJC6 Mutations Associated with Early-Onset Parkinson’s Disease. Ann. Neurol. 2016, 79 (2), 244–256. 10.1002/ana.24553.

(26) Cookson, M. R. The Role of Leucine-Rich Repeat Kinase 2 (LRRK2) in Parkinson’s Disease. Nat Rev Neurosci 2010, 11 (12), 791–797. 10.1038/nrn2935.

(27) Alessi, D. R.; Sammler, E. LRRK2 Kinase in Parkinson’s Disease. Science. American Association for the Advancement of Science 2018, pp 36–37. 10.1126/science.aar5683.

(28) Tokars, V.; Chen, C.; Parisiadou, L. Closing the Structure-to-Function Gap for LRRK2. Trends Biochem. Sci. 2022, 47 (3), 187–188. 10.1016/J.TIBS.2021.10.003.

(29) Di Maio, R.; Hoffman, E. K.; Rocha, E. M.; Keeney, M. T.; Sanders, L. H.; De Miranda, B. R.; Zharikov, A.; Van Laar, A.; Stepan, A. F.; Lanz, T. A.; Kofler, J. K.; Burton, E. A.; Alessi, D. R.; Hastings, T. G.; Timothy Greenamyre, J. LRRK2 Activation in Idiopathic Parkinson’s Disease. Sci. Transl. Med. 2018, 10 (451). 10.1126/scitranslmed.aar5429.

(30) Jennings, D.; Huntwork-Rodriguez, S.; Henry, A. G.; Sasaki, J. C.; Meisner, R.; Diaz, D.; Solanoy, H.; Wang, X.; Negrou, E.; Bondar, V. V; Ghosh, R.; Maloney, M. T.; Propson, N. E.; Zhu, Y.; Maciuca, R. D.; Harris, L.; Kay, A.; LeWitt, P.; Alex King, T.; Kern, D.; Ellenbogen, A.; Goodman, I.; Siderowf, A.; Aldred, J.; Omidvar, O.; Masoud, S. T.; Davis, S. S.; Arguello, A.; Estrada, A. A.; de Vicente, J.; Sweeney, Z. K.; Astarita, G.; Borin, M. T.; Wong, B. K.; Wong, H.; Nguyen, H.; Scearce-Levie, K.; Ho, C.; Troyer, M. D. Preclinical and Clinical Evaluation of the LRRK2 Inhibitor DNL201 for Parkinson’s Disease; 2022; Vol. 14. https://www.science.org.

(31) Jennings, D.; Huntwork-Rodriguez, S.; Vissers, M. F. J. M.; Daryani, V. M.; Diaz, D.; Goo, M. S.; Chen, J. J.; Maciuca, R.; Fraser, K.; Mabrouk, O. S.; van de Wetering de Rooij, J.; Heuberger, J. A. A. C.; Groeneveld, G. J.; Borin, M. T.; Cruz-Herranz, A.; Graham, D.; Scearce-Levie, K.; De Vicente, J.; Henry, A. G.; Chin, P.; Ho, C.; Troyer, M.D. LRRK2 Inhibition by BIIB122 in Healthy Participants and Patients with Parkinson’s Disease. Movement Disorders 2023, 38 (3), 386–398. 10.1002/mds.29297.

(32) Volta, M.; Melrose, H. LRRK2 Mouse Models: Dissecting the Behavior, Striatal Neurochemistry and Neurophysiology of PD Pathogenesis. Biochemical Society Transactions. Portland Press Ltd 2017, pp 113–122. 10.1042/BST20160238.

(33) Chang, E. E. S.; Ho, P. W. L.; Liu, H. F.; Pang, S. Y. Y.; Leung, C. T.; Malki, Y.; Choi, Z. Y. K.; Ramsden, D. B.; Ho, S. L. LRRK2 Mutant Knock-in Mouse Models: Therapeutic Relevance in Parkinson’s Disease. Translational Neurodegeneration. BioMed Central Ltd December 1, 2022. 10.1186/s40035-022-00285-2.

(34) Yue, M.; Hinkle, K. M.; Davies, P.; Trushina, E.; Fiesel, F. C.; Christenson, T. A.; Schroeder, A. S.; Zhang, L.; Bowles, E.; Behrouz, B.; Lincoln, S. J.; Beevers, J. E.; Milnerwood, A. J.; Kurti, A.; McLean, P. J.; Fryer, J. D.; Springer, W.; Dickson, D. W.; Farrer, M. J.; Melrose, H. L. Progressive Dopaminergic Alterations and Mitochondrial Abnormalities in LRRK2 G2019S Knock-in Mice. Neurobiol. Dis. 2015, 78, 172–195. 10.1016/j.nbd.2015.02.031.

(35) Xenias, H. S.; Chen, C.; Kang, S.; Cherian, S.; Situ, X.; Shanmugasundaram, B.; Liu, G.; Scesa, G.; Savio Chan, C.; Parisiadou, L. R 1441C and G2019S LRRK2 Knockin Mice Have Distinct Striatal Molecular, Physiological, and Behavioral Alterations. Commun. Biol. 2022, 5 (1). 10.1038/s42003-022-04136-8.

(36) Tong, Y.; Pisani, A.; Martella, G.; Karouani, M.; Yamaguchi, H.; Pothos, E. N.; Shen, J. R 1441C Mutation in LRRK2 Impairs Dopaminergic Neurotransmission in Mice. Proc Natl Acad Sci U S A 2009, 106 (34), 14622–14627. 10.1073/pnas.0906334106.

(37) Tozzi, A.; Tantucci, M.; Marchi, S.; Mazzocchetti, P.; Morari, M.; Pinton, P.; Mancini, A.; Calabresi, P. Dopamine D2 Receptor-Mediated Neuroprotection in a G2019S Lrrk2 Genetic Model of Parkinson’s Disease. Cell Death Dis. 2018, 9 (2). 10.1038/s41419-017-0221-2.

(38) Li, Y.; Liu, W.; Oo, T. F.; Wang, L.; Tang, Y.; Jackson-Lewis, V.; Zhou, C.; Geghman, K.; Bogdanov, M.; Przedborski, S.; Beal, M. F.; Burke, R. E.; Li, C. Mutant LRRK2R1441G BAC Transgenic Mice Recapitulate Cardinal Features of Parkinson’s Disease. Nat. Neurosci. 2009, 12 (7), 826–828. 10.1038/nn.2349.

(39) West, A. B.; Cowell, R. M.; Daher, J. P. L.; Moehle, M. S.; Hinkle, K. M.; Melrose, H. L.; Standaert, D. G.; Volpicelli-Daley, L. A. Differential LRRK2 Expression in the Cortex, Striatum, and Substantia Nigra in Transgenic and Nontransgenic Rodents. J. Comp. Neurol. 2014, 522 (11), 2465–2480. 10.1002/cne.23583.

(40) Lin, X.; Parisiadou, L.; Sgobio, C.; Liu, G.; Yu, J.; Sun, L.; Shim, H.; Gu, X. L.; Luo, J.; Long, C. X.; Ding, J.; Mateo, Y.; Sullivan, P. H.; Wu, L. G.; Goldstein, D. S.; Lovinger, D.; Cai, H. Conditional Expression of Parkinson’s Disease-Related Mutant Alpha-Synuclein in the Midbrain Dopaminergic Neurons Causes Progressive Neurodegeneration and Degradation of Transcription Factor Nuclear Receptor Related 1. J Neurosci 2012, 32 (27), 9248–9264. 10.1523/JNEUROSCI.1731-12.2012.

(41) Xiong, Y.; Neifert, S.; Karuppagounder, S. S.; Liu, Q.; Stankowski, J. N.; Lee, B. D.; Ko, H. S.; Lee, Y.; Grima, J. C.; Mao, X.; Jiang, H.; Kang, S. U.; Swing, D. A.; Iacovitti, L.; Tessarollo, L.; Dawson, T. M.; Dawson, V. L. Robust Kinase- and Age-Dependent Dopaminergic and Norepinephrine Neurodegeneration in LRRK2 G2019S Transgenic Mice. Proc. Natl. Acad. Sci. U. S. A. 2018, 115 (7), 1635–1640. 10.1073/pnas.1712648115.

(42) Steger, M.; Diez, F.; Dhekne, H. S.; Lis, P.; Nirujogi, R. S.; Karayel, O.; Tonelli, F.; Martinez, T. N.; Lorentzen, E.; Pfeffer, S. R.; Alessi, D. R.; Mann, M. Systematic Proteomic Analysis of LRRK2-Mediated Rab GTPase Phosphorylation Establishes a Connection to Ciliogenesis. Elife 2017, 6. 10.7554/eLife.31012.

(43) Steger, M.; Tonelli, F.; Ito, G.; Davies, P.; Trost, M.; Vetter, M.; Wachter, S.; Lorentzen, E.; Duddy, G.; Wilson, S.; Baptista, M. A.; Fiske, B. K.; Fell, M. J.; Morrow, J. A.; Reith, A. D.; Alessi, D. R.; Mann, M. Phosphoproteomics Reveals That Parkinson’s Disease Kinase LRRK2 Regulates a Subset of Rab GTPases. Elife 2016, 5. 10.7554/eLife.12813.

(44) Yamada, T.; Mcgeer, P. L.; Baimbridge, K. G.; Mcgeer, E. G. Relative Sparing in Parkinson’s Disease of Substantia Nigra Dopamine Neurons Containing Calbindin-D2sK; 1990; Vol. 526.

(45) Lam, S. S.; Martell, J. D.; Kamer, K. J.; Deerinck, T. J.; Ellisman, M. H.; Mootha, V. K.; Ting, A. Y. Directed Evolution of APEX2 for Electron Microscopy and Proximity Labeling. Nat. Methods 2014, 12 (1), 51–54. 10.1038/nmeth.3179.

(46) Dumrongprechachan, V.; Salisbury, R. B.; Butler, L.; Macdonald, M. L.; Kozorovitskiy, Y. Dynamic Proteomic and Phosphoproteomic Atlas of Corticostriatal Axons in Neurodevelopment. Elife 2022, 11. 10.7554/eLife.78847.

(47) Dumrongprechachan, V.; Soto, G.; Macdonald, M. L.; Kozorovitskiy, Y. Cell Type and Subcellular Compartment Specific APEX2 Proximity Labeling Proteomics in the Mouse Brain. 10.1101/2021.04.08.439091.

(48) Hobson, B. D.; Choi, S. J.; Mosharov, E. V; Soni, R. K.; Sulzer, D.; Sims, P. A. Subcellular Proteomics of Dopamine Neurons in the Mouse Brain. Elife 2022, 11. 10.7554/eLife.70921.

(49) Kamath, T.; Abdulraouf, A.; Burris, S. J.; Langlieb, J.; Gazestani, V.; Nadaf, N. M.; Balderrama, K.; Vanderburg, C.; Macosko, E. Z. Single-Cell Genomic Profiling of Human Dopamine Neurons Identifies a Population That Selectively Degenerates in Parkinson’s Disease. Nat. Neurosci. 2022, 25 (5), 588–595. 10.1038/s41593-022-01061-1.

(50) Fell, M. J.; Mirescu, C.; Basu, K.; Cheewatrakoolpong, B.; DeMong, D. E.; Ellis, J. M.; Hyde, L. A.; Lin, Y.; Markgraf, C. G.; Mei, H.; Miller, M.; Poulet, F. M.; Scott, J. D.; Smith, M. D.; Yin, Z.; Zhou, X.; Parker, E. M.; Kennedy, M. E.; Morrow, J. A. MLi-2, a Potent, Selective, and Centrally Active Compound for Exploring the Therapeutic Potential and Safety of LRRK2 Kinase Inhibition. Journal of Pharmacology and Experimental Therapeutics 2015, 355 (3), 397–409. 10.1124/jpet.115.227587.

(51) Kluss, J. H.; Mazza, M. C.; Li, Y.; Manzoni, C.; Lewis, P. A.; Cookson, M. R.; Mamais, A. Preclinical Modeling of Chronic Inhibition of the Parkinson’s Disease Associated Kinase LRRK2 Reveals Altered Function of the Endolysosomal System in Vivo. Mol. Neurodegener. 2021, 16 (1). 10.1186/S13024-021-00441-8.

(52) Chen, C.; Soto, G.; Dumrongprechachan, V.; Bannon, N.; Kang, S.; Kozorovitskiy, Y.; Parisiadou, L. Pathway-Specific Dysregulation of Striatal Excitatory Synapses by LRRK2 Mutations. Elife 2020, 9, 1–26. 10.7554/eLife.58997.

(53) He, H.; Ai, R.; Fang, E. F.; Palikaras, K. The Rab3 Family Proteins in Age-Related Neurodegeneration: Unraveling Molecular Pathways and Potential Therapeutic Targets. npj Aging. Nature Research December 1, 2025. 10.1038/s41514-025-00257-6.

(54) Wang, Y. F. H.; Okamoto, M.; Schmitz, K.; Südhof, T. Rim Is a Putative Rab3 Effector in Regulating Synaptic-Vesicle Fusion. Nature 1997.

(55) Dhekne, H. S.; Tonelli, F.; Yeshaw, W. M.; Chiang, C. Y.; Limouse, C.; Jaimon, E.; Purlyte, E.; Alessi, D. R.; Pfeffer, S. R. Genome-Wide Screen Reveals Rab12 GTPase as a Critical Activator of Parkinson’s Disease-Linked LRRK2 Kinase. Elife 2023, 12. 10.7554/eLife.87098.

(56) Schlüter, O. M.; Basu, J.; Südhof, T. C.; Rosenmund, C. Rab3 Superprimes Synaptic Vesicles for Release: Implications for Short-Term Synaptic Plasticity. Journal of Neuroscience 2006, 26 (4), 1239–1246. 10.1523/JNEUROSCI.3553-05.2006.

(57) Jenkins, M. L.; Harris, N. J.; Dalwadi, U.; Fleming, K. D.; Ziemianowicz, D. S.; Rafiei, A.; Martin, E. M.; Schriemer, D. C.; Yip, C. K.; Burke, J. E. The Substrate Specificity of the Human TRAPPII Complex’s Rab-Guanine Nucleotide Exchange Factor Activity. Commun. Biol. 2020, 3 (1). 10.1038/s42003-020-01459-2.

(58) Kerryn Berndsen; Lis, P.; Yeshaw, W. M.; Wawro, P. S.; Nirujogi, R. S.; Wightman, M.; Macartney, T.; Dorward, M.; Knebel, A.; Tonelli, F.; Pfeffer, S. R.; Alessi, D. R. PPM1H Phosphatase Counteracts LRRK2 Signaling by Selectively Dephosphorylating Rab Proteins. Elife 2019, 8:e50416.

(59) Vieweg, S.; Mulholland, K.; Bräuning, B.; Kachariya, N.; Lai, Y. C.; Toth, R.; Singh, P. K.; Volpi, I.; Sattler, M.; Groll, M.; Itzen, A.; Muqit, M. M. K. PINK1-Dependent Phosphorylation of Serine111 within the SF3 Motif of Rab GTPases Impairs Effector Interactions and LRRK2-Mediated Phosphorylation at Threonine72. Biochemical Journal 2020, 477 (9), 1651–1668. 10.1042/BCJ20190664.

(60) Phung, T. K.; Berndsen, K.; Shastry, R.; Phan, T. L. C. H. B.; Muqit, M. M. K.; Alessi, D. R.; Nirujogi, R. S. CURTAIN—A Unique Web-Based Tool for Exploration and Sharing of MS-Based Proteomics Data. Proc. Natl. Acad. Sci. U. S. A. 2024, 121 (7). 10.1073/pnas.2312676121.

(61) Liu, C.; Kaeser, P. S. Mechanisms and Regulation of Dopamine Release. Current Opinion in Neurobiology. Elsevier Ltd August 1, 2019, pp 46–53. 10.1016/j.conb.2019.01.001.

(62) Waschbüsch, D.; Purlyte, E.; Pal, P.; McGrath, E.; Alessi, D. R.; Khan, A. R. Structural Basis for Rab8a Recruitment of RILPL2 via LRRK2 Phosphorylation of Switch 2. Structure 2020, 28 (4), 406–417.e6. 10.1016/j.str.2020.01.005.

(63) Dhekne, H. S.; Yanatori, I.; Gomez, R. C.; Tonelli, F.; Diez, F.; Schü le, B.; Steger, M.; Alessi, D. R.; Pfeffer, S. R. A Pathway for Parkinson’s Disease LRRK2 Kinase to Block Primary Cilia and Sonic Hedgehog Signaling in the Brain. 2018. 10.7554/eLife.40202.001.

(64) Liu, C.; Kershberg, L.; Wang, J.; Schneeberger, S.; Kaeser, P. S. Dopamine Secretion Is Mediated by Sparse Active Zone-like Release Sites. Cell 2018. 10.1016/j.cell.2018.01.008.

(65) Pereira, D. B.; Schmitz, Y.; Mészros, J.; Merchant, P.; Hu, G.; Li, S.; Henke, A.; Lizardi-Ortiz, J. E.; Karpowicz, R. J.; Morgenstern, T. J.; Sonders, M. S.; Kanter, E.; Rodriguez, P. C.; Mosharov, E. V.; Sames, D.; Sulzer, D. Fluorescent False Neurotransmitter Reveals Functionally Silent Dopamine Vesicle Clusters in the Striatum. Nat. Neurosci. 2016, 19 (4), 578–586. 10.1038/nn.4252.

(66) Ducrot, C.; Bourque, M. J.; Delmas, C. V. L.; Racine, A. S.; Guadarrama Bello, D.; Delignat-Lavaud, B.; Domenic Lycas, M.; Fallon, A.; Michaud-Tardif, C.; Burke Nanni, S.; Herborg, F.; Gether, U.; Nanci, A.; Takahashi, H.; Parent, M.; Trudeau, L. E. Dopaminergic Neurons Establish a Distinctive Axonal Arbor with a Majority of Non-Synaptic Terminals. FASEB Journal 2021, 35 (8). 10.1096/fj.202100201RR.

(67) Banerjee, A.; Imig, C.; Balakrishnan, K.; Kershberg, L.; Lipstein, N.; Uronen, R. L.; Wang, J.; Cai, X.; Benseler, F.; Rhee, J. S.; Cooper, B. H.; Liu, C.; Wojcik, S. M.; Brose, N.; Kaeser, P. S. Molecular and Functional Architecture of Striatal Dopamine Release Sites. Neuron 2022, 110 (2), 248–265.e9. 10.1016/j.neuron.2021.10.028.

(68) Pylypenko, O.; Rak, A.; Reents, R.; Niculae, A.; Sidorovitch, V.; Cioaca, M.-D.; Bessolitsyna, E.; Thomä, N. H.; Waldmann, H.; Schlichting, I.; Goody, R. S.; Alexandrov, K. Structure of Rab Escort Protein-1 in Complex with Rab Geranylgeranyltransferase Transferase (RabGGTase). RabGGTase Is a Heterodimer Composed of Tightly Associated 68 KDa and 45 KDa Subunits and Belongs to the Family of Protein Prenyltrans-Ferases, Which Includes Farnesyl Transferase (FTase) and Geranylgeranyltransferase I (GGTaseI) (Casey and Sea; 2003; Vol. 11.

(69) Marshel, J. H.; Kim, Y. S.; Machado, T. A.; Quirin, S.; Benson, B.; Kadmon, J.; Raja, C.; Chibukhchyan, A.; Ramakrishnan, C.; Inoue, M.; Shane, J. C.; McKnight, D. J.; Yoshizawa, S.; Kato, H. E.; Ganguli, S.; Deisseroth, K. Cortical Layer-Specific Critical Dynamics Triggering Perception. Science (1979). 2019, 365 (6453). 10.1126/science.aaw5202.

(70) Zhuo, Y.; Luo, B.; Yi, X.; Dong, H.; Miao, X.; Wan, J.; Williams, J. T.; Campbell, M. G.; Cai, R.; Qian, T.; Li, F.; Weber, S. J.; Wang, L.; Li, B.; Wei, Y.; Li, G.; Wang, H.; Zheng, Y.; Zhao, Y.; Wolf, M. E.; Zhu, Y.; Watabe-Uchida, M.; Li, Y. Improved Green and Red GRAB Sensors for Monitoring Dopaminergic Activity in Vivo. Nat. Methods 2024, 21 (4), 680–691. 10.1038/s41592-023-02100-w.

(71) Howe, M. W.; Dombeck, D. A. Rapid Signalling in Distinct Dopaminergic Axons during Locomotion and Reward. Nature 2016, 535 (7613), 505–510. 10.1038/nature18942.

(72) Volta, M.; Beccano-Kelly, D. A.; Paschall, S. A.; Cataldi, S.; MacIsaac, S. E.; Kuhlmann, N.; Kadgien, C. A.; Tatarnikov, I.; Fox, J.; Khinda, J.; Mitchell, E.; Bergeron, S.; Melrose, H.; Farrer, M. J.; Milnerwood, A. J. Initial Elevations in Glutamate and Dopamine Neurotransmission Decline with Age, as Does Exploratory Behavior, in LRRK2 G2019S Knock-in Mice. Elife 2017, 6. 10.7554/eLife.28377.

(73) Wang, S. S. H.; Held, R. G.; Wong, M. Y.; Liu, C.; Karakhanyan, A.; Kaeser, P. S. Fusion Competent Synaptic Vesicles Persist upon Active Zone Disruption and Loss of Vesicle Docking. Neuron 2016, 91 (4), 777–791. 10.1016/j.neuron.2016.07.005.

(74) Kaeser, P. S.; Deng, L.; Wang, Y.; Dulubova, I.; Liu, X.; Rizo, J.; Südhof, T. C. RIM Proteins Tether Ca2+ Channels to Presynaptic Active Zones via a Direct PDZ-Domain Interaction. Cell 2011, 144 (2), 282–295. 10.1016/j.cell.2010.12.029.

(75) Leonard HL; Global Parkinson’s Genetics Program (GP2). Novel Parkinson’s Disease Genetic Risk Factors Within and Across European Populations. medRxiv. 2025 10.1101/2025.03.14.24319455.

(76) Liu G, Peng J, Liao Z, Locascio JJ, et al., Genome-wide survival study identifies a novel synaptic locus and polygenic score for cognitive progression in Parkinson’s disease. Nat Genet. 2021 53:787–793. doi: 10.1038/s41588-021-00847-6

(77) Schoch, S.; Castillo, P. E.; Jok, T.; Mukherjee, K.; Geppertk, M.; Wang, Y.; Schmitz, F.; Malenka, R. C.; Su, T. C. RIM1a Forms a Protein Scaffold for Regulating Neurotransmitter Release at the Active Zone; 2002. www.nature.com.

(78) Südhof, T. C. Neurotransmitter Release: The Last Millisecond in the Life of a Synaptic Vesicle. Neuron. 2013, pp 675–690. 10.1016/j.neuron.2013.10.022.

(79) Persoon, C. M.; Hoogstraaten, R. I.; Nassal, J. P.; van Weering, J. R. T.; Kaeser, P. S.; Toonen, R. F.; Verhage, M. The RAB3-RIM Pathway Is Essential for the Release of Neuromodulators. Neuron 2019, 0 (0), 1–16. 10.1016/j.neuron.2019.09.015.

(80) Dou, D.; Aiken, J.; Holzbaur, E. L. F. RAB3 Phosphorylation by Pathogenic LRRK2 Impairs Trafficking of Synaptic Vesicle Precursors. Journal of Cell Biology 2024, 223 (6). 10.1083/jcb.202307092.

(81) Skelton, P. D.; Tokars, V.; Parisiadou, L. LRRK2 at Striatal Synapses: Cell-Type Specificity and Mechanistic Insights. Cells 2022, 11 (1). 10.3390/CELLS11010169.

(82) Kuhlmann, N.; Milnerwood, A. J. A Critical LRRK at the Synapse? The Neurobiological Function and Pathophysiological Dysfunction of LRRK2. Frontiers in Molecular Neuroscience. Frontiers Media S.A. 2020. 10.3389/fnmol.2020.00153.

(83) Wong, Y. C.; Luk, K.; Purtell, K.; Burke Nanni, S.; Stoessl, A. J.; Trudeau, L. E.; Yue, Z.; Krainc, D.; Oertel, W.; Obeso, J. A.; Volpicelli-Daley, L. A. Neuronal Vulnerability in Parkinson Disease: Should the Focus Be on Axons and Synaptic Terminals? Movement Disorders. John Wiley and Sons Inc. 2019, pp 1406–1422. 10.1002/mds.27823.

(84) Parisiadou, L.; Yu, J.; Sgobio, C.; Xie, C.; Liu, G.; Sun, L.; Gu, X. L.; Lin, X.; Crowley, N. A.; Lovinger, D. M.; Cai, H. LRRK2 Regulates Synaptogenesis and Dopamine Receptor Activation through Modulation of PKA Activity. Nat Neurosci 2014, 17 (3), 367–376. 10.1038/nn.3636.

(85) Chen, C.; Masotti, M.; Shepard, N.; Promes, V.; Tombesi, G.; Arango, D.; Manzoni, C.; Greggio, E.; Hilfiker, S.; Kozorovitskiy, Y.; Parisiadou, L. LRRK2 Mediates Haloperidol-Induced Changes in Indirect Pathway Striatal Projection Neurons. Mol. Psychiatry 2025. 10.1038/s41380-025-03030-z.

(86) Matikainen-Ankney, B. A.; Kezunovic, N.; Mesias, R. E.; Tian, Y.; Williams, F. M.; Huntley, G. W.; Benson, D. L. Altered Development of Synapse Structure and Function in Striatum Caused by Parkinson’s Disease-Linked LRRK2-G2019S Mutation. J Neurosci 2016, 36 (27), 7128–7141. 10.1523/JNEUROSCI.3314-15.2016.

(87) Cheng HC, Ulane CM, Burke RE. Clinical progression in Parkinson disease and the neurobiology of axons. Ann Neurol. 2010 67(6):715–25. doi: 10.1002/ana.21995.

(88) Burke RE, O’Malley K. Axon degeneration in Parkinson’s disease. Exp Neurol. 2013 246:72–83. doi: 10.1016/j.expneurol.2012.01.011

(89) Nandhagopal, R.; Mak, E.; Schulzer, M.; McKenzie, J.; McCormick, S.; Sossi, V.; Ruth, T. J.; Strongosky, A.; Farrer, M. J.; Wszolek, Z. K.; Stoessl, A. J. Progression of Dopaminergic Dysfunction in a LRRK2 Kindred: A Multitracer PET Study. Neurology 2008, 71 (22), 1790–1795. 10.1212/01.wnl.0000335973.66333.58.

(90) Simuni, T.; Uribe, L.; Cho, H. R.; Caspell-Garcia, C.; Coffey, et al., Clinical and Dopamine Transporter Imaging Characteristics of Non-Manifest LRRK2 and GBA Mutation Carriers in the Parkinson’s Progression Markers Initiative (PPMI): A Cross-Sectional Study. Lancet Neurol. 2020, 19 (1), 71–80. 10.1016/S1474-4422(19)30319-9.

(91) Palanivel M, Ghosh KK, Mallam M, Bhuvanakantham R, Padmanabhan P, Lim KL, Gulyás B. Imaging advances to detect non-motor prodromal markers of Parkinson’s disease and explore therapeutic translation opportunities. NPJ Parkinsons Dis. 2025, 11(1):174. doi: 10.1038/s41531-025-01004-0.

(92) Janezic, S.; Threlfell, S.; Dodson, P. D.; Dowie, M. J.; Taylor, T. N.; Potgieter, D.; Parkkinen, L.; Senior, S. L.; Anwar, S.; Ryan, B.; Deltheil, T.; Kosillo, P.; Cioroch, M.; Wagner, K.; Ansorge, O.; Bannerman, D. M.; Bolam, J. P.; Magill, P. J.; Cragg, S. J.; Wade-Martins, R. Deficits in Dopaminergic Transmission Precede Neuron Loss and Dysfunction in a New Parkinson Model. Proc. Natl. Acad. Sci. U. S. A. 2013, 110 (42). 10.1073/pnas.1309143110.

(93) Vidyadhara, D. J.; Somayaji, M.; Wade, N.; Yücel, B.; Zhao, H.; Shashaank, N.; Ribaudo, J.; Gupta, J.; Lam, T. K. T.; Sames, D.; Greene, L. E.; Sulzer, D. L.; Chandra, S. S. Dopamine Transporter and Synaptic Vesicle Sorting Defects Underlie Auxilin-Associated Parkinson’s Disease. Cell Rep. 2023, 42 (3). 10.1016/j.celrep.2023.112231.

(94) Schindelin, J.; Arganda-Carreras, I.; Frise, E.; Kaynig, V.; Longair, M.; Pietzsch, T.; Preibisch, S.; Rueden, C.; Saalfeld, S.; Schmid, B.; Tinevez, J. Y.; White, D. J.; Hartenstein, V.; Eliceiri, K.; Tomancak, P.; Cardona, A. Fiji: An Open-Source Platform for Biological-Image Analysis. Nature Methods. 2012, pp 676–682. 10.1038/nmeth.2019.

(95) Bermejo, M. K.; Milenkovic, M.; Salahpour, A.; Ramsey, A. J. Preparation of Synaptic Plasma Membrane and Postsynaptic Density Proteins Using a Discontinuous Sucrose Gradient. Journal of Visualized Experiments 2014, No. 91. 10.3791/51896.

(96) Parisiadou, L.; Yu, J.; Sgobio, C.; Xie, C.; Liu, G.; Sun, L.; Gu, X. L.; Lin, X.; Crowley, N. A.; Lovinger, D. M.; Cai, H. LRRK2 Regulates Synaptogenesis and Dopamine Receptor Activation through Modulation of PKA Activity. Nat Neurosci 2014, 17 (3), 367–376. 10.1038/nn.3636.

(97) Bruderer, R.; Bernhardt, O. M.; Gandhi, T.; Xuan, Y.; Sondermann, J.; Schmidt, M.; Gomez-Varela, D.; Reiter, L. Optimization of Experimental Parameters in Data-Independent Mass Spectrometry Significantly Increases Depth and Reproducibility of Results. Molecular and Cellular Proteomics 2017, 16 (12), 2296–2309. 10.1074/mcp.RA117.000314.

(98) Tognetti, M.; Sklodowski, K.; Müller, S.; Kamber, D.; Muntel, J.; Bruderer, R.; Reiter, L. Biomarker Candidates for Tumors Identified from Deep-Profiled Plasma Stem Predominantly from the Low Abundant Area. J. Proteome Res. 2022, 21 (7), 1718–1735. 10.1021/acs.jproteome.2c00122.

(99) Ouyang, J. F.; Kamaraj, U. S.; Cao, E. Y.; Rackham, O. J. L. ShinyCell: Simple and Sharable Visualization of Single-Cell Gene Expression Data. Bioinformatics 2021, 37 (19), 3374–3376. 10.1093/bioinformatics/btab209.

